# Cell type-aware analysis of RNA-seq data

**DOI:** 10.1101/2020.07.13.201061

**Authors:** Chong Jin, Mengjie Chen, Danyu Lin, Wei Sun

**Affiliations:** Department of Biostatistics, University of North Carolina at Chapel Hill; Genetic Medicine, University of Chicago; Public Health Science Division, Fred Hutchinson Cancer Research Center; Department of Biostatistics, University of Washington

## Abstract

Most tissue samples are composed of different cell types. Differential expression analysis without accounting for cell type composition cannot separate the changes due to cell type composition or cell type-specific expression. We propose a computational framework to address these limitations: **C**ell Type **A**ware analysis of **R**NA-**seq** (CARseq). CARseq employs a negative binomial distribution that appropriately models the count data from RNA-seq experiments. Simulation studies show that CARseq has substantially higher power than a linear model-based approach and it also provides more accurate estimate of the rankings of differentially expressed genes. We have applied CARseq to compare gene expression of schizophrenia/autism subjects versus controls, and identified the cell types underlying the difference and similarities of these two neuron-developmental diseases. Our results are consistent with the results from differential expression analysis using single cell RNA-seq data.

## Background

RNA-seq data are often collected from bulk tissue samples, most of which comprise a heterogeneous population of different cell types. Several recent studies have demonstrated that studying cell type-specific gene expression and cell type composition is crucial for many scientific and clinical questions; for example, identifying genes and cell types related to virus infection [?] or tumor [?]. Most methods for differential expression (DE) studies [?,?,?] do not consider cell type compositions. A few exceptions include csSAM [?] and TOAST [?], which are designed for continuous gene expression data and do not fully utilize the count features of RNA-seq data. There are also a few methods with similar goals that were developed for DNA methylation data [?,?].

We develop a framework of cell type aware analysis of RNA-seq data (CARseq). We assume cell type compositions have been estimated by an existing method [?,?]. CARseq takes the input of bulk RNA-seq data and cell type fraction estimates and performs two tasks: comparison of cell type compositions and cell type-specific DE (CT-specific-DE). For CT-specific-DE, CARseq employs a negative binomial distribution to fully utilize the count features of RNA-seq data, which can substantially improve the statistical power. CARseq is a tribute to both the tradition that the gene expression of a mixture is the summation of non-negative expression of each cell type (i.e., deconvolution on a linear scale) [?], and that cell type-independent covariates are adjusted on a log scale. Our shrunken estimates of log fold change (LFC), currently unaddressed in other methods [?,?], produces a robust and interpretable quantification of CT-specific DE. We benchmark CARseq together with other methods under various simulation setups, illustrating CARseq can have substantially higher power while maintaining type I error control.

We apply CARseq to assess gene expression difference of schizophrenia (SCZ) or autism spectrum disorder (ASD) subjects versus healthy controls. SCZ and ASD are two severe neuropsychiatric disorders that are likely caused by disruption of brain development in early life (particularly in the prenatal and early postnatal period) due to environmental exposure combined with genetic predispositions [?]. The two diseases have shared vulnerability genes and overlapping symptoms [?]. For example, ASD is characterized by deficit social interaction and repetitive behaviors, which are similar to the negative symptoms (“negative” means taking away from normal state) of SCZ including social withdrawal and impaired motivation. There are also many differences, however, between the two diseases. For example, ASD is an early childhood disease (onset at 6 months to 3 years old) and most SCZ are diagnosed at young adulthood. Compared with ASD, SCZ has additional positive symptoms (“positive” means addition to the normal state) of delusions and hallucinations. The underlying biological mechanisms of the two diseases are not very well understood yet. Our results bring some insights on the molecular basis of these diseases. Specifically, we identified the relevant cell types underlying the difference and connection between the two diseases, for example, microglia for the connections of the two diseases through inflammation and oxidative stress [?]. We also reported an imbalance of excitation/inhibition neurons in SCZ but not ASD, which may explain the hallucination symptom in SCZ but not ASD [?].

Analyzing single cell RNA-seq (scRNA-seq) data is a promising solution for cell type-aware analysis. However, due to high cost and logistic difficulties (e.g., collection of high quality tissue samples, unbiased sampling of single cells), currently, it is very challenging, if not infeasible, to collect scRNA-seq data from large cohorts. In addition, scRNA-seq data may not capture the complete transcriptome. For example, 10x genomics platform measures gene expression at 5’ or 3’ of each transcript instead of the complete transcripts. If the massive amount of existing bulk RNA-seq data could be re-analyzed to study CT-specific expression and cell type composition, it could bring paradigm-shifting changes to many fields. Our work is one step towards this goal.

## Results

### Introduction to cell type-aware analysis

To assess the associations between cell type fractions and the covariate of interest, one needs to pay attention to the compositional nature of the data, e.g., we cannot modify the proportion of one cell type without altering the proportion of at least one other cell type [?]. Therefore, following a commonly used practice for compositional data analysis, we transform the *k* cell type fractions to *k* − 1 log ratios: log of the fraction of each of *k* − 1 cell types vs. a reference cell type.

The more challenging part is to assess CT-specific-DE, while we only observe the gene expression in bulk samples where the variability can come from both CT-specific expression and cell type fractions. Our model is built around the assumption that the expression in bulk samples is the summation of CT-specific expression weighted by cell fractions in linear scale (Figure 1). The model also allows the inclusion of cell type-independent covariates, such as age, gender, batch etc. We defer the details of our methods to the Methods Section.

**Figure 1:**
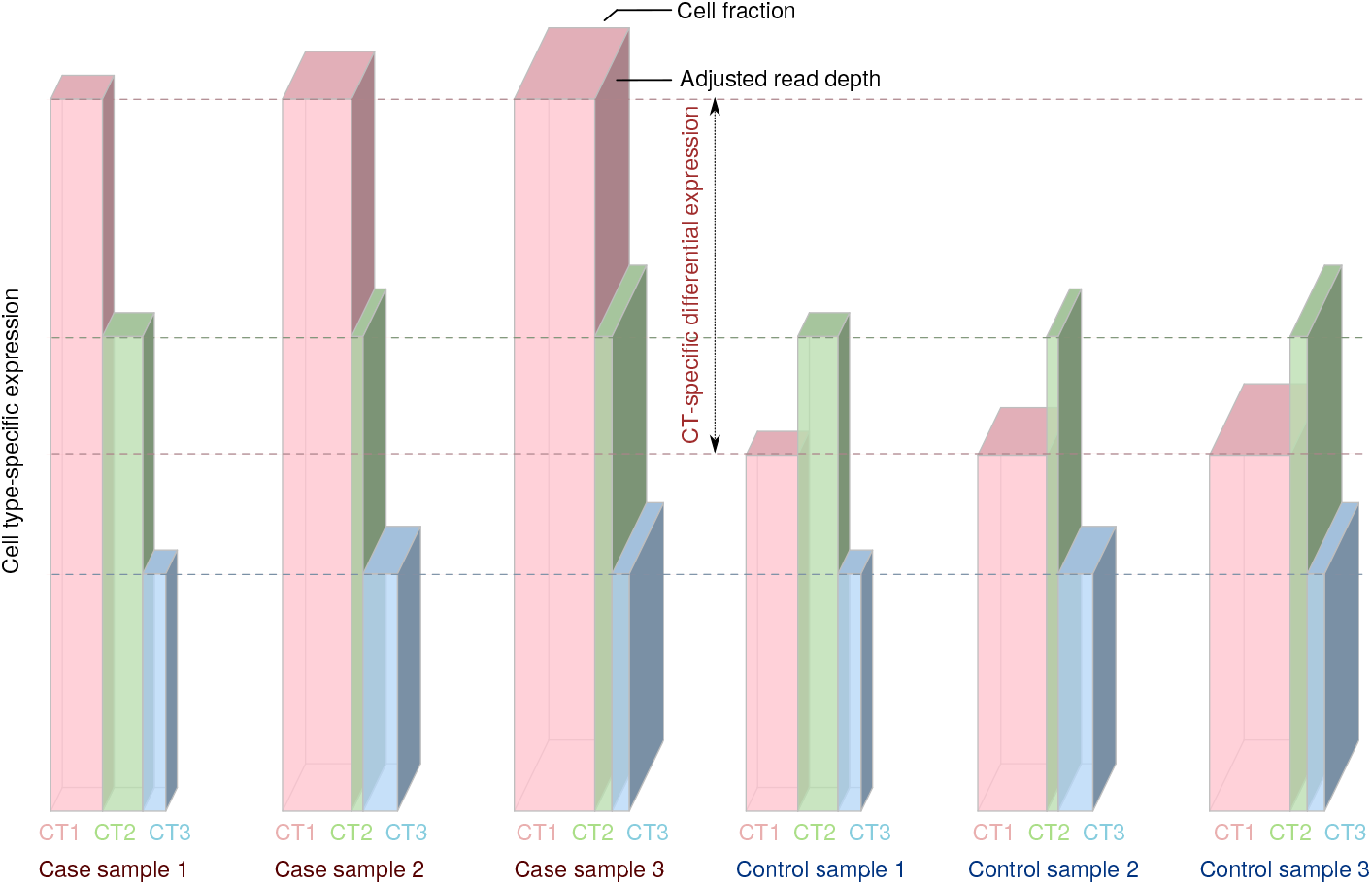
Illustration of CT-specific expression. Each grouped bar illustrates the total expression of a gene in a bulk sample (the volume of the grouped bar) that is the summation of gene expression from individual cell types (each bar for a cell type). The depth of each grouped bar is proportional to the covariate-adjusted read-depth. The width of each bar is proportional to its cell fraction, and the height of each bar is proportional to the CT-specific expression. The left/right three columns show three case/control samples, respectively. Our method estimates the mean value of CT-specific expression for case and control groups separately. In this toy example, cell type 1 (pink) has twice expression in cases than controls, while cell type 2 (green) and cell type 3 (blue) are not differentially expressed.

### Benchmarking methods through simulations

#### Simulation setup

We use a simulation study to evaluate the power and type I error of CT-specific-DE by CARseq, csSAM [?] and TOAST [?]. csSAM assesses CT-specific-DE by a two-step approach: estimation of CT-specific expression followed by testing using permutations. It has lower power than TOAST [?] and it cannot account for covariates. Both TOAST and CARseq combine the estimation of CT-specific expression and CT-specific-DE testing in one likelihood framework that allows adjustment for covariates. The difference is that CARseq adopts a negative binomial distribution that models gene expression decomposition on a linear scale. In contrast, TOAST uses a linear model that is less desirable to model count data. An alternative is to use TPM (transcripts per million) to replace count, which is a linear transformation of counts after adjusting for gene length and read-depth. We evaluated the performance of TOAST using both counts and TPM and observed similar results. Here we reported the results of TOAST using TPM and left the results using counts in Supplementary Figures 4-7.

We simulated CT-specific expression data that mirror the gene expression data from single nucleus RNA-seq (snRNA-seq) of human brains [?]. We simulated the cell fractions of 6 cell types to resemble our estimates (using ICeD-T [?] by default) from the Common Mind Consortium (CMC) bulk RNA-seq data [?] (Supplementary Figure 18). The major cell type, intended to imitate the excitatory neuron, taking the lion’s share of around 60% of the cells in each sample. The minor cell type and four other cell types, with much smaller fractions, were intended to represent inhibitory neurons and four non-neuron cell types, respectively. We also simulated a covariate in the mold of RNA integrity number (RIN) and specified its effect size based on estimates of RIN effect from the CMC data. We considered four series of simulation setups. In each series, one or two cell types are differentially expressed with different magnitudes. When two cell types are differentially expressed, their effect sizes could be on the same direction or opposite directions. More details of the simulation procedure can be found in Section B.1 of the Supplementary Materials.

#### CARseq has higher power and more accurate ranking

The conclusions from all the simulation setups are similar. Here we focus on the setup where the major cell type is differentially expressed (Figure 2) and presented the results where the minor cell type is differentially expressed or both the major and minor cell types are differentially expressed in Supplementary Figures 1-7). The simulation results demonstrated that CARseq is more powerful than TOAST, which is more powerful than csSAM, and correct specification of covariates can improve power and ensure the control of FDR [?] (Figure 2(A), Supplementary Figures 1-7). It worth noting the power of CT-specific-DE can be low when the sample size is small (e.g., *n* = 50, 25 cases vs. 25 controls), due to the uncertainty to estimate CT-specific expression. This is also the situation where CARseq shows much higher power than TOAST, with two to four folds of improvement (Figure 2(A)). The power gain of CARseq is even higher to uncover DE genes when the gene expression is low (Supplementary Figure 8), though overall most discoveries are from the genes with higher expression. Additional simulations show that CARseq is robust to noise/bias in cell type fraction estimates (Supplementary Materials B.1.5).

**Figure 2:**
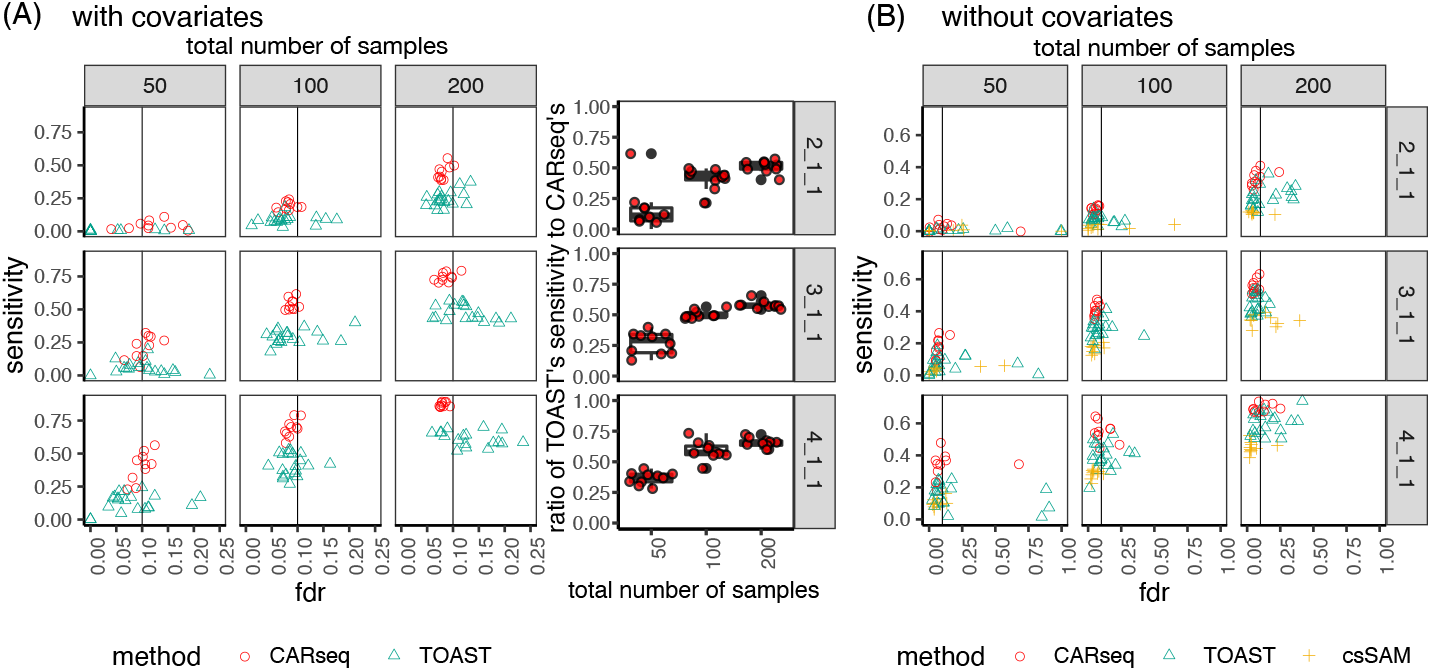
Simulation results. The FDR vs. sensitivity of several methods testing for CT-specific DE, when (a) the covariate is provided to the method, and (b) when the covariate is not provided to the method. The ratio of TOAST’s sensitivity to CARseq’s when the covariate is provided is illustrated in the boxplot. In the simulation, the cell type fractions are estimated using ICeD-T, a method that estimates cell fractions from expression of a bulk tissue sample using a reference of expression of purified cells. There are 10 simulation replicates for each combination of sample size (columns) as the total number of case-control samples and patterns of DE (rows). For each replicate, there are 2,000 genes following the pre-specified pattern of DE and 8,000 genes with no DE in any of the six cell types. In the notation for the DE pattern, the three numbers separated by underscores each represent the fold change in the major cell type, the minor cell type, and four other cell types. For example, 2_1_1 indicates that the major cell type is differentially expressed with fold change 2, the minor cell type and the other four cell type are not differentially expressed. The vertical line indicates the intended FDR level of 0.1. Note that csSAM does not support the inclusion of covariates; the scales of the x-axis in the two subfigures are different.

We further checked whether the ranking by the p-values produced by CARseq or TOAST is a meaningful indicator of DE genes using precision-recall curve.

CARseq has consistently higher AUC in all simulation setups (Supplementary Figure 9). For example, when there are 100 cases vs. 100 controls and the major cell type is differentially expressed with a fold change of 2, the precision-recall AUC is 0.57 and 0.30 for CARseq and TOAST. As a reference, when all the genes are ranked randomly, we expect a precision-recall AUC of 1/30 because one out of every 30 tests corresponds to a true DE relation.

#### CARseq delivers accurate estimates of effect sizes

CARseq quantify the effect size of CT-specific-DE by log fold change or shrunken log fold change (see Method Section for more details). TOAST defines the effect size as *β*/(*μ+β*/2), where *μ* is base-line expression in one group, and *β* is the gene expression difference between two groups. To make the results more comparable between CARseq and TOAST, we amend the effect size definition in TOAST and propose to define LFC as log(|*μ+β*|) −log(|*μ*|). To examine the reproducibility of effect size estimation, we divided the samples in a simulation replicate into two subsets of equal sizes and then compare the effect size estimates in the two subsets. It is clear that CARseq’s shrunken log fold change is best reproduced between the two subsets (Figure 3). For example, when sample size is 25 cases vs. 25 controls for each subset (middle panel of Figure 3), the percentage of genes whose direction of DE is correctly estimated in both two replicates are 64.1%, 76.9%, 29.3%, and 29.3% for effect size qualified by CARseq LFC, CARseq shrunken LFC, TOAST effect size, and TOAST LFC, respectively.

**Figure 3:**
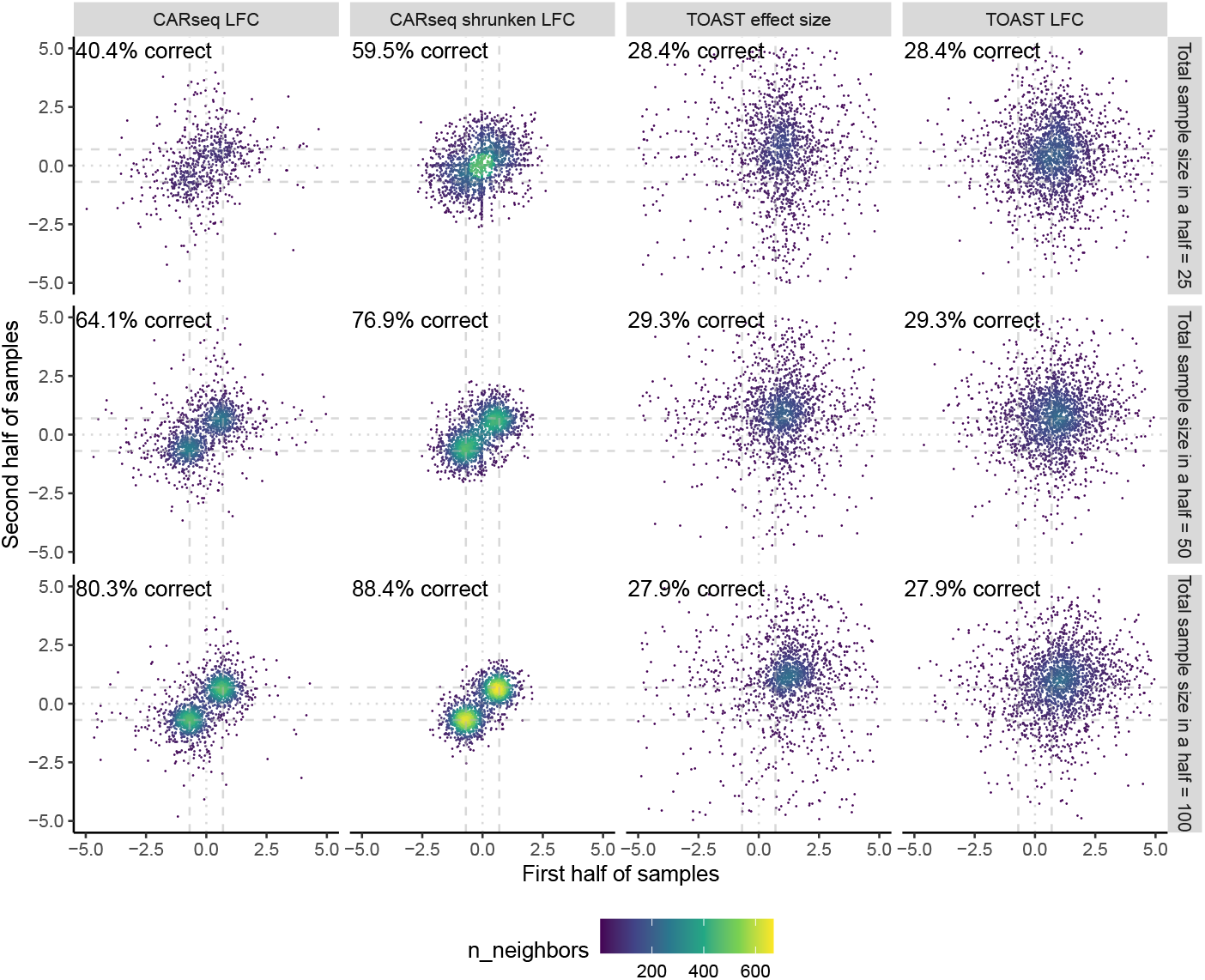
Reproducibility of effect size estimation. The reproducibility of effect size estimation among 2,000 differentially expressed genes in the major cell type (fold change of 2 or log fold change of 0.7) when CARseq and TOAST are applied to a simulation dataset of a mixture of three cell types where only the major cell type is differentially expressed. The percentage of genes whose direction of DE is correctly estimated in both two replicates is added at the top right corner of each plot.

### DE analysis between schizophrenia subjects and controls

We applied CARseq to study the gene expression of SCZ patients vs. controls using the bulk RNA-seq data of prefrontal cortex samples, generated by the CommonMind Consortium (CMC) [?], hereafter referred to as CMC-SCZ study. After filtering out the outlier samples reported by Fromer et al. [?], we had 250 SCZ subjects and 277 controls. In addition to the covariates used by Fromer et al. [?], we also included two surrogate variables to capture latent batch effects [?]. We found that the relative cellular abundance of the inhibitory neuron quantified by ICeD-T [?] is significantly higher in SCZ samples than control samples (*p*-value 1.5 ×10^−5^), and there is a similar trend for cell fraction estimates by CIBERSORT [?], though the difference is not significant (*p*-value 0.12). There is also a trend of relative depletion of oligodendrocyte, though it is not significant (*p*-values 0.32 for ICeD-T and 0.12 for CIBERSORT).

Gene set enrichment analysis (GSEA) recover that genes involved in unblocking or negative regulation of NMDA receptors are enriched among the DE genes in inhibitory neurons. The majority of the inhibitory-neuron-DE genes in NMDA pathways have lower expression levels in SCZ subjects than controls (Figure 4(C), Supplementary Figure 29), consistent with the hypofunction of NMDA. We found that the heat shock related genes are enriched in the DE genes in excitatory neurons, and they tend to have higher expression in SCZ subjects than controls (Figure 4(C), Supplementary Figure 29). This is consistent with previous findings that heat shock response plays a crucial role in the response of brain cells to prenatal environmental insults [?].

**Figure 4:**
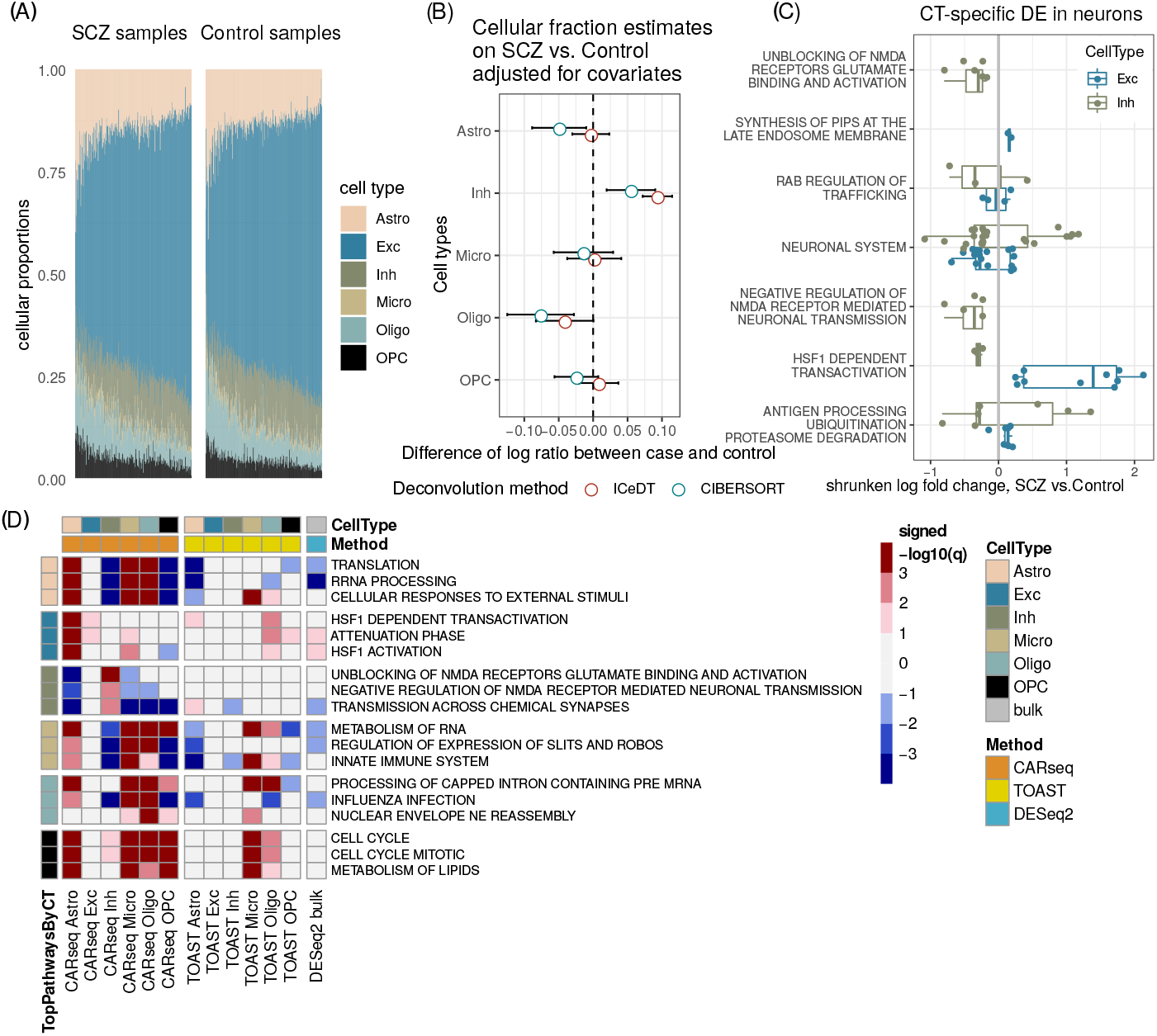
CT-specific DE results for SCZ vs. controls. CARseq on gene expression data between schizophrenia (SCZ) and controls. (A) Estimated cell fractions by ICeD-T sorted by increasing fractions of excitatory neurons. (B) The effect size of case-control status on relative cell fractions against excitatory neurons (log ratio of the cell type of interest vs. excitatory neuron). The standard errors are denoted by bars. The confidence interval are calculated based on the linear model estimates for the case-control status effect, conditioning on other covariates. (C) CT-specific DE in excitatory neurons and inhibitory neurons in significantly enriched pathways. Only genes with a *p*-value less than 0.05 are shown. The color scheme is consistent with the color used in panel (A) and (D) for excitatory neurons and inhibitory neurons. (D) Gene set enrichment analysis (GSEA) results on REACTOME pathways. Three top pathways were shown for each cell type, ranked by −log10 q value with the sign of normalized enrichment score (NES). Positive NES indicates enrichment of genes with small p-values. The p-value of GSEA were obtained from `“fgseaMultilevel”` function in `“fgsea”` R package. We further converted the p-values to *q*-values using `“get_qvalues_one_inflated”` in our CARseq package. More details of GSEA are presented in the subsection of “Gene set enrichment analysis” in Method section.

### DE analysis between ASD subjects and controls

We analyzed the bulk RNA-seq data from ASD subjects and controls, published by a UCLA group [?,?], hereafter referred to as UCLA-ASD study. They reported findings on 251 post-mortem samples of frontal and temporal cortex and cerebellum for 48 ASD subjects versus 49 controls and found significantly differentially expressed genes in cortex but not in cerebellum [?]. In this study, we focus on frontal cortex region based on positive findings of DE genes in this earlier study and that it matches the brain region of SCZ data analyzed in this paper. After filtering by brain regions, we ended up with 42 ASD subjects and 43 control subjects (See Method section for details).

We performed similar CARseq analysis as for the CMC-SCZ data and presented the results in Supplementary Materials Section B.3. Here we briefly summarize the results. First, we found the relative abundance of astrocyte and microglia are higher in ASD subjects than controls, consistent with the previous studies that suggest neuroinflammation plays an important role in the etiology of ASD [?]. Second, CARseq recovered much more DE genes than TOAST and the known ASD risk genes [?] are significantly enriched among the DE genes found by CARseq. Third, gene set enrichment analysis suggest the relevance of AMPA activity in the pathophysiology of ASD.

Next we compare the results from CARseq and TOAST versus the cell type-specific DE results from a single nucleus RNA-seq (snRNA-seq) dataset [?]. This dataset includes snRNA-seq data of 62,166 nuclei from the prefrontal cortex of 13 ASD cases vs. 10 healthy controls. We applied DESeq2 to assess cell type-specific DE between ASD and controls using the cell type-specific pseudo-bulk RNA-seq data, which is constructed by adding up the read counts across all the cells of the same cell type for each gene and each individual. Based on the p-value distribution for each cell type (whether there is enrichment of small p-values), we conclude that using this snRNA-seq data we can detect some DE signals in three cell types, Astro, Exc, and Inh, very limited DE signals in Oligo and OPC, and no DE signal in Microglia (Supplementary Figure 54). When comparing with our CARseq results, we found significant overlap in Astro, Exc, and Inh. Lack of overlap for Microglia, Oligo, and OPC could be partly due to the lack of DE signals in these cell types in snRNA-seq data. In contrast, there is no significant overlap for the results of TOAST or DESeq2 (Supplementary Figure 55).

### Concordant microglia-specific DE genes between SCZ and ASD

We found an interesting pattern that genome-wide microglia-specific-DE p-values show significant correlations between SCZ and ASD (Figure 5(A), Pearson and Spearman correlation are 0.14 and 0.23, respectively, and p-value < 2 × 10^−16^ for either correlation). In addition, the fold changes of microglia DE genes in different pathways also show consistent patterns between SCZ and ASD (Supplementary Figure 30(A) vs. 42(A)): up-regulation in innate immune system and cell cycle, and down-regulation in translation, slit-robo signaling pathway, and influenza infection. We further study the overlapping DE genes. Using a liberal p-value cutoff of 0.05, we identified 1,674 and 355 microglia-specific-DE genes in SCZ and ASD studies, respectively, with an overlap of 65 genes. This overlap is significantly larger than 33 overlaps expected by chance (p-value 9.6 × 10^−9^ by Chi-squared test). Several REACTOME pathways are over-represented by these 65 genes (by R package goseq, Figure 5(B), Supplementary Table 6). One interesting finding is “Selenoamino acid metabolism”. Since selenium-dependent enzymes prevent and reverse oxidative damage in brain, our findings support that selenium-dependent enzymes could mediate the relation between antioxidants and SCZ/ASD [?,?].

**Figure 5:**
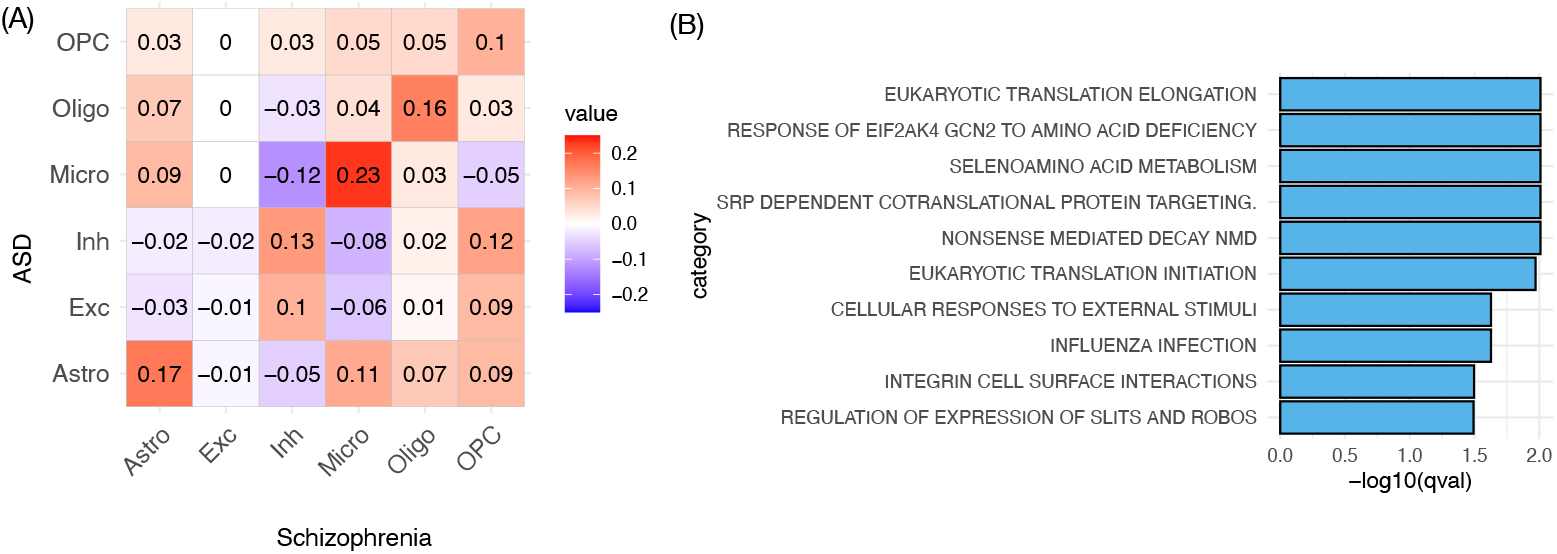
Microglia-specific-DE signals. (A) The Spearman correlation matrix of −log10(Microglia-specific-DE p-values by CARseq) (calculated by CARseq) between CMC-SCZ and UCLA-ASD studies. (B) The REACTOME pathways that are over-represented by the 65 genes with microglia-specific-DE p-values smaller than 0.05 in both CMC-SCZ and UCLA-ASD studies.

### Comparing DE testing by CARseq versus DESeq2

DESeq2 [?] is a good representative of existing methods for DE analysis of bulk tissue samples. In our default analysis, DESeq2 does not take any cell type composition as covariates. After accounting for cell type compositions (by including log ratios of cell type compositions), DESeq2 identified more DE genes using the CMC-SCZ data (from 1,009 to 1,888, with an intersection of 810 at q-value 0.1, Supplementary Table 1), but much less DE genes in UCLA-ASD study (from 1063 to 481, with an intersection of 185 at q-value 0.1, Supplementary Table 1). If a gene is associated with the case/control status before accounting for cell type compositions, but not so afterwards, its association with the case/control status is likely due to confounding effect of cell type composition. On the other hand, if a gene is not associated with the case/control status before accounting for cell type compositions, but the association becomes significant afterwards, it is likely because cell type composition explains part of the within group variance and thus increase the power of DE testing. In either case, as expected, the expression of these genes are more likely to be associated with cell type compositions (Supplementary Figures 48-49). In both SCZ and ASD analyses, most findings from DESeq2 were not identified as CT-specific-DE genes by CARseq (Supplementary Figure 28 and 42). There can be multiple reasons. First, some DESeq2 DE genes may be associated with case/control status because of confounding with cell type composition. Second, those genes likely have CT-specific-DE patterns in a substantial proportion of cells (e.g., in the most abundant cell type or several cell types), and thus such DE patterns can be captured by bulk data. In contrast, CARseq may have lower power to detect such genes because CARseq needs to pay for the price of “CT-specific expression estimation uncertainty” that is incorporated in its likelihood framework. In addition, CARseq may have limited power if the proportion of the relevant cell type has small variance across individuals.

## Discussion

A practical consideration of using CARseq is that it may have limited power when the sample size is small. This is the price that we have to pay for the uncertainty of estimating CT-specific expression from bulk RNA-seq data. As a rule of thumb, we do not recommend using CARseq when the sample size minus the number of covariates is smaller than 20. For large studies, e.g., with hundreds of samples, it may worth considering a study design to generate scRNA-seq data in a subset of samples, and generate bulk RNA-seq data from all the samples. The scRNA-seq data can be used to generate cell type-specific gene expression reference for cell type fraction estimation, which can be used for the CARseq analysis on bulk RNA-seq data. In addition, the scRNA-seq data can also be used to validate the results of CARseq. CARseq also requires the estimates of cell type fractions, which relies on reference of cell type-specific gene expression data. We expect with the development of human cell atlas [?], such resource in other tissues will be generated in the near future, and thus enable CARseq analysis in broader tissues and relevant diseases.

Although CARseq has higher higher power than TOAST, CARseq is computationally much more demanding than TOAST. For example, using 32 threads on a compute node of E5-2680 v3 CPUs, CARseq took about 21 minutes to run the real data analysis for 250 Schizophrenia patients vs. 277 Control with 20,788 genes while TOAST took less one minute.

We have applied both CARseq and TOST to perform cell type aware analysis of postmortem gene expression data from SCZ and ASD. The molecular mechanisms underlying SCZ and ASD can be divided into two categories: alterations in neurotransmitter systems and stress-associated signaling including immune/inflammatory-related processes and oxidative stress [?]. NMDA and AMPA are two types of receptors for neurotransmitter glutamate. We found evidence for hypofunction of NMDA in SCZ (particularly in inhibitory neurons) and dysregulation of AMPA in ASD. While excitation-inhibition (E-I) imbalance has been suggested as a common feature of SCZ and ASD, we found the E-I imbalance in SCZ but not in ASD. This is consistent with previous finding that the hypofunction of NMDA could cause E-I imbalance [?] and that E-I imbalance is the underlying mechanism for hallucination [?], which is a symptom of SCZ but not ASD. Thus our finding of E-I imbalance in SCZ but not in ASD may explain part of the symptom difference between the two diseases. A recent study also found no E-I difference between ASD and controls [?].

Prenatal stress referred as maternal immune activation (regardless the cause such as infection by different pathogens or immune stimulation) can lead to SCZ or ASD [?], implying the role of immune system in disease pathology. Microglia is the tissue resident macrophages in brain and plays a central role in immune response in brain. We found microglia-specific DE genes have significant overlap between SCZ and ASD and they have higher expression in SCZ/ASD subjects than in controls, suggesting microglia are in more active states in SCZ/ASD than controls, CARseq can certainly be applied to study other diseases or traits. As an example, we illustrate its usage to study melanoma cancer in Supplementary Materials Section B.7.

## Acknowledgements

R01GM105785 W.S. and C.J., R21CA224026 W.S., R01GM126550 W.S., R01HG009974 D.L., P01CA142538 D.L., R01GM126553 M.C., NSF 2016307 M.C., and a Sloan Foundation Fellowship M.C. We also appreciate helpful discussions with Dr. Paul Little.

## Author Contributions

W.S. and C.J. conceived the approach. C.J. implemented the methods and performed analysis, with inputs from W.S., M.C., and D.L. W.S. and C.J. wrote the paper, with inputs from M.C. and D.L.

## Competing Interests statement

None declared.

## Methods

### Likelihood function of CARseq model

Let *T_ji_* be the RNA-seq read count (or fragment count for paired-end reads) for gene *j* ∈ {1,…, *J*}and sample *i* ∈ {1,…, *n*}, where *J* is the total number of genes and *n* is the number of bulk samples. We denote the cell fraction for cell type *h* ∈ {1,…, *H*}in the *i*-th sample by 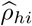.

We assume *T_ji_* follows a negative binomial distribution: *T_ji_* ~ *f_NB_*(*p_ji_, ϕ_j_*), with mean value *μ_ji_* and dispersion parameter *ϕ_j_*. Since deconvolution on a linear (non-log) scale yields better accuracy [?], we let:

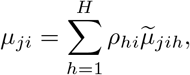

where 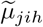 is the mean expression of the *j*-th gene in the *h*-th cell type of the *i*-th sample. The above deconvolution states that the expected total read count is the summation of expected CT-specific read count weighted by cell fractions across all cell types *h* ∈{1,…, *H*}. In practice, cell fraction estimates 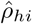 are used in place of *ρ_hi_*.

We model the relation between 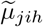 and *M*_0_ CT-specific covariates through a log link function, which is commonly used for negative binomial regression: 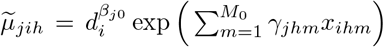, where *d_i_* is the sequencing read depth of sample *i*, *γ_jhm_* and *x_ihm_* are the regression coefficient observed data for the *m*-th covariate. In all the analyses of this paper, we use 75 percentile of the expression across all the genes within a sample.

The effect sizes of many covariates may not vary across cell types. For example, since RNA integrity number (RIN) quantifies sample RNA quality, it would associate with observed gene expression in the same way regardless of the original cell type. By separating cell type-independent covariates from CT-specific covariates, we can construct a model with less degrees of freedom. Suppose *M* out of *M*_0_ parameters are CT-specific and the rest *K* = *M*_0_ − *M* parameters are cell type-independent, we have:

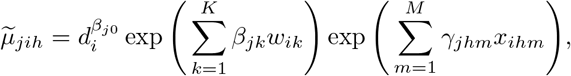

where *w_ik_* is the value of the *k*-th cell type-independent covariate in sample *i*.

The log-likelihood can be maximized using iteratively weighted least squares (IWLS) with some tweaks (Supplementary Materials Section A). After that, we can construct likelihood ratio statistics to conduct the CT-specific-DE tests. Although the likelihood-based testing framework can be generalized to accommodate a variety of tasks [?], e.g., test for a continuous variable or test for a linear combination of regression coefficients, our main focus in the article is CT-specific-DE tests among two or more groups.

We noted that CARseq reports large LFC estimates in some cell types, which probably reflect estimation uncertainty, particularly for the cell types with low proportion or when the sample size is small. To mitigate this problem, we developed a shrunken LFC estimation procedure, see Section A.5 in Supplementary Materials for details.

### Comparison with other methods

There are a few alternatives to our methods, though they were indeed developed for different purposes and not best suited for RNA-seq data analysis. TOAST [?] is a method for CT-specific DE or differential methylation analysis. It uses a linear model that is more flexible than our negative binomial model to handle different types of data, though for RNA-seq count data with a very strong mean-variance relationship, a linear model that assumes homogeneous variance has to choose between variance stabilization (e.g., by log-transformation of gene expression) or deconvolution in linear scale. See more discussions of our method and TOAST in Section A.6.1 in Supplementary Materials. All the analysis performed in real data has been done using both CARseq and TOAST and additional results for TOAST are available in Supplementary Figures.

Accounting for observed/unobserved confounding covariates is crucial for DE analysis, and the unobserved covariates can be estimated by surrogate variable analysis (SVA) [?]. In simulation data, we found that not accounting for relevant covariates can lead to inflated type I error. This limits the application of csSAM that cannot adjust for covariates in a lot of practical settings. For this reason, we did not apply csSAM in real data analysis.

### Details of simulations

We generated simulate data in three steps. First simulated cell fractions using a Dirichlet distribution with parameters estimated from cell fractions estimated from CMC-SCZ data followed by cell size correction. Next simulated CT-specific gene expression and finally simulated bulk gene expression. Finally we used the simulated bulk and CT-specific data to estimate cell-type fractions by ICDeT and use the estimated cell type fractions, instead of simulated cell type fractions for our analysis. We benchmarked CARseq against csSAM 1.2.4, TOAST 0.99.8, and DESeq2 1.24.0. Next we provide more details of simulations.

We generated read count data of bulk tissue as a mixture of six cell types, with sample sizes 50, 100. and 200, which was further divided equally into case and control groups. The total number of genes was 10,000, among which 2,000 genes had spiked-in cell type-specific differential expression between groups.

The read counts in mixture samples were generated from a negative binomial distribution:

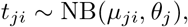

with the mean structure being

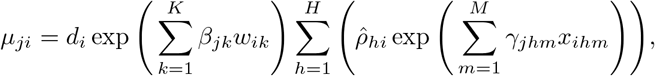

where *H* = 6 is the number of cell types, *M* = 2 corresponds to cell type-specific effects (case/control groups), *K* = 1 is cell type-independent batch effect (RIN), and 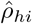 are cell type fractions simulated from a Dirichlet distribution with parameters (12.09, 53.01, 8.39,1.88, 5.54, 3.75).

The number of cell types *H* was 6, reflecting the reference matrix with 6 cell types constructed from MTG single cell data. This was achieved by combining more refined cell type definitions into six categories: excitatory neurons (Exc) and inhibitory neurons (Inh), astrocytes (Astro), microglia (Micro), oligodendrocyte (Oligo), Oligodendrocyte Progenitor Cell (OPC).

To set realistic parameters in data simulation, we used CARseq to fit a model of read counts with CMC data. For every gene, we fitted a baseline model without cell types composition effects: *t_ji_* ~ NB(*μ_ji_*, *θ_j_*) with the mean structure *μ_ji_*= *d_i_* exp(*β_j1_w_i1_*)*γ_j_*, where *d_i_* is sample-level read depth, *w*_*i*1_ is the RIN per sample. Both *d_i_* and *w*_*i*1_ are known when fitting the model using CMC data. We fitted a negative-binomial model using CARseq to obtain estimates of the triple (*β_ji_, γ_j_*, *θ_j_*), where *θ_j_* is the over-dispersion parameter.

In the scatterplot of the triple (*β*_*j*1_, *γ_j_*, *θ_j_*) across 20,614 highly expressed genes in the log scale (no log transformation is needed for *β*_*j*1_, which is already parametrized in the log scale), we noticed that multivariate Gaussian distribution is a good approximation after we remove 31 genes with a much higher overdispersion than the others. We compute and base our simulation on the estimated covariance matrix of expression, batch effect, and overdispersion.

There is a large proportion of zeros in estimated cell type-specific expression because it is hard to accurately estimate cell type-specific expression when the variance of cell fractions was low. Therefore using estimates of CT-specific gene expression for simulation is challenging. To mitigate the problem, when generating the reference matrix, we assumed that the expected expression of every cell type *γ_jh_* followed a log-normal distribution with the same mean and variance across cell types, and the mean and variance of mixture expression *γ_j_* can be used as a substitute. However, we needed to add correlation between cell types to make the expression pattern to be more realistic, as in a large number of genes, the gene expression would be similar across cell types, the most prominent examples being the housekeeping genes. We also noticed that gene expression and gene lengths are positively correlated. Gene lengths were needed to calculate TPM to generate the input data of csSAM.

To estimate the correlation between cell type-specific gene expression *γ_jh_*, and the correlation between gene expression *γ_jh_* and gene lengths *ℓ_j_*, we used MTG single cell data to create an average cell expression for 6 cell types, and estimated the correlation structure between cell type-specific expression *γ_jh_*, *h* ∈{1,…, 6}, and gene lengths *ℓ_j_*, all in log scale. We then took the median of all pairwise correlations of gene expression across cell types (or the median of all correlations between gene expression and gene length) and fill it into the correlation matrix to simulate data,. We assumed the correlation was zero between gene lengths and batch effect or overdispersion. This completed the specification of covariance matrix of three cell types, batch effect, overdispersion and, gene lengths. Thus we generated the tuple (*γ*_*j*1_, *γ*_*j*2_, *γ*_*j*3_, *β*_*j*1_, *θ_j_*, *ℓ_j_*) under log scale except for *β*_*j*1_ using a multivariate Gaussian distribution:

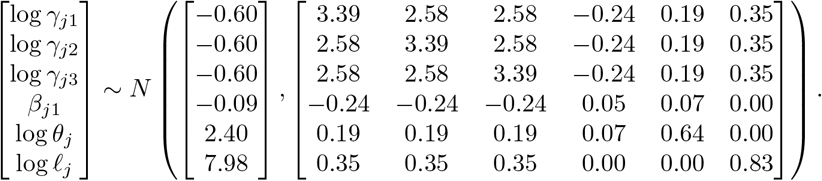

The batch effect variable, RIN wi1, was generated using a normal distribution with mean and variance estimated from variables collected in CMC data: log *w*_*i*1_ ~ *N*(7.61, 0.89).

We generated read depths di using log-normal distribution with mean and variance estimated from mixture expression of CMC data: log *d_i_* ~ *N*(6.70,0.27).

When estimating cell type fractions, we need a signature matrix. To select the signature genes, for each cell type, we collected the top 100 genes with the largest fold change of its gene expression compared to gene expression from all other cell types in the non-log scale. We also incorporated the batch effect from RIN to construct adjusted reference matrix for ICeD-T.

We specified the pattern of differential expression we would like to spike in as c = (*c*_1_, *c*_2_, *c*_3_) for major, minor and all other cell types. Among 10,000 genes, suppose 8,000 genes were not differentially expressed between case and control samples, 1,000 genes had higher expression in cases than controls, and 1,000 genes had higher expression in controls than cases. Then we calculated *μ_ji_* and generated the observed mixture read count matrix with each entry *t_ji_* following a negative binomial distribution. Using the same approach, we implemented more complicated patterns of differential expression where more cell types were differentially expressed. The test we implemented in the simulation studies has the null hypothesis *γ*_*jh*1_ = *γ*_*jh*2_ for any gene *j* and cell type *h*, which says the cell type-specific mean are the same across two groups when controlled for other covariates.

### Estimation of cell type compositions

We use two reference-based methods to estimate cell type compositions of bulk tissue samples: CIBERSORT [?] and ICeD-T [?]. CIBERSORT is a popular method that use a support vector regression to estimate cell type proportions. ICeD-T is a likelihood-based method that model gene expression using log-normal distribution. It allows a subset of genes to be “aberrant” in the sense that the CT-specific gene expression of such genes are inconsistent between bulk tissue samples and external reference. Such aberrant genes are down-weighted in estimating cell type proportions. Unless specifically noted, the cell type composition of bulk tissue samples are estimated using ICeD-T by default.

We generated CT-specific gene expression reference using snRNA-seq data from the middle temporal gyrus (MTG) of the human brain [?]. This is not a perfect match for the bulk RNA-seq data that are from pre-frontal cortex (PFC). We have compared MTG with another snRNA-seq data generated from human PFC as well as other brain regions using DroNC technique. The CT-specific gene expression is similar between the two datasets, except for endothelial cells. We chose to use MTG data to generate reference since it has much higher depth and better coverage, making it more similar to bulk RNA-seq data. We exclude endothelial in our analysis since there are only 8 endothelial cells in MTG data and its expression has very weak similarity to the endothelial cells from DroNC data. See Supplementary Materials Section B.2.1 for more details.

A related question is that when estimating cell type fractions, an implicit assumption is that the signature genes’ cell type-specific expression level does not change in different conditions. Then it seems to be a contradiction when we assess its DE. A more rigorous approach is to assess DE only for non-signature genes. However, since cell fractions are estimated using hundreds of genes with robust models, removing any one signature gene will not lead to a noticeable change of cell type fraction estimates. Therefore, it is as if we assess DE of a signature gene without using it as part of the signature matrix. On the other hand, if too many genes within the signature gene set are detected to be differentially expressed, the accuracy of the cell type fraction estimates is questionable, and an alternative signature gene set should be selected.

### CARseq analysis for SCZ

The gene expression data and sample characteristics data were downloaded from CommonMind Consortium (CMC) Knowledge Portal (see section URLs). We include the following covariates in our CARseq CT-specific DE analysis:

- log transformed read-depth (75 percentile of gene expression across all the genes within a sample),
- institution (a factor of three levels for the three institutes where the samples were collected),
- age, gender, and PMI (Post-mortem interval),
- RIN (RNA integrity number) and its square transformation RIN^2^,
- a batch variable “Libclust”, which is clusters of library batches into 8 groups,
- two genotype PCs, and two surrogate variables.

The surrogate variables were calculated after accounting for cell type compositions. Specifically, we add the log ratios of cell type compositions (with excitatory neuron as baseline) as the covariates and then calculate surrogate variables using R function sva from R package sva [?].

The covariates selected in our model are mostly similar to those included in the original analysis [?] except two differences. One is that we included two instead of five genotype PCs in our analysis since other PCs are not associated with gene expression data (Supplementary Figure 1). Surrogate variables were computing using the R package “sva” [?]. Two surrogate variables are included because adding these two surrogate variables increased the variance explained (*R*^2^) in a linear model to fit log-scale mixture expression from 0.55 to 0.68, while more surrogate variables offered a comparably limited increase in *R*^2^. Prior to inclusion in the model, all the continuous covariates were scaled to ensure numerical stability [?].

### CARseq analysis for ASD

The gene expression data (expected read counts derived from RSEM) were downloaded from Freeze1 of PsychENCODE Consortium (PEC) Capstone Collection, and the accompanying meta data and clinical data were downloaded from PsychENCODE Knowledge Portal, see Section URLs for the exact links. There are 341 samples from 100 individuals. We kept the samples from BrodmannArea 9 (BA9), including 89 samples from 85 individuals. Four individuals have duplicated samples and we chose the one with higher RIN. These 85 individuals include 42 ASD subjects and 43 controls, and they were from two brain banks: 53 from Autism Tissue Program (ATP) and 32 from NICHD, see Parikshak et al. [?] for more details of this dataset.

We examined the association between each potential covariates and genomewide gene expression and found PMI and Sex are not associated with gene expression, as evidenced by a uniform distribution of p-values, therefore we removed these two covariates and used the following covariates in our analysis.

- log transformed read-depth (75 percentile of gene expression across all the genes within a sample),
- BrainBank (a factor of two levels),
- SequencingBatch (a factor of 3 levels),
- age, RIN (RNA integrity number),
- four sequencing surrogate variables (SeqSVs).

The SeqSVs, which are the notations used by Parikshak et al. [?] are PCs derived from sequencing QC metrics. We used 4 principal components because they explained 99% of the variance of the sequencing metrics. Prior to inclusion in the model, all the continuous covariates were scaled to ensure numerical stability [?].

### Gene set enrichment analysis

The gene set enrichment was done on REACTOME pathways downloaded from https://www.gsea-msigdb.org/gsea/msigdb/download_file.jsp?filePath=/msigdb/release/7.1/c2.cp.reactome.v7.1.symbols.gmt. There are originally 1,532 pathways, of which 1,090 pathways have a size between 10 and 1,000 genes.

For each cell type, we used `“fgseaMultilevel”` function in `“fgsea”` R package to simultaneously calculate p-values and normalized enrichment scores (NES) across the 1,090 pathways without any weights (fgseaMultilevel argument gseaParam = 0) in a gene list ranked by potentially CT-specific *p*-values from the DE analysis. The p-values across the 1,090 pathways were then converted to *q*-values using `“get_qvalues_one_inflated”` in our CARseq package. Next, we collected in a table all the candidates of the pathway-cell type pairs satisfying NES > 0 (genes in the pathway tend to have smaller *p*-values), and sorted them by the rank of increasing *q*-values and decreasing NES within each cell type. We then deduplicated the pathways by only retaining the first appearance of each pathway in the table. The top *N* pathway-cell type pairs were subsequently chosen. For illustrative purposes, *N* was picked to be 3 in our paper.

In the Main Figures, the primary DE method was CARseq, and the top pathways were defined by GSEA results from genes ranked by CARseq CT-specific-DE *p*-values. In the Supplementary Figures, we also reported heatmaps featuring top pathways defined by GSEA results based on rankings by TOAST CT-specific-DE *p*-values.

### FDR control procedure

We use *q*-value to control FDR [?]. The calculation of *q*-value requires an estimate of the overall proportion of null *p*-values 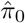. We use the following formula that specifically accommodates the situation where a proportion of p-values equal to 1, implemented in function `get_qvalues_one_inflated of R package CARseq`:

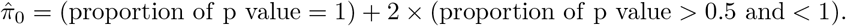

### Data availability

The data used in this study are available in the following public repositories. snRNA-seq data for CT-specific expression reference were generated by Allen Brain Institute. File human_MTG_gene_expression_matrices_2018-06-14.zip was downloaded from http://celltypes.brain-map.org/api/v2/well_known_file_download/694416044.

The gene expression and clinical data of Schizophrenia patients and healthy controls were generated by CommonMind Consortium (CMC) and the relevant data were obtained from the following links. Data access is governed by the NIMH Repository and Genomics Resources.

CMC gene expression data:

https://www.synapse.org/#!Synapse:syn3346749

CMC gene expression meta data:

https://www.synapse.org/#!Synapse:syn18103174

CMC clinical data:

https://www.synapse.org/#!Synapse:syn3275213.

The gene expression and clinical data of Autism patients and healthy controls were part of The PsychENCODE (PEC) Capstone Collection https://www.synapse.org/#!Synapse:syn12080241 and the relevant data were obtained from the following links. Data access is governed by the NIMH Repository and Genomics Resources.

UCLA-ASD gene expression data:

https://www.synapse.org/#!Synapse:syn8365527

UCLA-ASD gene expression meta data:

https://www.synapse.org/#!Synapse:syn5602933

UCLA-ASD clinical data:

https://www.synapse.org/#!Synapse:syn5602932

The list of SFARI ASD risk genes were downloaded from

https://gene.sfari.org/database/human-gene/

Source Data for Figures 2–5 are available with this manuscript.

### Code availability

The codes for generating CT-specific gene expression reference panel are included in GitHub repository `scRNAseq_pipelines` (https://github.com/Sun-lab/scRNAseq_pipelines). we have analyzed three scRNA-seq datasets: MTG, dronc, and psychENCODE, and the codes were saved in corresponding folders.

The codes to compare different references and generate final references were saved in folder `_brain_cell_type`.

The codes for CARseq analyses (including simulation, and analyses of SCZ and ASD datasets) were included in GitHub repository `CARseq_pipelines` (https://github.com/Sun-lab/CARseq_pipelines). The file `reproducible_figures.html` has the code to generate most Figures in this paper. The R package CARseq were deposited at GitHub repository `CARseq` (https://github.com/Sun-lab/CARseq).

## Notes

https://github.com/Sun-lab/scRNAseq_pipelines

### Competing Interest Statement

The authors have declared no competing interest.

