## Supplementary Materials for "Cell type-aware analysis of RNA-seq data"

### Supplementary Materials for "CARseq: Cell Type Aware analysis of RNA-seq"

**C Supplementary Tables** **32**

**D Supplementary Figures** **36**

### A Supplementary Notes

#### A.1 Optimization of regression coefficients through IWLS

For each gene, we have $K + MH$ regression coefficients and an overdispersion parameter $\phi_j$ to estimate. We will have an initial value of $\phi_j$, and estimate regression coefficients and the overdispersion parameter until convergence.

From now on, assume we already know the overdispersion parameter. The problem is to maximize the log-likelihood with respect to regression coefficients $\beta_{jk}$ and $\gamma_{jhm}$.

Let the cell type-specific mean expression weighted by cell fractions be

$$\mu_{jih} := d_i \exp\left(\sum_{k=1}^{K} \beta_{jk} w_{ik}\right) \hat{\rho}_{hi} \exp\left(\sum_{m=1}^{M} \gamma_{jhm} x_{ihm}\right),$$

satisfying

$$\mu_{ji} = \sum_{h=1}^{H} \mu_{jih},$$

then

$$\frac{\partial \mu_{jih}}{\partial \beta_{jk}} = w_{ik} \mu_{jih},$$

$$\frac{\partial \mu_{jih}}{\partial \gamma_{jhm}} = x_{ihm} \mu_{jih} = z_{jihm} \mu_{jih},$$

when we introduce $z_{jihm} := x_{ihm}$ for generality.

The bulk RNA-seq log-likelihood function is:

$$\ell_j = \sum_{i=1}^{n} \Big[ \log\Gamma(T_{ji}+\phi_j) - \log\Gamma(\phi_j) - \log\Gamma(T_{ji}+1) + \phi_j \log\phi_j + T_{ji} \log\mu_{ji} - (T_{ji}+\phi_j) \log(\phi_j+\mu_{ji}) \Big].$$

We further define

$$d_{ji} := \frac{\phi_j}{\mu_{ji}(\phi_j + \mu_{ji})},$$

$$r_{ji} := \frac{(T_{ji} - \mu_{ji})\phi_j}{\mu_{ji}(\phi_j + \mu_{ji})}.$$

Once we have defined $\mu_{jih}$, $z_{jihk}$ and $r_{ji}$, we can rewrite the score function:

$$\frac{\partial \ell_j}{\partial \beta_{jk}} = \sum_{i=1}^{n} w_{ik} r_{ji} \mu_{ji},$$
$$\frac{\partial \ell_j}{\partial \gamma_{jhm}} = \sum_{i=1}^{n} z_{jihm} r_{ji} \mu_{jih},$$

Now we derive the information matrix. First, let

$$B_{jik} := \frac{\partial r_{ji}}{\partial \beta_{jk}} = \frac{w_{ik}\phi_j \mu_{ji}(\mu_{ji}^2 - 2T_{ji}\mu_{ji} - T_{ji}\phi_j)}{\mu_{ji}^2(\mu_{ji}+\phi_j)^2},$$

$$G_{jigp} := \frac{\partial r_{ji}}{\partial \gamma_{jgp}} = \frac{z_{jigp}\phi_j \mu_{jig}(\mu_{ji}^2 - 2T_{ji}\mu_{ji} - T_{ji}\phi_j)}{\mu_{ji}^2(\mu_{ji}+\phi_j)^2}.$$

Second,

$$\frac{\partial^2 \ell_j}{\partial \beta_{jp}\beta_{jk}} = \sum_{i=1}^{n}\left[B_{jip} w_{ik}\mu_{ji} + \frac{\partial(w_{ik}\mu_{ji})}{\partial \beta_{jp}} r_{ji}\right] = \sum_{i=1}^{n}\left[B_{jip} w_{ik}\mu_{ji} + w_{ik}w_{ip}\mu_{ji} r_{ji}\right],$$

$$\frac{\partial^2 \ell_j}{\partial \gamma_{jgp}\beta_{jk}} = \sum_{i=1}^{n}\left[G_{jigp} w_{ik}\mu_{ji} + \frac{\partial(w_{ik}\mu_{ji})}{\partial \gamma_{jgp}} r_{ji}\right] = \sum_{i=1}^{n}\left[G_{jigp} w_{ik}\mu_{ji} + w_{ik}z_{jigp}\mu_{jig} r_{ji}\right],$$

$$\frac{\partial^2 \ell_j}{\partial \gamma_{jgp}\gamma_{jhm}} = \sum_{i=1}^{n}\left[G_{jigp} z_{jihm}\mu_{jih} + \frac{\partial(z_{jihm}\mu_{jih})}{\partial \gamma_{jgp}} r_{ji}\right].$$

Then we calculate the expectations with respect to the observed read counts $T_{ji}$. Observe that

$$\mathrm{E}[r_{ji}] = 0,$$
$$\mathrm{E}[B_{jik}] = -\frac{w_{ik}\phi_j\mu_{ji}}{\mu_{ji}(\mu_{ji}+\phi_j)},$$
$$\mathrm{E}[G_{jigp}] = -\frac{z_{jigp}\phi_j\mu_{jig}}{\mu_{ji}(\mu_{ji}+\phi_j)}.$$

It follows that

$$\mathrm{E}\left[\frac{\partial^2 \ell_j}{\partial \beta_{jp}\beta_{jk}}\right] = -\sum_{i=1}^{n}\frac{w_{ik}w_{ip}\phi_j\mu_{ji}^2}{\mu_{ji}(\mu_{ji}+\phi_j)} = -\sum_{i=1}^{n} w_{ik}w_{ip}d_{ji}\mu_{ji}^2,$$

$$\mathrm{E}\left[\frac{\partial^2 \ell_j}{\partial \gamma_{jgp}\beta_{jk}}\right] = -\sum_{i=1}^{n} \frac{w_{ik} z_{jigp} \phi_j \mu_{ji} \mu_{jip}}{\mu_{ji}(\mu_{ji} + \phi_j)} = -\sum_{i=1}^{n} w_{ik} z_{jigp} d_{ji} \mu_{ji} \mu_{jip},$$

$$\mathrm{E}\left[\frac{\partial^2 \ell_j}{\partial \gamma_{jgp}\gamma_{jhm}}\right] = -\sum_{i=1}^{n} \frac{z_{jihm} z_{jigp} \phi_j \mu_{jim} \mu_{jip}}{\mu_{ji}(\mu_{ji} + \phi_j)} = -\sum_{i=1}^{n} z_{jihm} z_{jigp} d_{ji} \mu_{jih} \mu_{jip}.$$

We now have the information matrix. Let us define $D_j$ as the $n$-dimension diagonal matrix with diagonal elements as $d_{ji},\ i \in \{1, 2, \ldots, n\}$. Define $X_j$ as matrix with $n$ rows and $K + HM$ columns:

$$\begin{pmatrix}
\mu_{j1} w_{11} \cdots & \mu_{j1} w_{1k} & \cdots & \mu_{j1} w_{1K} & \mu_{j11} z_{j111} & \mu_{j12} z_{j121} & \cdots & \mu_{j1h} z_{j1hm} & \cdots & \mu_{j1H} z_{j1HM} \\
\vdots & & & \vdots & \vdots & & & & & \vdots \\
\mu_{ji} w_{i1} \cdots & \mu_{ji} w_{ik} & \cdots & \mu_{ji} w_{iK} & \mu_{ji1} z_{ji11} & \mu_{ji2} z_{ji21} & \cdots & \mu_{jih} z_{jihm} & \cdots & \mu_{jiH} z_{jiHM} \\
\vdots & & & \vdots & \vdots & & & & & \vdots \\
\mu_{jn} w_{n1} \cdots & \mu_{jn} w_{nk} & \cdots & \mu_{jn} w_{nK} & \mu_{jn1} z_{jn11} & \mu_{jn2} z_{jn21} & \cdots & \mu_{jnh} z_{jnhm} & \cdots & \mu_{jnH} z_{jnHM}
\end{pmatrix}$$

Then the information matrix is

$$X_j^T D_j X_j.$$

Define the column vector with $(K + HM)$ rows when we concatenate $\beta_{jk},\ k \in \{1, 2, \ldots, K\}$ and $\gamma_{jhm},\ h \in \{1, 2, \ldots, H\},\ m \in \{1, 2, \ldots, M\}$ as $\tilde{\boldsymbol{\beta}}_j$, and the column vector with $n$ rows as $\boldsymbol{Y}_j^*$ whose each entry is $T_{ji} - \mu_{ji}$. Then the score function is

$$X_j^T D_j \boldsymbol{Y}_j^*.$$

With the information matrix and score function formulated, using Fisher scoring method,

$$\tilde{\boldsymbol{\beta}}_j^{(k+1)} = \tilde{\boldsymbol{\beta}}_j^{(k)} + \left(X_j^{(k)^T} D_j^{(k)} X_j^{(k)}\right)^{-1} X_j^{(k)^T} D_j^{(k)} \boldsymbol{Y}_j^{*(k)},$$

or equivalently

$$\tilde{\boldsymbol{\beta}}_j^{(k+1)} = \left(X_j^{(k)^T} D_j^{(k)} X_j^{(k)}\right)^{-1} X_j^{(k)^T} D_j^{(k)} \left(X_j^{(k)} \tilde{\boldsymbol{\beta}}_j^{(k)} + \boldsymbol{Y}_j^{*(k)}\right).$$

Define

$$\boldsymbol{Z}_j^{(k)} = X_j^{(k)} \tilde{\boldsymbol{\beta}}_j^{(k)} + \boldsymbol{Y}_j^{*(k)},$$

Now we see that $\tilde{\boldsymbol{\beta}}_j^{(k+1)}$ is the solution of a weighted least squares with response vector $\boldsymbol{Z}_j^{(k)}$, design matrix $X_j^{(k)}$, and weights $D_j^{(k)}$. Applying this iteratively, we should now have an optimization algorithm with quadratic convergence speed.

#### A.2 Practical considerations in IWLS

In practice, however, the vanilla version of IWLS will not work as intended, to put it mildly. From now on, we will focus on a typical use case where $x_{ihm}$'s are group labels taking binary $\{0, 1\}$ values. Unlike negative binomial regression without cell types, where IWLS will generally converge nicely to a finite estimate as in a typical GLM framework, now we would start to see cases that the estimates of some $\gamma_{jhm}$'s tend to negative infinity. This phenomenon is more prominent when the sample size is smaller or the expected cell fraction is lower. Intuitively, since deconvolution is done on a non-log scale [Zhong and Liu, 2012], when the standard error of cell type-specific expression on a non-log scale is large, the estimate of cell type-specific expression may become zero or even negative without a non-negativity constraint implied through the log-scale modeling of expression. Meanwhile, some $\mu_{jih}$'s may decrease to 0, which means some columns of $X_j^{(k)}$ may decrease to 0's, thus the condition number of ${X_j^{(k)}}^T D_j^{(k)} X_j^{(k)}$ becomes either extremely large or infinity. As a result, the IWLS iterations will get stuck somewhere when some regression coefficient estimates reach the vicinity of negative infinity, and where it may get stuck depends on the initial values.

One issue standing out is that the negative log-likelihood is not always decreasing during the iterations. To solve the issue, recall that the search direction is:

$$\boldsymbol{p}^{(k)} = \left( {X_j^{(k)}}^T D_j^{(k)} X_j^{(k)} \right)^{-1} {X_j^{(k)}}^T D_j^{(k)} \boldsymbol{Y}_j^{*(k)}.$$

We find the step size $\rho^u$ that guarantees sufficient decrease in negative log-likelihood, also called the Armijo condition [Nocedal and Wright, 2006]:

$$-\ell_j(\tilde{\boldsymbol{\beta}}_j^{(k)} + \rho^u \boldsymbol{p}^{(k)}) \leq -\ell_j(\tilde{\boldsymbol{\beta}}_j^{(k)}) + c\rho^u [-\dot{\ell}_j(\tilde{\boldsymbol{\beta}}_j^{(k)})]^T \boldsymbol{p}^{(k)},$$

where we take $c$ as 1e$-$4, $\rho$ as 0.5, and $u$ as the smallest non-negative integer that makes the inequality hold. The process of finding the right step size is called backtracking line search.

While backtracking line search will facilitate the convergence of iterations, it does not directly address the singularity problem. To solve this, our approach ("optimization on a non-log scale") is to re-parametrize the cell type-specific regression coefficients by letting $\widetilde{\gamma}_{jhm} = \exp(\gamma_{jhm})$, $\widetilde{\gamma}_{jhm} \geq 0$:

$$\widetilde{\mu}_{jih} = d_i \exp\left( \sum_{k=1}^{K} \beta_{jk} w_{ik} \right) \prod_{m=1}^{M} \widetilde{\gamma}_{jhm}^{x_{ihm}}.$$

Then we minimize the negative log-likelihood using IWLS exactly as what is stated above, except that we need to set $z_{jihm} = x_{ihm}/\gamma_{jhm}$ and that the weighted least squares performed at each iteration need to be replaced with non-negative weighted least squares. Bounded variable least squares [Stark and L. Parker, 1995] specifying $\widetilde{\gamma}_{jhm} \geq$ 1e−30 is the actual implementation being adopted here. When boundary restraints are taken good care of, the IWLS algorithm with backtracking line search and non-negative least squares will converge to the MLE fairly easily. We further note that although not the default option used in the analyses, setting an uninformative prior of $N(0, 1\text{e}6)$ and using iteratively reweighted ridge regression will also lead to the convergence to the same MLE as we obtained using optimization on a non-log scale. This is the optimization strategy used in DESeq2, and more on that will be covered in the "Shrunken log fold change" section.

##### A.3 Estimation of the overdispersion parameter

To initialize the overdispersion parameter, we take the overdispersion parameter estimated by sampling $x^*_{im}$ from $x_{ihm}, h \in \{1, \ldots, H\}$ and solving a negative binomial regression: $T_{ji} \sim f_{NB}(\mu_{ji}, \phi_j)$ and $\mu_{ji} = d_i \exp\left(\sum_{k=1}^K \beta_{jk} w_{ik} + \sum_{m=1}^M \gamma^*_{jm} x^*_{im}\right)$. We take the MLE of $\phi_j$ as the initial value of the overdispersion parameter. The MLE of $\beta_{jk}$ is also used to initialize the regression coefficients $\beta_{jk}$, and the MLE of $\gamma^*_{jm}$ is replicated across $H$ cell types to populate the initial value of $\gamma_{jkm}$.

The bulk RNA-seq log-likelihood is:

$$\ell_j = \sum_{i=1}^n \Big[ \log \Gamma(T_{ji}+\phi_j) - \log \Gamma(\phi_j) - \log \Gamma(T_{ji}+1) + \phi_j \log \phi_j + T_{ji} \log \mu_{ji} - (T_{ji}+\phi_j) \log(\phi_j+\mu_{ji}) \Big].$$

And define the Cox-Reid adjusted profile log-likelihood as:

$$\ell_j^{\text{CR}}(\phi_j) = \ell_j(\phi_j) - \frac{1}{2} \log(\det(X_j^T D_j(\phi_j) X_j))$$

where $D_j(\phi_j)$ and $X_j$ have been defined in the previous section "Practical considerations in IWLS" and their calculation involves combining cell type-specific covariates, cell type-independent covariates, and estimated cell fractions.

The Cox-Reid approximate conditional inference [Cox and Reid, 1987] was proposed to estimate negative binomial overdispersion in the analysis of SAGE data to control type I error in small-sample tests [Robinson and Smyth, 2008]. The adjustment term $-\frac{1}{2} \log(\det(X_j^T D_j(\phi_j) X_j))$ is derived from the observed information of $\phi_j$. Since then, the conditional maximum likelihood estimate has been used by edgeR [Robinson et al., 2010] and DESeq2 [Love et al., 2014] to reduce the bias of overdispersion estimation, and thus the hypothesis testing of differential expression becomes more conservative than using the MLE of overdispersion parameter.

The overdispersion parameter is updated after the regression coefficients are estimated using IWLS by doing a one-dimensional optimization on the adjusted profile likelihood of $\phi_j$ introduced above. The process of updating regression coefficients and then the overdispersion parameter will continue until the overdispersion parameter will not be changed for more than 0.1 on a log scale.

Since cell type-specific differential expression usually requires a larger sample size to detect cell type-specific expression variability, overdispersion estimation is done gene by gene. This is different from the default options of edgeR [Robinson et al., 2010] and DESeq2 [Love et al., 2014], where mean-overdispersion relationship across genes is leveraged to generate moderated estimation of overdispersion parameters. This approach could improve the sensitivity and specificity of differential expression tests when dealing with very small sample sizes. However, with sample of modest size, there is not a clear-cut case supporting its necessity [Zhou et al., 2011].

##### A.4 Cell type-specific tests using likelihood ratio statistics

We introduce likelihood ratio statistics to test cell type-specific expression. Although the general testing framework is very flexible to be inclusive of all kinds of tests discussed in TOAST [Li et al., 2019], we only emphasize on the application that the test is about whether there is cell type-specific differential expression analysis across groups.

Consider the cell type-specific expression:

$$\widetilde{\mu}_{jih} := d_i \exp\left(\sum_{k=1}^{K} \beta_{jk} w_{ik}\right) \exp\left(\sum_{m=1}^{M} \gamma_{jhm} x_{ihm}\right).$$

Under the full model, $\gamma_{jhm}$ is the cell type-specific mean expression in cell type $h$ and group $m$, controlled for other covariates. Then $x_{ihm} = 1$ when sample $i$ belongs to group $m$, and $x_{ihm} = 0$ otherwise.

Under the reduced model, within a certain cell type $h$, there is no differential expression across groups. Instead, the expression is always $\gamma_{jhm}$ regardless of which group sample $i$ belongs to. Without loss of generality, let us suppose that the cell type-specific expression stays at $\gamma_{jh1}$. In this way, $x_{ih1} = 1$ and $x_{ihm} = 0$, $m \in \{2, \ldots, M\}$, and the corresponding $\gamma_{jhm}$'s are no longer possible to estimate since they are not included in the reduced model as a matter of fact. Thus, during estimation of regression coefficients, these pairs of $h$ and $m$ are left out of the IWLS algorithm.

Once we have the log-likelihood under the full model and the reduced model, we can calculate the likelihood ratio statistics and compare it to a chi-squared distribution with $M-1$ degrees of freedom. This approximation to the distribution of the statistics is asymptotic (a $t$ distribution has been proposed as an alternative in the literature), and is only appropriate when the true parameter is not on the boundary. Biologically speaking, the cell type-specific expression should indeed be always positive. However, in a finite sample case, the closeness to a boundary case might change the null distribution into a mixture distribution without a closed-form expression and more complex than a mixture chi-squared distribution [Self and Liang, 1987,Molenberghs and Verbeke, 2007]. With that being said, when the sample size is moderately large, any departure from the asymptotic distribution is commonly too small to warrant much scrutiny.

##### A.5 Shrunken log fold change

While a sufficiently small $p$-value from a differential expression test can be interpreted as statistical significance in the association between expression and the parameter of interest, it does not indicate the biological strength of association. Log fold change (LFC) is up to this task. It is customary to plot LFC and $p$-value of genes on a scatterplot—aptly named as volcano plot—and dictate which genes are candidates for further investigation using both LFC and $p$-value thresholds. LFC can also be used in gene ontology (GO) or gene set enrichment analysis (GSEA).

Nevertheless, the interpretation of LFC can be difficult especially when the sample size is small and the variability is large. Smaller studies tend to report large LFCs even when the differential expression is absent [The Brainstorm Consortium et al., 2018]. Shrunken log fold change is a stabilized estimation of the strength of differential expression. Notably, since deconvolution is done on a non-log scale, without any shrinkage, the estimated cell type-specific expression can go to zero due to the (implicit) constraint that cell type-specific expression is non-negative, and the raw LFC goes to positive or negative infinity. A similarly hard-to-interpret scenario can be found in TOAST results, where the estimated cell type-specific expression can even become negative, and thus LFC has no definition. These ill-posed problems severely limit the use cases of raw LFC to quantify the strength of cell type-specific expression. With a prior that is sufficiently informative, such as an empirical Bayes prior, the shrunken log fold change is generally finite and easier to interpret. Furthermore, if the whole experiment is replicated or if it is replicated with a larger sample size, the consistency between replicated studies would be better if the shrunken LFC, instead of the raw LFC, is used as the measure of the strength of differential expression [Love et al., 2014].

The shrunken log fold change implemented in CARseq largely follows that in DESeq2 [Love et al., 2014], though there are some subtleties of CARseq implementation. The additive structure of mixture expression depending on cell type-specific expression in CARseq requires some weak shrinkage on the cell type-specific group mean. Though an unconventional choice, it compresses the estimated group mean towards zero on a log scale and mitigates the optimization difficulties when any estimate of cell type-specific expression approaches the boundary. Another pertinent distinction lies in CARseq forgoing the moderation of overdispersion parameters. Although this does not affect the raw LFC, the adaptation of moderated overdispersion parameter can further stabilize the shrunken LFC, resulting the impression that DESeq2 produces more aggressively shrunken log fold change.

Recall that when setting the design matrix, to code a factor with multiple levels, the default choice is to set the first level as intercept and code each other level as a contrast with the first. While the designation of which level as intercept will not affect estimation and testing without a prior, the asymmetric application of prior to contrasts will be problematic. Therefore, we borrow the concept of expanded design matrix from DESeq2. Essentially, we set the group mean $\gamma_{jh0}$ of cell type-specific expression as intercept, and code each level as a contrast with the group mean by $\gamma_{jhm}$. Without setting a prior on the contrasts, the design matrix is singular and the linear model is not estimable. With a prior on the contrasts, however, the estimation can still proceed. The contrasts have been applied with a comparatively strong prior and the group mean has a weaker prior.

In a normal design matrix, there are $HM$ cell type-specific variables. In an expanded design matrix, for each cell type $h$, there are $M$ contrasts $\gamma_{jhm}$ and one group mean $\gamma_{jh0}$, so there are $H(M+1)$ cell type-specific variables altogether. When applying the empirical Bayes prior to log fold change, the $K$ cell type-independent variables are not being penalized on. This is because we assume the parameters of interest are the cell type-specific ones. The posterior of LFC will be shrunken towards zero with a prior of zero-centered normal distribution:

$$\widehat{\boldsymbol{\beta}_j}^{\text{MAP}} = \arg\max \left[ \ell_j(\tilde{\boldsymbol{\beta}}_j^*) + P(\tilde{\boldsymbol{\beta}}_j^*) \right],$$

where $\tilde{\boldsymbol{\beta}}_j^*$ is the column vector with $K + H(M+1)$ rows when we concatenate

$\beta_{jk},\ k \in \{1, 2, \ldots, K\}$ and $\gamma_{jhm},\ h \in \{0, 1, 2, \ldots, H\},\ m \in \{1, 2, \ldots, M\}$, and

$$P(\tilde{\boldsymbol{\beta}}_j^*) = \sum_{h=1}^{H} \frac{-\gamma_{jh0}^2}{2\sigma_0^2} + \sum_{h=1}^{H}\sum_{m=1}^{M} \frac{-\gamma_{jhm}^2}{2\tau_h^2}.$$

Here $\sigma_0^2$ is chosen to be $10^4$ so that we have a relatively weak prior that can offer enough shrinkage on the group mean to avoid numerical issues. $\sigma_h^2$ is obtained by fitting the MLE of the unshrunken contrasts, i.e., the cell type-specific effect estimates that are still finite on a log scale, to a zero-centered Gaussian distribution.

To estimate the empirical Bayes prior, we first estimate the MLEs on a log scale of cell type-specific expression across genes. Any infinite values are excluded in the calculation of empirical distribution afterwards. We set a comparatively weak prior on the cell type-specific group means as $N(0, 10^4)$. The inclusion of such a prior is to require the posterior of the cell type-specific group means to be finite. Then we compute all combinations of contrasts between levels of each cell type $h$, collect the constrasts belonging to each cell type together, and align the 0.95 quantile of the empirical distribution of the absolute value of all the cell type-specific contrasts with the 0.975 quantile of a zero-centered normal distribution $N(0, \sigma^2)$. Note that the prior is the same for different cell types, because we do not have a preponderance of evidence favoring the other choice. We now have a comparatively strong prior on the cell type-specific contrasts.

The update rule in iteratively reweighted ridge regression [Park, 2006, Love et al., 2014] is:

$$\tilde{\boldsymbol{\beta}}_j^{*(k+1)} = \left( X_j^{*(k)^T} D_j^{(k)} X_j^{*(k)} + \boldsymbol{\lambda} I \right)^{-1} X_j^{*(k)^T} D_j^{(k)} \left( X_j^{*(k)} \tilde{\boldsymbol{\beta}}_j^{(k)} + \boldsymbol{Y}_j^{*(k)} \right),$$

where $I$ is the identity matrix, $\boldsymbol{\lambda}$ is a vector of length $K + H(M+1)$ obtained by taking $\tilde{\boldsymbol{\beta}}_j^*$ and replacing $\beta_{jk}$ with 0 and $\gamma_{jhm}$ with $1/\sigma_m^2$, $D_j$ is the $n$-dimension diagonal matrix with diagonal elements as $d_{ji} = \frac{\phi_j}{\mu_{ji}(\phi_j + \mu_{ji})}$, $i \in \{1, 2, \ldots, n\}$, $\boldsymbol{Y}_j^*$ is a column vector with $n$ rows whose each entry is $T_{ji} - \mu_{ji}$. , and $X_j^*$ is a matrix with $n$ rows and $K + H(M+1)$ columns:

$$\begin{pmatrix} \mu_{j1}w_{11} \cdots & \mu_{j1}w_{1k} & \cdots & \mu_{j1}w_{1K} & \mu_{j11}x_{110} & \mu_{j12}x_{120} & \cdots & \mu_{j1h}x_{1hm} & \cdots & \mu_{j1H}x_{1HM} \\ \vdots & & & \vdots & \vdots & & & & & \vdots \\ \mu_{ji}w_{i1} \cdots & \mu_{ji}w_{ik} & \cdots & \mu_{ji}w_{iK} & \mu_{ji1}x_{i10} & \mu_{ji2}x_{i20} & \cdots & \mu_{jih}x_{ihm} & \cdots & \mu_{jiH}x_{iHM} \\ \vdots & & & \vdots & \vdots & & & & & \vdots \\ \mu_{jn}w_{n1} \cdots & \mu_{jn}w_{nk} & \cdots & \mu_{jn}w_{nK} & \mu_{jn1}x_{n10} & \mu_{jn2}x_{n20} & \cdots & \mu_{jnh}x_{nhm} & \cdots & \mu_{jnH}x_{nHM} \end{pmatrix}.$$

##### A.6 Relation between CARseq model and TOAST model

The major difference between CARseq and TOAST is that the former employs a negative binomial distribution while the latter uses a linear model that is consistent with normal distribution assumption. In addition, the way to connect the mean expression in bulk tissue to CT-specific expression are different. Here we demonstrate the relation of these two approaches.

Note that $\mu_{ji}$ is the expected total read count of gene $j$ and sample $i$. As TOAST does not allow for cell type-independent variables $\beta_{jk}$'s, which are actually possible to be recast into cell type-specific variables at the expense of added degrees of freedom, and it does not model sample-level read depth $d_i$'s, we will adjust for them when comparing the CARseq model and the TOAST model.

$$\frac{\mu_{ji}}{d_i \exp\left(\sum_{k=1}^{K} \beta_{jk} w_{ik}\right)} = \sum_{h=1}^{H} \hat{\rho}_{hi} \exp\left(\sum_{m=1}^{M} \gamma_{jhm} x_{ihm}\right),$$

Let $\xi_{jhm} = \exp(\gamma_{jhm}) - 1$. It follows that

$$\frac{\mu_{ji}}{d_i \exp\left(\sum_{k=1}^{K} \beta_{jk} w_{ik}\right)} = \sum_{h=1}^{H} \left[\hat{\rho}_{hi} \prod_{m=1}^{M} (1 + \xi_{jhm})^{x_{ihm}}\right].$$

When $|\xi_{jhm}| \ll 1$ and $|x_{ihm}\xi_{jhm}| \ll 1$, the above equation can be approximated by:

$$\sum_{h=1}^{H} \left[\hat{\rho}_{hi} \prod_{m=1}^{M} (1 + x_{ihm}\xi_{jhm})\right] \approx \sum_{h=1}^{H} \left[\hat{\rho}_{hi} \Big(1 + \sum_{m=1}^{M} x_{ihm}\xi_{jhm}\Big)\right].$$

In the typical circumstance where $x_{ihm} \in \{0, 1\}$ are group indicators, the "approximately equal to" relations are all "equal to" relations.

We now arrive at a form very close to how the measurement in $TOAST$ is modeled with cellular proportion as main effects, which are parametrized below using $\eta_{hi}$'s, and proportion by covariate as interactions:

$$\sum_{h=1}^{H} \left[\sum_{m=1}^{M} (\hat{\rho}_{hi}\eta_{jh} + x_{ihm}\hat{\rho}_{hi}\xi_{jhm})\right].$$

##### A.7 Connections between cell fractions, cell type-specific transcript fractions and cell size

We explain how to convert between the proportion of cells, or the cell fractions in the model involving read counts, $\rho_{hi}$, and the proportion of transcripts from a cell, or cell fractions in the model involving TPM, $\rho_{hi}^{\text{TPM}}$. Suppose $j$ is a subscript for gene, $i$ is a subscript for sample, and $h$ is a subscript for cell types. The reference of purified cell types defined by counts is denoted using $\gamma_{jh}$. Gene lengths are $\ell_j$.

When we obtain cell fractions by deconvolving mixture expression in TPM, such as by CIBERSORT or ICeD-T, we need to adjust for cell size to obtain cell fractions in the literal sense.

Suppose the total read counts follow a negative binomial distribution:

$$T_{ji} \sim f_{NB}(d_i \sum_{h=1}^{H} \rho_{hi}\gamma_{jh}, \phi_j).$$

Taking expectation gives:

$$\text{E}[T_{ji}] = d_i \sum_{h=1}^{H} \rho_{hi}\gamma_{jh},$$

$$\text{E}[T_{ji}/\ell_j] = d_i \sum_{h=1}^{H} (\rho_{hi}\gamma_{jh}/\ell_j).$$

Since the total number of genes $G$ is very large, using Approximations for Mean and Variance of a Ratio, we get:

$$\text{E}\Big[\frac{T_{ji}/\ell_j}{\sum_{j=1}^{G}(T_{ji}/\ell_j)}\Big] \approx \frac{\text{E}[T_{ji}/\ell_j]}{\text{E}[\sum_{j=1}^{G}(T_{ji}/\ell_j)]} = d_i \sum_{h=1}^{H} \rho_{hi} \frac{\sum_{j=1}^{G}(\gamma_{jh}/\ell_j)}{\text{E}[\sum_{j=1}^{G}(T_{ji}/\ell_j)]} \frac{(\gamma_{jh}/\ell_j)}{\sum_{j=1}^{G}(\gamma_{jh}/\ell_j)}.$$

Let $r_{hi} = \dfrac{\sum_{j=1}^{G}(\gamma_{jh}/\ell_j)}{\text{E}[\sum_{j=1}^{G}(T_{ji}/\ell_j)]}$, and $\gamma_{jh}^{\text{TPM}} = \dfrac{(\gamma_{jh}/\ell_j)}{\sum_{j=1}^{G}(\gamma_{jh}/\ell_j)}$ by the definition of TPM.

Then we have mixture expression and cell type-specific expression in TPM:

$$\text{E}[T_{ji}^{\text{TPM}}] = d_i \sum_{h=1}^{H} \rho_{hi} r_{hi} \gamma_{jh}^{\text{TPM}}$$

If we add up all the genes

$$\sum_{j=1}^{G} \mathrm{E}[T_{ji}^{\mathrm{TPM}}] = \sum_{j=1}^{G} d_i \sum_{h=1}^{H} \rho_{hi} r_{hi} \gamma_{jh}^{\mathrm{TPM}},$$

we get $\sum_{h=1}^{H} d_i \rho_{hi} r_{hi} = 1$. Let $\rho_{hi}^{\mathrm{TPM}} = d_i \rho_{hi} r_{hi}$, which follows that

$$\rho_{hi}^{\mathrm{TPM}} \propto \rho_{hi} \sum_{j=1}^{G} (\gamma_{jh}/\ell_j).$$

This justifies us to define the total number of transcripts in a cell:

$$s_h = \sum_{j=1}^{G} (\gamma_{jh}/\ell_j)$$

as the cell size to convert between $\rho_{hi}^{\mathrm{TPM}}$ and $\rho_{hi}$.

### B Supplementary Materials for Data Analyses

#### B.1 Details of simulations

##### B.1.1 Methods to benchmark

1. CARseq (w/ and w/o clinical variables)

   `R/simulation_test_CARseq.R`

   Input 1: read counts; true cell fractions.

   Input 2: read counts; cell fractions from ICeD-T and adjusted for cell size.

   Output: p-value for every gene/cell type pair.

   To use CARseq, the cell fraction estimates are required. We recommend to use ICeD-T or CIBERSORT to obtain the cell fraction estimates $\hat{\rho}_{hi}$ that sum up to 1 for every sample $i$. Here the cell fractions are true cell fractions, which is to say, if ICeD-T or CIBERSORT is used, the fractions need to be adjusted for cell size factors.

   CARseq has the a higher power than csSAM while controlling for type I error. One possible reason is CARseq can leverage discrete distributions, which would provide a higher power. Another reason is csSAM does not allow for the incorporation of known batch effects. The third reason is csSAM uses permutation test, and the power is generally lower than model-based tests when the model holds for the data.

2. csSAM (w/o clinical variables) csSamWrapper with nonNeg = TRUE as other methods does not allow negative expression values

   `R/simulation_test_csSAM.R`

   Input: TPM; the true cell fractions in the scale of TPM (can be alternatively called the cell type-specific transcript fractions).

   When we assume the cell sizes are all the same across different cell types, the cell fractions do not need to be adjusted.

   Output: FDR for every gene/cell type pair.

3. TOAST (w/ and w/o clinical variables; using TPM)

   `R/simulation_test_TOAST_TPM.R`

   Input 1: read counts; true cell fractions.

   Input 2: TPM; true cell fractions in the scale of TPM.

Output: p-value for every gene/cell type pair.

TOAST uses a linear regression model. Based on our experience, it is better to supply TPM instead of read counts, as TPM is already adjusted for read depth, while the read depth needs to be included as a covariate when modeling read counts. Read counts, in contrast, is better to be modeled using a generalized linear model to fully utilize the mean-variance structure. Unless specifically mentioned, we use TPM instead of read counts as the input of TOAST. The code to run TOAST using read counts is included as `R/simulation_test_TOAST_count.R`.

As TOAST used a linear regression model, it actually allows for negative estimates of cell type-specific expression. While this is not biologically possible, it does not affect statistical modeling. This is not the case in CARseq, where the cell type-specific expression has to be non-negative to satisfy the model requirements of a non-negative mean in a negative binomial regression.

TOAST sought to make the effect of additional covariates proportional to baseline expression, but due to the limitation of linear regression, it model them using interaction terms. This can quickly increase the degree of freedom when there is larger number of covariates.

##### B.1.2 Explanation inflated type I error

To only highlight major problems in the figure, we need to display the FDR in a more stable way. When its definition is ill-defined (0/0 = NaN) or almost ill-defined (the denominator, or the total discoveries, is no greater than 5), the FDR is set to 0 and the sensitivity is set to 0.

In the plots, FDR and type I error refer to false discoveries in the 8000 non-DE genes in a certain cell type. Be careful that "power" and "sensitivity" refer to discoveries in the 2000 DE genes, which can be either true or false depending on which cell type is differentially expressed; the information is coded in the "DE pattern" of fold changes between two groups.

In general, the simulation performs as expected. CARseq strikes a good balance between controlling type I error and being powerful.

We find that CARseq would produce inflated type I error in very few cases when the covariate (RIN) is not provided. The inflated type I error we see in simulation setup `n_200_DE_pattern_2_1_1_replicate_1` (without covariates) is caused by the collinearity between RIN and cell fractions of the minor cell type in one group. In general, in a regression framework, if there is considerable correlation between an effect to test and an effect not included in the model, then the problem of inflated type I error could arise. When RIN is not included in the model, the variation in expression attributed to RIN effect is instead falsely attributed to differential expression in the minor cell type.

As this symptom is not a methodological pitfall of CARseq, other algorithms can also manifest inflation of type I error when the variable to test is correlated to a batch effect not incorporated in the model. Since the cell type-specific test in csSAM is conceptually the same as CARseq without batch effects, when `CARseq_without_RIN` is plagued with inflated type I error, csSAM is also bound to fail. An example can be found in simulation setup `n_200_DE_pattern_2_1_1_replicate_1`. When randomly generated RIN is highly correlated with group labels, DESeq2 could have inflated type I error among non-DE genes. An example can be found at `DESeq2_without_RIN` in simulation setup `n_100_DE_pattern_4_1_1_replicate_1`.

##### B.1.3 Impact of noise/bias in cell type fraction estimates

While our previous simulation settings have shown the performance of different methods when cell type fractions are estimated by ICeD-T. Here we further demonstrate the impact of cell type fraction estimate noise by explicitly adding different patterns of noise/bias. First, we added a zero-centered Gaussian noise with a standard deviation of 0.1 to the cell fractions on a logit scale and rescaled the cell fractions so that their summation is 1 for each sample (Supplementary Figure 10). In this situation, CARseq has similar advantages over other methods (Supplementary Figure 11).

Next we consider potential bias of cell type fraction estimates due to mis-specification of cell size factors. The total amount of transcripts per cell may vary across cell types. Most computational methods for cell type decomposition estimate the fractions of gene expression instead of the fractions of cells from each cell type. For example, the two methods that we use (CIBERSORT and ICeD-T) estimate cell type fraction in terms of TPM. Therefore if one cell type has more transcripts than other cell types, its proportion is over-estimated. Such bias can be corrected using cell size factor, which are the relative amount of transcripts per cell across cell types. For example, if there are only two cell types, denoted by $A$ and $B$, and the total gene expression of cell type $A$ is twice of cell type $B$, then the cell size factors of cell types $A$ and $B$ are 2 and 1, respectively. We can estimate the fraction of cells from the fractions of gene expression after correcting for cell size factor. If a method models the amount of transcripts per gene (e.g., when using TPM to quantify expression level in TOAST), it needs the proportion of transcripts for each cell type and thus there is no need to correct for cell size factor. To interrogate the performance of CARseq when the cell size factor is misspecified, we intentionally applied wrong size factors (1.2, 1, 1) instead of the true ones (1, 1, 1) when evaluating CT-specific-DE. This misspecification of cell size factor slightly reduces the power of CARseq, though it still has higher power than TOAST (Supplementary Figure 12). Only under extreme and unrealistic misspecification of cell size factor (e.g., (2, 1, 1) vs. true values of (1,1,1)) does the power of CARseq drops to become similar to that of TOAST (Supplementary Figure 13).

###### B.1.4 Code description for simulations

First, `R/simulation_step1_get_distribution_of_parameters_from_real_data.R` fits models from CMC data using MTG single cell data as reference. Second, the joint distribution of fitted parameters is used to generate simulation data in `R/simulation_step2_simulate_data.R`. There are separate code snippets starting with `R/simulation_step3` to run each method in the `simulation` folder. Then we use `R/simulation_step4_compare_methods_multiple_replicates.R` to calculate metrics to compare the methods across ten replicates. The methods of using CIBERSORTx high resolution mode and TOAST with read counts are summarized in `R/simulation_step4_compare_methods_multiple_replicates.R` where only one replicate has been investigated.

##### B.2 Details of real data analysis

###### B.2.1 Prepare cell type-specific reference from single cell data

Estimates of cell fractions were obtained using a reference of cell type-specific expression constructed from MTG snRNA-seq data, which was generated using SMART-Seq v4 Ultra Low Input RNA Kit, which is an improved version of SMART-seq2 protocol [Hodge et al., 2019]. We used both ICeD-T (without weights) and CIBERSORT to estimate cell fractions.

We analyzed the MTG snRNA-seq data (`https://github.com/Sun-lab/scRNAseq_pipelines/blob/master/MTG/human_MTG.html` followed by an R code `step1_expression_signature.R` in the same folder) using a pipeline that is slightly different from the original paper, mostly based on Bioconductor workflows for scRNAseq [Lun et al., 2016]. Then we clustered all the cells/nuclei using K-means, ranging from 10 to 20 clusters. Based on manual inspection, we choose to compare the annotated cell type labels with K-means with 15 clusters, as shown below.

```
Astro Endo  Exc  Inh Micro Oligo  OPC unknown
```

|  |  |  |  |  |  |  |  |  |
|---|---|---|---|---|---|---|---|---|
| 1 | 0 | 0 | 0 | 1279 | 0 | 0 | 0 | 15 |
| 2 | 0 | 0 | 1867 | 0 | 0 | 0 | 0 | 24 |
| 3 | 0 | 1 | 8 | 11 | 0 | 310 | 1 | 12 |
| 4 | 0 | 0 | 260 | 0 | 0 | 0 | 0 | 22 |
| 5 | 287 | 0 | 12 | 1 | 0 | 2 | 4 | 21 |
| 6 | 0 | 0 | 1494 | 0 | 0 | 0 | 0 | 73 |
| 7 | 0 | 0 | 1483 | 0 | 0 | 0 | 0 | 17 |
| 8 | 0 | 0 | 1552 | 1 | 0 | 0 | 0 | 15 |
| 9 | 0 | 0 | 1 | 1210 | 0 | 0 | 0 | 4 |
| 10 | 0 | 0 | 2 | 4 | 62 | 1 | 0 | 9 |
| 11 | 0 | 0 | 0 | 835 | 0 | 0 | 0 | 3 |
| 12 | 1 | 8 | 16 | 807 | 1 | 0 | 233 | 35 |
| 13 | 0 | 0 | 326 | 1 | 0 | 0 | 0 | 7 |
| 14 | 0 | 0 | 1798 | 1 | 0 | 0 | 0 | 38 |
| 15 | 0 | 0 | 1654 | 1 | 0 | 0 | 0 | 28 |

There is a high consistency between our clustering results and annotated cell types. To generate the cell type-specific gene expression profile for each cell type, we select those clusters that are either is the largest cluster for this cell type or includes more than 200 cells of this cell type. This helps us filter out some cells for each cell type. In total, we kept 15,465 nuclei for the following analysis and they are separated into 7 cell types:

|  | Cell_Type | nCells_All |
|---|---|---|
| 1 | Inh | 4131 |
| 2 | Exc | 10434 |
| 3 | Oligo | 310 |
| 4 | OPC | 233 |
| 5 | Astro | 287 |
| 6 | Micro | 62 |
| 7 | Endo | 8 |

We compared this MTG snRNAseq data (SMART-Seq v4) with another snRNA-seq dataset generated using drop-seq technique named DroNc-seq [Habib et al., 2017] (hereafter referred to as DroNC data). We have re-analyzed DroNC data using a pipeline similar to the one for MTG data (`https://github.com/Sun-lab/scRNAseq_pipelines/blob/master/dronc/dronc_seq.html` followed by an R code `step1_expression_signature.R` in the same folder) and selected the cells belonging to each cell type by taking intersections of clustering results and cell type labels provided by Habib et al. [Habib et al., 2017]. DroNC data includes human hippocampus and PFC from five adults. There are 11,585 nuclei from 11 cell types, and we also consider a subset of them (4,536 nuclei) from PFC since it matches the tissues of bulk RNA-seq data.

| | Cell_Type | nCells_All | nCells_PFC |
|---|---|---|---|
| 1 | exCA3 | 630 | 8 |
| 2 | exCA1 | 179 | 0 |
| 3 | exPFC | 3107 | 3072 |
| 4 | ASC | 1584 | 339 |
| 5 | GABA | 1154 | 758 |
| 6 | ODC | 2582 | 167 |
| 7 | exDG | 1380 | 0 |
| 8 | END | 68 | 8 |
| 9 | MG | 223 | 19 |
| 10 | OPC | 533 | 165 |
| 11 | NSC | 145 | 0 |

In the above table, exCA3, exCA1, exPFC, and exDG are four types of excitatory neurons or glutamatergic neurons. ASC is astrocyte. GABA is GABAergic interneuron or inhibitory neuron. ODC is oligodendrocyte. END is endothelial cell. MG is microglia. OPC is oligodendrocyte precursor cell. NSC is neuronal stem cell.

Then we compare the cell type-specific gene expression data from DroNC (all cells or only the cells in PFC) vs. MTG data. We added up all the counts per gene across all the cells and make comparison, and collapse four types of excitatory neurons of DroNC data into one category. Detailed comparison of astrocyte and endothelial cell are shown in Supplementary Figures S14 and S15, respectively, and similar figures for other cell types can be found at `https://github.com/Sun-lab/scRNAseq_pipelines/tree/master/_brain_cell_type/figure`. It is clear that MTG data does not have much more cells than DroNC data, but the total read count is often 100 times more than DroNC data. Overall, gene expression of the same cell type are similar between MTG and DroNC, except endothelial, possibly due to small number endothelial cells (Supplementary Figure S16). We chose to use MTG data to generate reference since it has much higher depth and better coverage, making it more similar to bulk RNA-seq data. We exclude endothelial in our analysis since there are only 8 endothelial cells in MTG data and its expression has very weak similarity to the endothelial cells from DroNC data. The small number of cells also make the next step of selecting genes with cell type-specific expression much harder.

Next we selected around 120-130 genes per cell type for 6 cell types: excitatory neurons (Exc), inhibitory neurons (Inh), astrocyte (Astro), microglia (Micro), oligodendrocyte (Oligo), and oligodendrocyte precursor cell (OPC) by differential expression testing using MAST [Finak et al., 2015]. All the genes we chose have FDR $< 0.001$ and fold change large than 2. Among these genes, we chose those with smallest FDR and largest fold changes. Specifically, we did a percentile grid search of FC $>$ quantile(FC,pp) and FDR $<$ quantile(FDR,1-pp) until there are anywhere between 120 and 130 genes for some percentile pp.

##### B.2.2 Cell type fraction estimation

Cell fractions were estimated using both CIBERSORT and ICeD-T. The input of the cell type deconvolution methods were bulk gene expression in TPMs and reference matrix from MTG. For ICeD-T, we used the unweighted version. For CIBERSORT, we used their website interface of `https://cibersort.stanford.edu/index.php`.

Since the deconvolution was done in the scale of TPM, which have adjusted for cell level read-depth, the cell type fraction estimates from CIBERSORT of ICeD-T are the fraction of expression from each cell types, not necessary the fraction of cells, if the total amount of RNA molecules are different across cell types. For each cell type, we estimate cell size factor and such it to adjust cell fraction estimates. Similar procedure has been used in earlier studies for immune cell types [Racle et al., 2017b]. See section A.7 for details on transformation between cell fraction by expression and cell fraction by the number of cells.

##### B.2.3 Gene filtering, surrogate variables, and multiple testing correction.

In our analysis, only about 20,000 genes whose third quartile of read counts are larger than 20 are included. The sample depth is defined by the third quartile of read counts among the aforementioned genes.

The surrogate variables were calculated using the "sva" R package using a linear model with transformed data matrix as response and covariates included in the differential expression as predictors. To control for cell fractions, the log transformed cell fraction estimates from ICeDT are also included.

To annotate the genes, particularly to match gene names and calculate gene lengths to calculate TPMs, we used "`Homo_sapiens.GRCh37.70.processed.gtf.gz`" for the SCZ dataset and "`gencode.v19.annotation.gtf.gz`" for the ASD dataset as the GENCODE GTF files. Gene lengths are defined by the length of the union of all exons. The annotated gene expressions were wrapped in the "SummarizedExperiment" container.

Default options were adopted in running CARseq, TOAST, and DESeq2. The gene expression was supplied in the form of TPM for TOAST, and the cell fraction estimates were not adjusted for cell sizes. CARseq and DESeq2 required read counts, and CARseq also demanded cell fraction estimates adjusted for cell sizes. The p-values obtained from each method and cell type were transformed to q-values using function `get_qvalues_one_inflated` in the "CARseq" package.

#### B.3 Supplementary results for SCZ analysis

We estimated cell type proportions for six cell types: excitatory neurons (Exc), inhibitory neurons (Inh), astrocyte (Astro), microglia (Micro), oligodendrocyte (Oligo) using CIBERSORT [Newman et al., 2015] and ICeD-T [Wilson et al., 2019]. The estimates from these two methods are highly correlated, though with some noticeable differences (Supplementary Materials Section B.2.1, Figures S18-S19). We choose excitatory neuron as our reference cell type because it is the most abundant cell type and the results are easier to explain when studying excitation/inhibition imbalance. We examined whether relative cell fractions with respect to excitatory neuron are associated with the case-control status by a linear regression with log(cell type fraction ratio) as response and the covariates include case-control status as well as log transformed read depth, age, gender, RNAseq QC metrics, batch effects, genotype PCs, and two surrogate variables that were estimated conditioning on cell type fractions.

Since cell type fraction estimates from CIBERSORT and ICeD-T are highly correlated, we present the CARseq results using cell type fractions estimated by ICeD-T for simplicity. CARseq estimates the CT-specific expression for SCZ and control subjects separately and test DE for each gene. Before looking into the testing results for each gene, we observed that the CT-specific expression for both SCZ and control groups are strongly correlated with the CT-specific gene expression measured by snRNA-seq (Supplementary Figure 27), suggesting a reliable estimate of CT-specific gene expression from bulk tissues in a genome-wide scale. CARseq found 1 differentially expressed gene (DEG) ($q$-value $< 0.1$) in astrocytes, 138 DEGs in microglia, and 656 DEGs in oligodendrocytes (see Supplementary Figure 21 for p-value distributions). In contrast, TOAST identified 3 DEGs in inhibitory neurons, 30 DEGs in microglia, and 1 DEG in oligodendrocytes (See Supplementary Figure 23 for p-value distributions). Both methods could control type I error/FDR, indicated by the fact that if the case-control label was permuted, the only false discovery ($q$-value $< 0.1$) is 1 gene in microglia reported by CARseq, and the p-value distribution is uniform

(Figures S22 and S24). These results are consistent with our simulation results that CARseq can identify DEGs with a higher power than TOAST, while controlling FDR.

Our GSEA reveals some functional categories related with the DE genes. We have described the results for neuron cells in main text. Here we shift our attention to glial cells. For microglia, we found the pathways of innate immune system and cell cycle are enriched in the CT-specific-DE genes and they are over-expressed in SCZ subjects (Figure 4(D), Supplementary Figure 30), supporting the observations of activation of microglia in SCZ subjects [Prata et al., 2017]. It is interesting that these pathways are also enriched in oligodendrocyte, but they are down regulated in SCZ subjects (Figure 4(D), Supplementary Figure 30), suggesting inactivation of oligodendrocyte. We also found Slit-Robo signaling pathway, which is involved in the neurogenesis and migration of neuronal precursors toward the lesions, is down/up -regulated in microglia and oligodendrocyte, respectively [Kaneko et al., 2018] (Supplementary Figure 30). Our findings suggest microglia and oligodendrocyte may take different roles in this process in SCZ subjects.

##### B.4 Supplementary results for ASD analysis

Comparing relative cell type fractions (with respect to excitatory neurons) between ASD subjects and controls, we found the relative abundance of astrocyte is significantly higher in ASD subjects than controls ($p = 0.021$ and 0.024 for cell type fractions estimated by ICeD-T and CIBERSORT, respectively, Figure S34(A)). Microglia also show a trend of higher relative abundance in ASD subjects than controls. These observations support the hypothesis that pro-inflammatory maternal cytokines in the developing brain can lead to neuroinflammation and proliferation of astrocyte and microglia [Petrelli et al., 2016].

CARseq reports 232 DEGs ($q$-value $< 0.1$) in excitatory neurons and 855 DEGs in inhibitory neurons, and no DEGs in the other four cell types (Figure S35). TOAST recovers 2 DEGs in excitatory neurons and no DEGs in the other five cell types (Figure S37). We also sought to evaluate the FDR control by repeating our analysis after permuting case/control labels (Figure S36 and S38) and noticed inflation of type I error in some permutations. This is likely due to the fact that the model is mis-specified after permuting case/control labels, and small sample size and/or unaccounted covariates could further exaggerate such effects. Thus the results of this analysis should be interpreted with caution. Nevertheless, as discussed next, we observed expected functional category enrichment and some consistent signals between SCZ and ASD, suggesting our analysis in this dataset with relatively small sample size still captures meaningful signals.

First, we considered a list of 328 autism risk genes curated by Simons Foundation Autism Research Initiative (SFARI). Most of these risk genes were identified because they harbor more disruptive mutations in the ASD cases than the general population. We found that these ASD risk genes are significantly enriched among the DE genes in inhibitory neurons (p-value $5.8 \times 10^{-7}$) and excitatory neurons (p-value $3.3 \times 10^{-4}$), and they are significantly depleted among the DE genes in microglia (p-value $2.9 \times 10^{-7}$) (Figure S34(C), Table S5). Such enrichment of autism risk genes in the inhibitory/excitatory neurons are consistent with the results reported using snRNA-seq data [Velmeshev et al., 2019]. Enrichment by TOAST results is consistent with CARseq for microglia ($4.1 \times 10^{-4}$), but not significant for inhibitory neurons (p-value 0.14) or excitatory neurons (p-value 0.89). In contrast, no enrichment is found from DE analysis on bulk tissue by DESeq2 [Love et al., 2014] (p-value 0.66).

The pathways enriched in DE genes of excitatory or inhibitory neurons include more generic and broad pathways such as "neuronal system" and "antigen processing", and more specific ones such as "synthesis of PIPs " and "RAB regulation of trafficking". Both "synthesis of PIPs" and "RAB regulation of trafficking" are related to one type of glutamate receptors named AMPA receptor [McCartney et al., 2014, Hausser and Schlett, 2019]. The log fold changes of DE genes show that these two pathways are up-regulated in inhibitory neurons of ASD subjects, but down-regulated in excitatory neurons of ASD subjects (Figure S42). Such up/down-regulation pattern is much cleaner in "synthesis of PIPs" than "RAB regulation of trafficking". This suggests the relevance of AMPA activity in the pathophysiology of ASD. Genes in the antigen processing pathways tend to be up-regulated in inhibitory neurons but down-regulated in excitatory neurons (Figure S42), suggesting increased/decreased interactions with the immune system in inhibitory neurons and excitatory neurons, respectively. The enriched pathways in glial cells include those related to translation initiation, elongation, and "Response of EIF2AK4 (GCN2) to amino acid deficiency". It has been shown that dysregulation of translation can cause neurodegeneration [Ishimura et al., 2016], which is corroborated by our findings that suggest their connections with ASD.

##### B.5 Additional analysis of DESeq2 DE genes

Let $q_{\texttt{noCT}}$ and $q_{\texttt{withCT}}$ be the DESeq2 q-values before and after accounting for cell type compositions, respectively. We divided the genes into four groups:

1. $q_{\texttt{noCT}} \geq 0.1$ and $q_{\texttt{withCT}} \geq 0.1$,

2. $q_{\texttt{noCT}} \geq 0.1$ and $q_{\texttt{withCT}} < 0.1$,

3. $q_{\texttt{noCT}} < 0.1$ and $q_{\texttt{withCT}} \geq 0.1$,

4. $q_{\texttt{noCT}} < 0.1$ and $q_{\texttt{withCT}} < 0.1$.

We assess the associations between the expression of each gene and cell type proportions by performing two linear regressions with log-transformed and read-depth corrected gene expression as response. The first model includes all the covariates used in the CAR-seq analysis. The second model includes all these covariates plus log-transformed cell type composition ratios for 5 cells types with excitatory neurons as reference. The cell type compositions can explain a substantial proportion of gene expression variance after conditioning on all the covariates including the case/control status (Figures S46-S47).

For a specific gene, if its association with case/control status becomes significant after accounting for cell composition, it is likely because including composition can reduce within group variance. We expect these genes tend to have weaker associations with case/control status (given cell type composition) because otherwise they may remain significant without including cell composition. This is indeed the case (Figures S50 and S51).

##### B.6 Comparison with the ASD vs. control DE results from snRNA-seq data

Velmeshev et al. [Velmeshev et al., 2019] have collected single nucleus RNA sequencing (snRNA-seq) data from 16 control and 15 ASD donors in two brain regions: prefrontal cortex (PFC) and anterior cingulate cortex (ACC). Here we will use the data from prefrontal cortex because it matches with the brain region where the bulk RNA-seq data were collected from the UCLA-ASD study [Parikshak et al., 2016]. There are altogether 62,166 nuclei collected from prefrontal cortex of 10 control and 13 ASD donors. Velmeshev et al. classified these nuclei into 17 cell types. To facilitate the comparison with our results, we grouped them into 6 cell types as follows:

- Astro: AST-FB, AST-PP,
- Inh: IN-PV, IN-SST, IN-SV2C, IN-VIP,
- Exc: L2_3, L4, L5_6, L5_6-CC,
- Micro: Microglia,

- Oligo: Oligodendrocytes,
- OPC: OPC,

For each gene and each of the 23 individuals, we collapsed the counts across all the cells of each cell type to create pseudo bulk sample for each cell type and each individual, and then performed differential expression analysis using DESeq2 while accounting for the covariates age, sex, Seqbatch (sequencing batch), and RIN (RNA integrity number).

This is the best dataset that we can find for validation, though there are at least three notable difference of this dataset and bulk RNA-seq data.

1. Gene expression from nucleus is different from that from a whole cell because the distribution of transcripts are not the same between nucleus and cytoplasm. Currently, due to technique limitation to isolate individual cells from postmortem brains, all the single cell level gene expression data from human postmortem brains are single nucleus RNA-seq data rather than single cell RNA-seq data.
2. This dataset were generated using 10x Genomics platform, which is the most popular platform for single cell gene expression data. In this platform, gene expression is only measured at 3' or 5' instead of the whole transcript. In contrast, bulk RNA-seq quantify gene expression for the whole transcript, and thus is more robust to noise due to RNA isoform level gene expression variation and RNA degradation.
3. The observed counts from the snRNA-seq data are very sparse with most observed counts being 0, 1, or 2, which is typical for 10x Genomics platform. Most genes are only expressed in a small proportion of cells and the level of gene expression also varies a lot across cell types (Figure S52). For example, among the 18,041 genes included in our DE analysis, if we filter out genes that are expressed in at least 20% of cells for each cell type, we will select 3%, 4%, 7%, 8%, 27%, and 44% of genes in Micro, Oligo, Astro, OPC, Inh, and Exc, respectively. When performing differential expression analysis, we add up the counts from all the cells of one cell type within each individual to create cell type-specific pseudo-bulk RNA-seq data. This can reduce the sparsity in the data, though even after that, for Microglia, most genes have less than 10 counts per individual in more than 75% of the individual (Figure S53).

These difference indeed demonstrate the typical limitations of single cell gene expression data and thus justify why our method is needed.

All the codes for these analyses can be found at `https://github.com/Sun-lab/CARseq_pipelines/tree/master/R`: `step_z1_DESeq2.R` for DE analysis of snRNA-seq data and `step_z2_compare_snRNAseq.R` for comparison with CARseq/TOAST results.

##### B.7 Supplementary results for TCGA SKCM analysis

We downloaded the RNA-seq count data from NCI Genomic Data Commons Data Portal (`https://portal.gdc.cancer.gov/`), and additional data from TCGA pan cancer atlas (`https://gdc.cancer.gov/about-data/publications/pancanatlas`, file `TCGA_mastercalls.abs_tables_JSedit.fixed.txt` for tumor purity and file `TCGA-CDR-SupplementalTableS1.xlsx` for clinical data).

To estimate cell type fractions, we used a reference data derived from scRNA-seq data using SMART-Seq2 protocol [Tirosh et al., 2016]. More specifically, we used the pre-processed reference data from R package EPIC [Racle et al., 2017a]. Then we estimated cell type fractions using CIBERSORTx [Newman et al., 2015] and corrected for cell size factors using the factors reported by EPIC [Racle et al., 2017a]. These cell fraction estimates do not account for tumor cells. We further added the tumor cell fraction using the tumor purity estimates reported by the TCTA pan cancer study [Ding et al., 2018]. By design, the TCGA study seeks to collect tumor tissues, and thus the samples are purposely selected to have high tumor purity. Some of the tumor samples have tumor purity estimates as high as 0.99 or 1. For our purpose of cell type aware DE analysis, we selected those samples with tumor purity smaller than 0.9. (Figure S56).

We seek to assess DE between patients with disease-specific survival (DSS) longer than 5 years. Some patients' DSS is censored before 5 years and thus they may or may not survive 5 years. We have excluded such patients in our analysis. Of course a more appropriate approach is to include such censored data and extend our method to handle survival outcome, which is among our future works.

As illustrated in our simulations, without accounting for relevant covariates can lead to type I error. When analyzing this dataset, we initially included a basic set of covariates of age, gender, and tumor stage, and observed inflated type I error because there are a moderate number of findings after permuting 5-year-survival indicator. This is fixed after we including the tumor tissue sites covariates into our analysis. Some tumor tissue sites only have a few samples, making it hard to adjust their effects. Therefore we excluded those tumor tissue sites covering 4 or less samples. After all the aforementioned filtering, the final sample size is 173. To minimize the degree of freedom of the model, we further estimated average gene expression for each tissue site and performed hierarchical clustering of those average expression profiles (Figure S57). Based on the clustering results, we chose to divide these tissue sites into 5 clusters and use the cluster membership as a covariate in the following analysis. At q-value 0.1, we identified 260 DE genes across the 8 cell types (Figure S58). After permuting the indicator of 5-year survival, we identified 41 DE genes in total, suggesting a realized FDR of 0.16.

We accounted for 10 covariates, including read-depth, age, gender, tumor stage (a factor with 4 levels, and thus 3 degree of freedom) and tissue sites (a factor with 5 levels, and thus 4 degree of freedom). TOAST failed to run in this data because it used the interactions between all the covariates and all the cell types, which is a design matrix with 88 columns (88 = 1 (intercept) + 10 (covariates) + 7 (cell types) + $7 \times 10$), which is singular. In contrast, CARseq assumes each of these covariates has the same effect on all cell types and thus does not model their interactions with cell types. We think this setup to use all the interaction terms is not essential for TOAST and thus should not be considered as a critique of TOAST, though it does not prevent us from making comparison with TOAST in this dataset.

All the codes for these analyses can be found at `https://github.com/Sun-lab/CARseq_pipelines/tree/master/R`: `SKCM_step1_CARseq.R` for CARseq analysis and `SKCM_step2_TOAST.R` for TOAST analysis.

### C Supplementary Tables

| | SCZ | | | ASD | | |
|---|---|---|---|---|---|---|
| | CARseq | TOAST | DESeq2 | CARseq | TOAST | DESeq2 |
| Astro | 1 | 0 | | 0 | 0 | |
| Exc | 0 | 0 | | 232 | 2 | |
| Inh | 0 | 3 | | 835 | 0 | |
| Micro | 138 | 30 | | 0 | 0 | |
| Oligo | 656 | 1 | | 0 | 0 | |
| OPC | 0 | 0 | | 0 | 0 | |
| bulk | | | 1009/1888/810 | | | 1063/481/185 |

Table S1: Number of differentially expressed genes by cell type and method using a $q$-value cutoff of 0.1, including DESeq2 for bulk samples. The results of DESeq2 include 3 numbers: the number of findings without accounting for cell type compositions, after accounting for cell type compositions, and their intersection.

| | SCZ (#case = 250, #ctrl = 277) | | | ASD (#case = 42, #ctrl = 43) | | |
|---|---|---|---|---|---|---|
| | Estimate | Std. error | $p$-value | Estimate | Std. error | $p$-value |
| Astro | | | | | | |
| ICeDT | −0.003 | 0.027 | 0.899 | 0.217 | 0.092 | 0.021 |
| CIBERSORT | −0.049 | 0.039 | 0.210 | 0.406 | 0.176 | 0.024 |
| Inh | | | | | | |
| ICeDT | 0.093 | 0.021 | <0.001 | −0.037 | 0.063 | 0.557 |
| CIBERSORT | 0.055 | 0.035 | 0.121 | −0.005 | 0.101 | 0.965 |
| Micro | | | | | | |
| ICeDT | 0.002 | 0.039 | 0.968 | 0.107 | 0.093 | 0.253 |
| CIBERSORT | −0.014 | 0.043 | 0.744 | 0.107 | 0.097 | 0.278 |
| Oligo | | | | | | |
| ICeDT | −0.041 | 0.042 | 0.323 | −0.160 | 0.147 | 0.279 |
| CIBERSORT | −0.076 | 0.048 | 0.116 | −0.165 | 0.169 | 0.331 |
| OPC | | | | | | |
| ICeDT | 0.008 | 0.029 | 0.777 | 0.127 | 0.073 | 0.085 |
| CIBERSORT | −0.024 | 0.032 | 0.444 | 0.060 | 0.051 | 0.244 |

Table S2: Associations between relative cell type fractions and case-control status, assessed by a linear model with log ratio of cell fractions (with Exc neurons as baseline) as response, adjusted for other covariates.

| Statistic | N | Mean | St. Dev. | Min | Pctl(25) | Pctl(75) | Max |
|---|---|---|---|---|---|---|---|
| Astro.ICeDT | 527 | 0.146 | 0.046 | 0.023 | 0.121 | 0.158 | 0.416 |
| Exc.ICeDT | 527 | 0.592 | 0.078 | 0.303 | 0.551 | 0.642 | 0.751 |
| Inh.ICeDT | 527 | 0.109 | 0.032 | 0.021 | 0.087 | 0.134 | 0.191 |
| Micro.ICeDT | 527 | 0.021 | 0.012 | 0.005 | 0.014 | 0.026 | 0.087 |
| Oligo.ICeDT | 527 | 0.086 | 0.045 | 0.018 | 0.053 | 0.112 | 0.308 |
| OPC.ICeDT | 527 | 0.046 | 0.020 | 0.006 | 0.033 | 0.054 | 0.129 |
| Astro.CIBERSORT | 527 | 0.103 | 0.062 | 0.000 | 0.071 | 0.108 | 0.493 |
| Exc.CIBERSORT | 527 | 0.784 | 0.096 | 0.350 | 0.752 | 0.846 | 0.957 |
| Inh.CIBERSORT | 527 | 0.017 | 0.012 | 0.000 | 0.008 | 0.024 | 0.066 |
| Micro.CIBERSORT | 527 | 0.007 | 0.008 | 0.000 | 0.000 | 0.010 | 0.059 |
| Oligo.CIBERSORT | 527 | 0.088 | 0.056 | 0.012 | 0.049 | 0.113 | 0.405 |
| OPC.CIBERSORT | 527 | 0.001 | 0.005 | 0 | 0 | 0 | 0.50 |

Table S3: Summary of deconvolution results from ICeDT [Wilson et al., 2019] and CIBERSORT [Newman et al., 2019] for CMC data [Fromer et al., 2016], combining schizophrenia patients and controls.

| Statistic | N | Mean | St. Dev. | Min | Pctl(25) | Pctl(75) | Max |
|---|---|---|---|---|---|---|---|
| Astro.ICeDT | 85 | 0.121 | 0.056 | 0.042 | 0.089 | 0.132 | 0.456 |
| Exc.ICeDT | 85 | 0.585 | 0.086 | 0.226 | 0.552 | 0.634 | 0.710 |
| Inh.ICeDT | 85 | 0.130 | 0.036 | 0.013 | 0.109 | 0.151 | 0.227 |
| Micro.ICeDT | 85 | 0.018 | 0.014 | 0.005 | 0.011 | 0.020 | 0.110 |
| Oligo.ICeDT | 85 | 0.084 | 0.058 | 0.010 | 0.045 | 0.102 | 0.359 |
| OPC.ICeDT | 85 | 0.062 | 0.022 | 0.025 | 0.047 | 0.071 | 0.137 |
| Astro.CIBERSORT | 85 | 0.066 | 0.077 | 0.000 | 0.022 | 0.069 | 0.551 |
| Exc.CIBERSORT | 85 | 0.794 | 0.121 | 0.177 | 0.755 | 0.864 | 0.939 |
| Inh.CIBERSORT | 85 | 0.040 | 0.020 | 0.001 | 0.028 | 0.051 | 0.107 |
| Micro.CIBERSORT | 85 | 0.003 | 0.008 | 0 | 0 | 0.001 | 0 |
| Oligo.CIBERSORT | 85 | 0.097 | 0.078 | 0.007 | 0.048 | 0.117 | 0.516 |
| OPC.CIBERSORT | 85 | 0.0001 | 0.001 | 0 | 0 | 0 | 0.006 |

Table S4: Summary of deconvolution results from ICeDT [Wilson et al., 2019] and CIBERSORT [Newman et al., 2019] for the UCLA-ASD data [Parikshak et al., 2016], combining ASD patients and controls.

| Cell type | Astro | Exc | Inh | Micro | Oligo | OPC |
|---|---|---|---|---|---|---|
| CARseq | 0.051 (1.6) | 3.3e−4 (2.4) | 5.8e−7 (3.2) | 2.9e−7 (−3.2) | 3.6e−3 (2.0) | 1.9e−5 (2.8) |
| TOAST | 0.12 (−1.4) | 0.89 (0.66) | 0.14 (−1.3) | 4.1e−4 (−2.3) | 0.36 (1.1) | 0.55 (−0.91) |

Table S5: Gene Set Enrichment Analysis (GSEA) results for 328 ASD risk genes curated by SFARI. GSEA were run using the rankings of all the genes by CT-specific-DE p-values. The number is the parenthesis is the normalized enrichment score (NES). Positive NES means ASD risk genes tend to have smaller p-values, while negative NES means ASD risk genes tend to have larger p-values.

| category | pval | nDE | nCat | qval |
|---|---|---|---|---|
| EUKARYOTIC TRANSLATION ELONGATION | 1.70E-05 | 6 | 90 | 0.0098 |
| RESPONSE OF EIF2AK4 GCN2 TO AMINO ACID DEFICIENCY | 2.50E-05 | 6 | 98 | 0.0098 |
| SELENOAMINO ACID METABOLISM | 3.70E-05 | 6 | 105 | 0.0098 |
| SRP DEPENDENT COTRANSLATIONAL PROTEIN TARGETING TO MEMBRANE | 4.50E-05 | 6 | 109 | 0.0098 |
| NONSENSE MEDIATED DECAY NMD | 4.60E-05 | 6 | 113 | 0.0098 |
| EUKARYOTIC TRANSLATION INITIATION | 6.00E-05 | 6 | 117 | 0.0107 |
| CELLULAR RESPONSES TO EXTERNAL STIMULI | 1.70E-04 | 10 | 487 | 0.0236 |
| INFLUENZA INFECTION | 1.80E-04 | 6 | 149 | 0.0236 |
| INTEGRIN CELL SURFACE INTERACTIONS | 2.70E-04 | 4 | 73 | 0.0319 |
| REGULATION OF EXPRESSION OF SLITS AND ROBOS | 3.00E-04 | 6 | 160 | 0.0322 |
| EXTRACELLULAR MATRIX ORGANIZATION | 5.70E-04 | 6 | 237 | 0.0548 |
| RRNA PROCESSING | 6.20E-04 | 6 | 188 | 0.0548 |
| SIGNALING BY ROBO RECEPTORS | 8.60E-04 | 6 | 204 | 0.0708 |
| FCGR ACTIVATION | 1.30E-03 | 2 | 11 | 0.0999 |

Table S6: Over-representation of functional categories among the 65 genes with marginal DE signals (p-value $<$ 0.05) in microglia in both SCZ and ASD studies. The column of nDE is the number of DE genes within each category and the column nCat is the number of genes within that category.

#### D Supplementary Figures

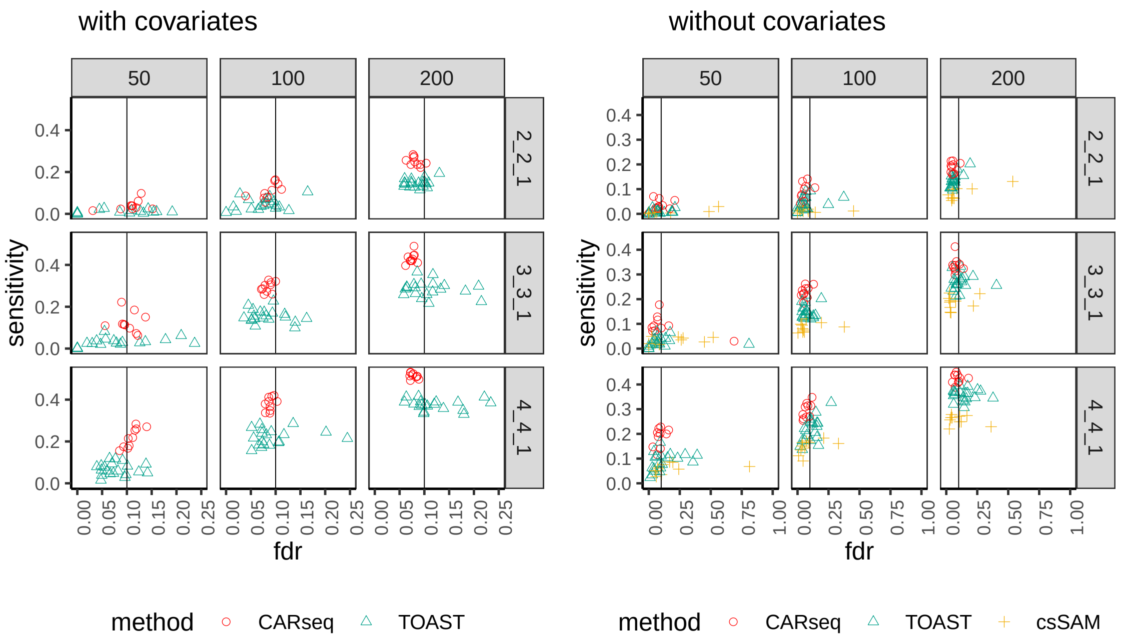

Figure S1: The FDR vs. sensitivity of several methods testing for CT-specific DE, when a confounding covariate is provided (a) or it is missing (b). There are 10 simulation replicates for each combination of total sample size with equal number of cases and controls (columns, e.g., $n = 50$ means 25 cases + 25 controls) and pattern of differential expression (rows). The notation for each pattern represents the fold changes in the major cell type, the minor cell type, and four other cell types, respectively. For example, 2_2_1 indicates that both the major cell type and the minor cell type are differentially expressed between the case and control groups by a fold change of 2 and the other four cell types are equivalently expressed between cases and controls. For each replicate, there are 2,000 genes following the prespecified pattern of differential expression and 8,000 genes with no differential expression in any of the three cell types. The vertical line indicates the intended FDR level of 0.1. Note that csSAM does not support the inclusion of covariates, and that the scales of the x axis in the two subfigures are different.

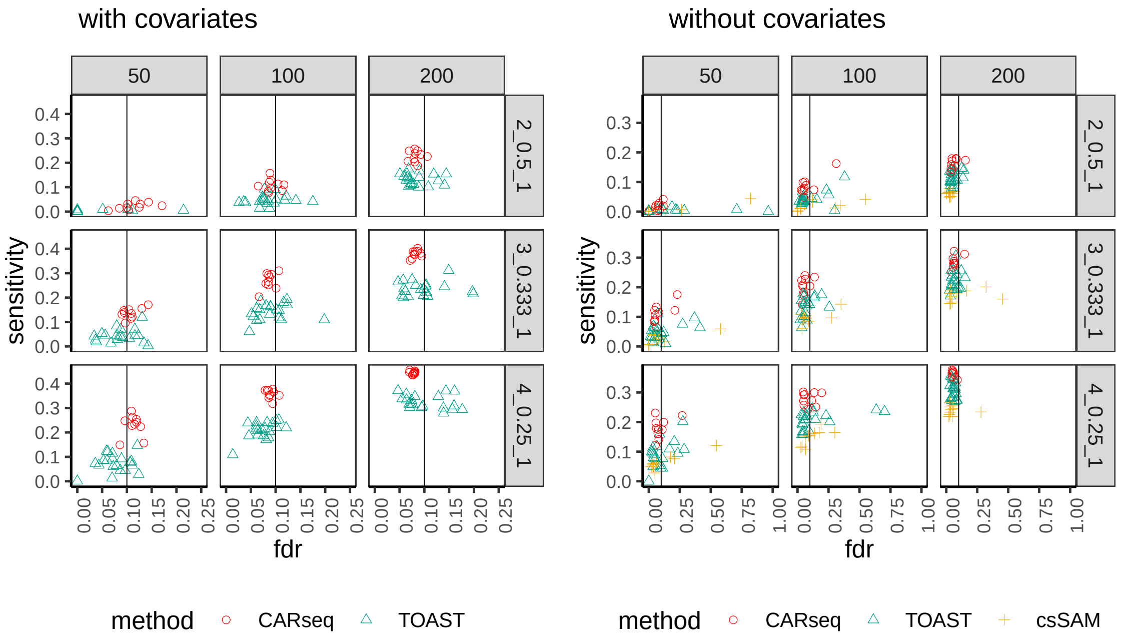

Figure S2: Similar to Figure S1, but with different patterns of differential expression where the major cell type is over-expressed and the minor cell type is under-expressed.

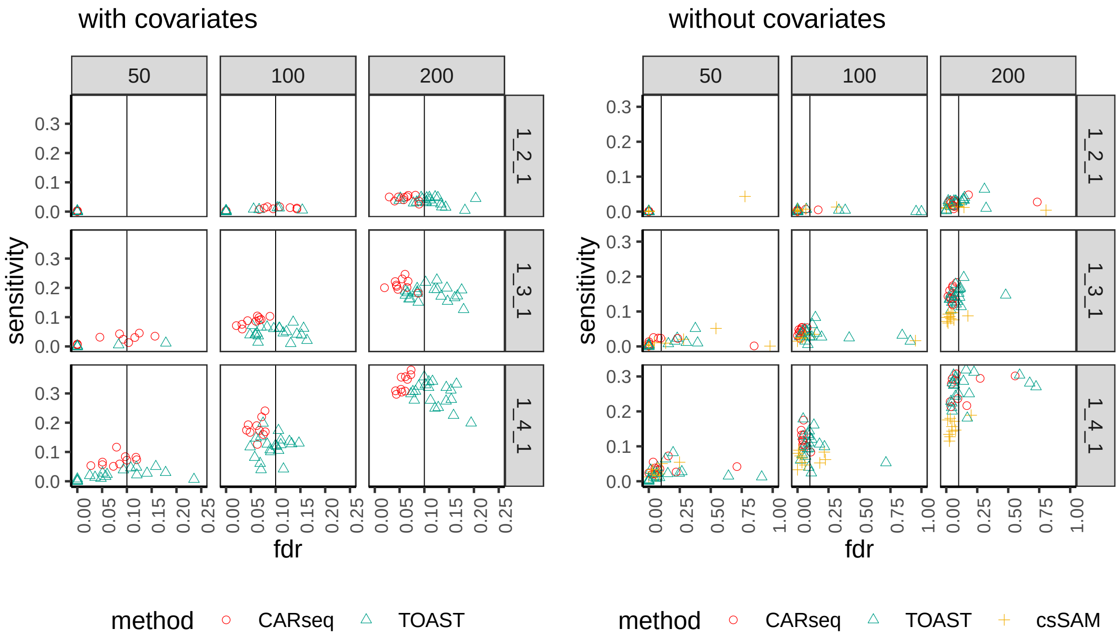

Figure S3: Similar to Figure S1, but with different patterns of differential expression where only the minor cell type is differentially expressed.

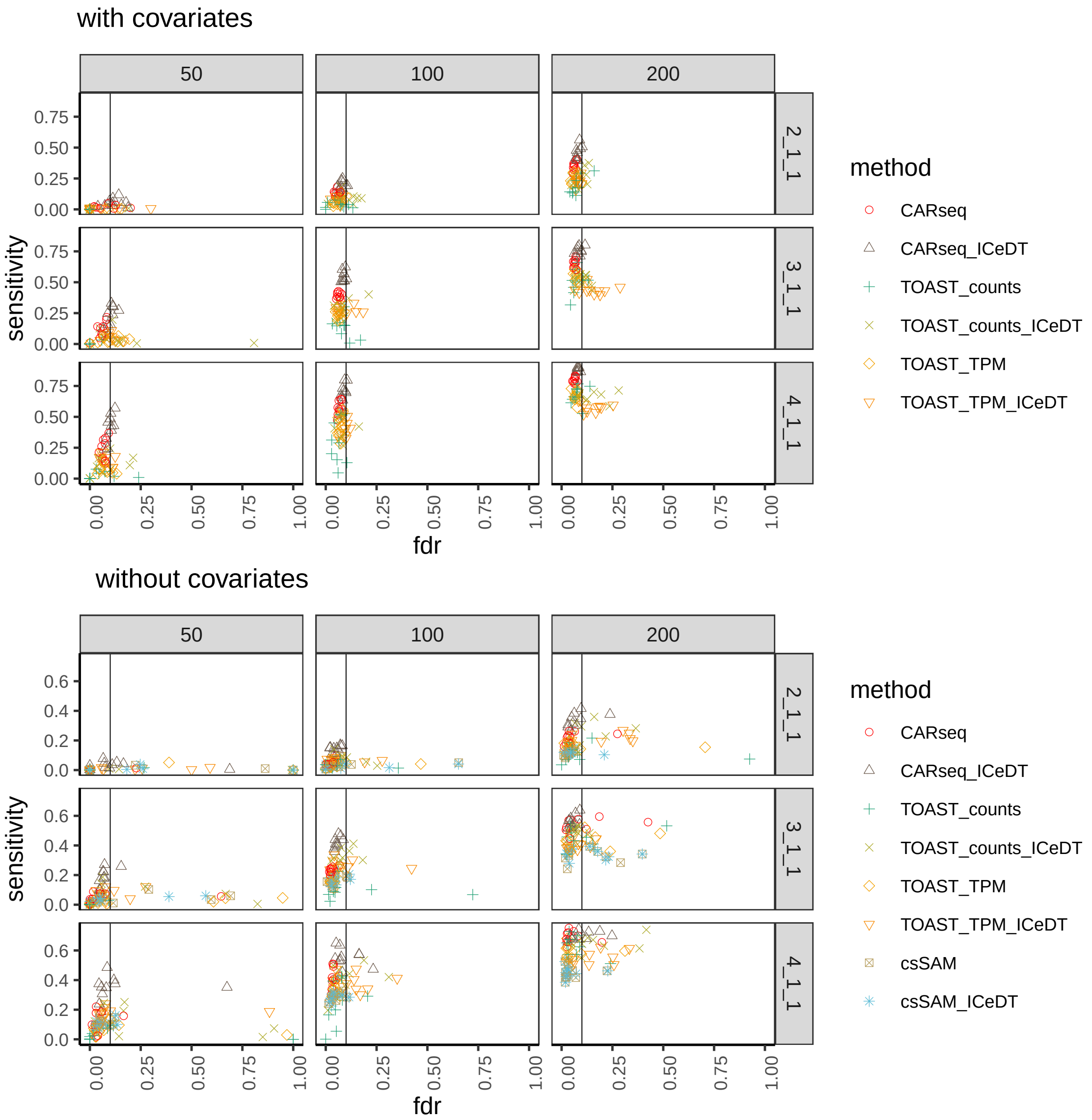

Figure S4: Results of this figure use the same simulation setup up as Figure 2 in main text, in terms sample sizes and effect sizes. The difference is that here we have one replicate per method (still with 2,000 DE genes and 8,000 non-DE genes per replicate), and more methods to compare.

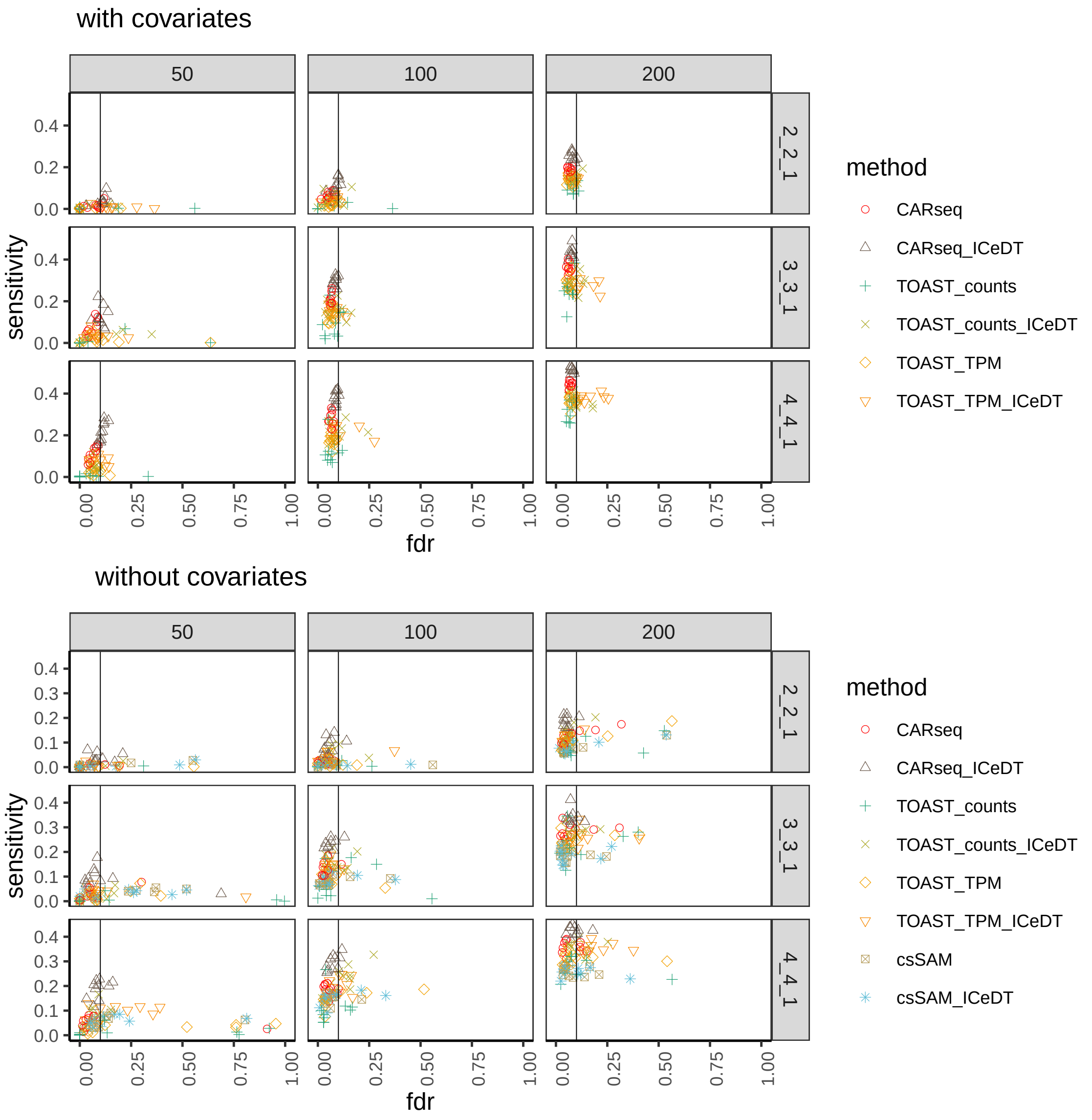

Figure S5: Similar to Figure S1 except comparing more methods with one replicate per method.

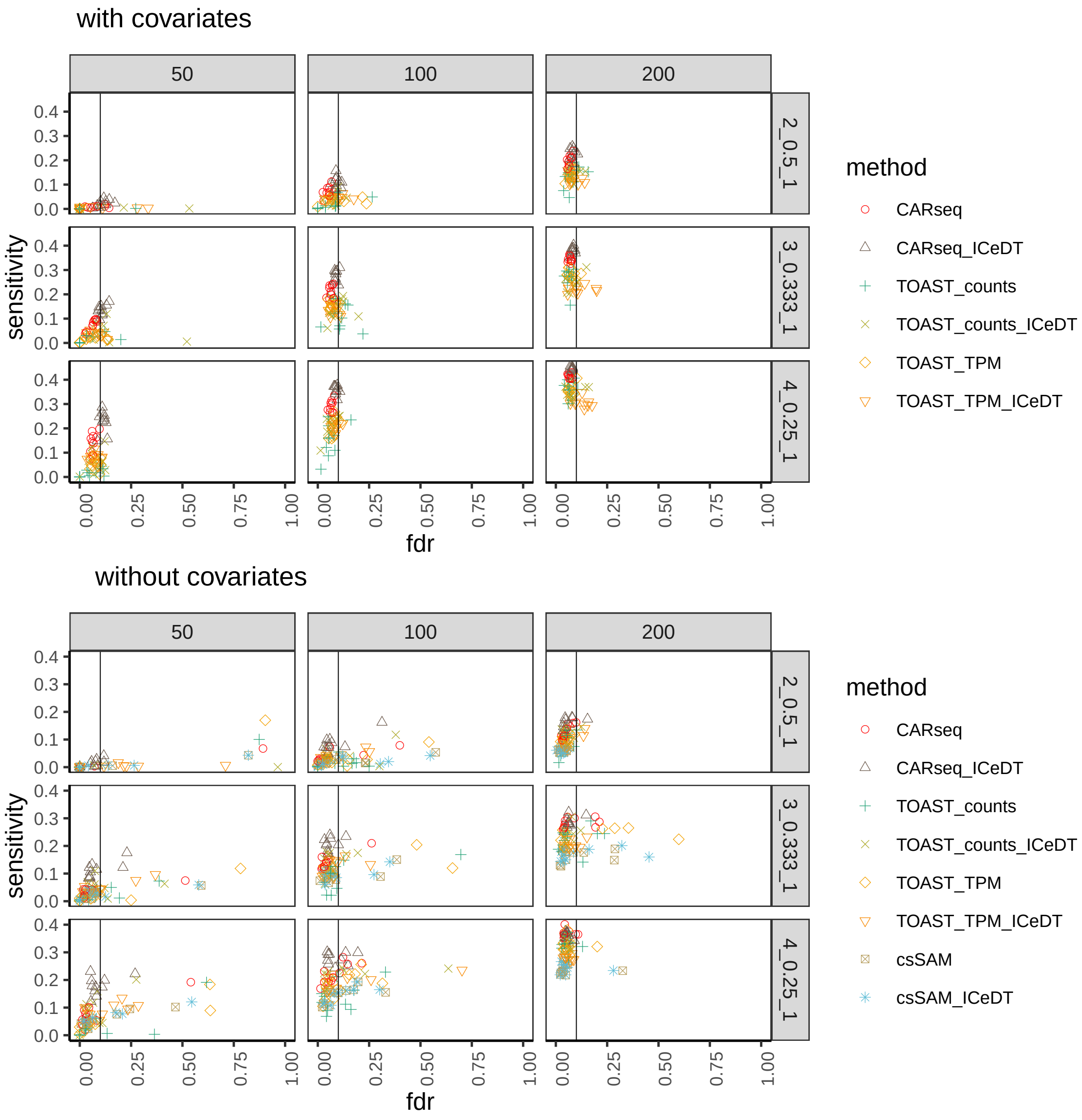

Figure S6: Similar to Figure S2 except comparing more methods with one replicate per method.

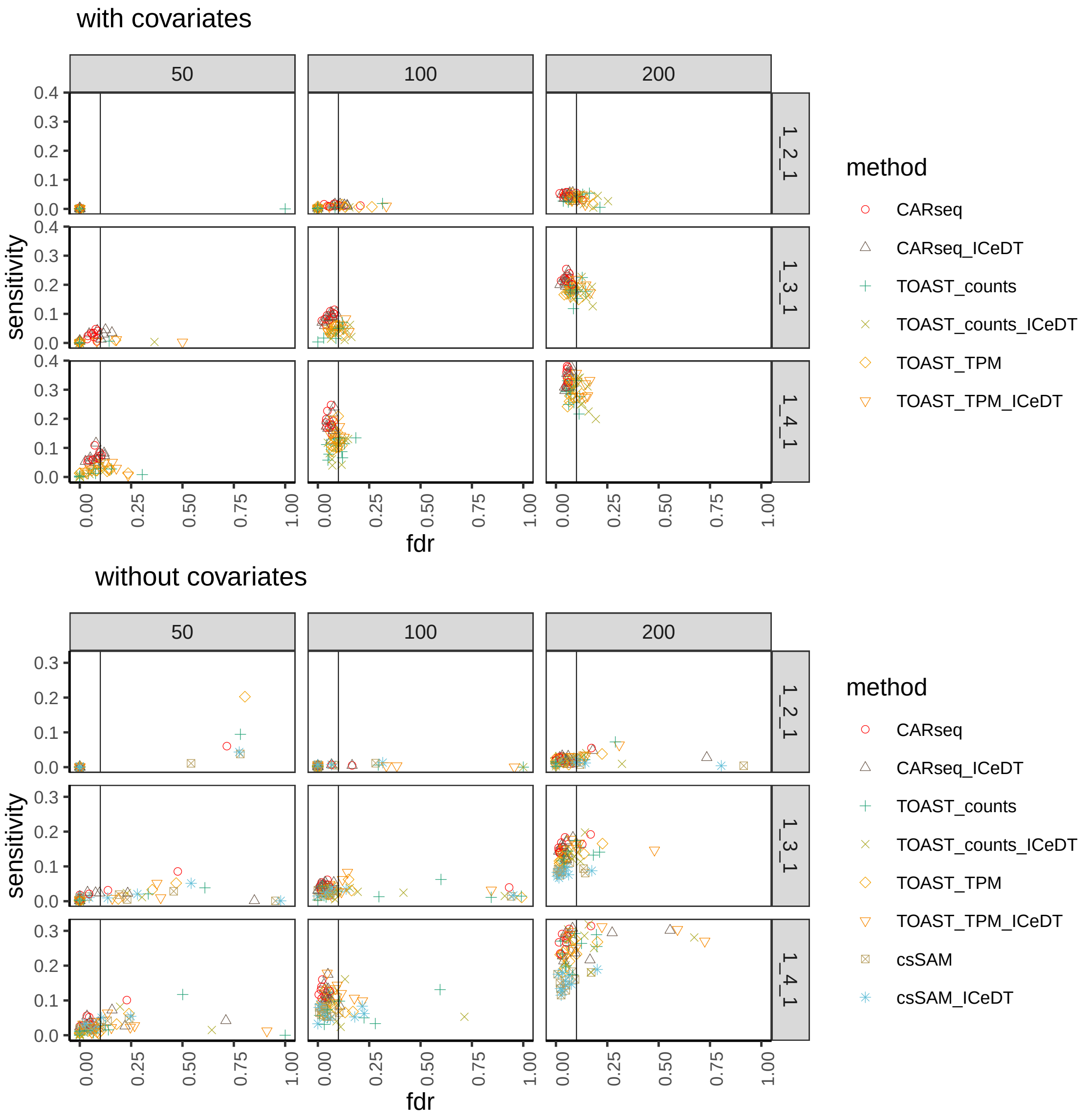

Figure S7: Similar to Figure S3 except comparing more methods with one replicate per method.

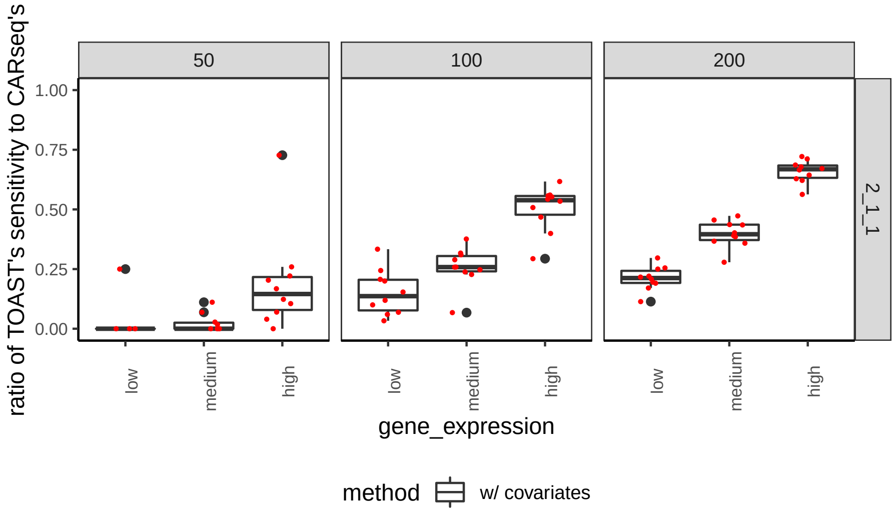

Figure S8: Ratio of the sensitivity of TOAST to the sensitivity of CARseq stratified by gene expression in the simulation setup where the major cell type is differentially expressed with fold change 2. A smaller ratio indicates that CARseq has a larger power gain over TOAST. Genes are stratified into three equally sized groups by the median read count of a gene across samples.

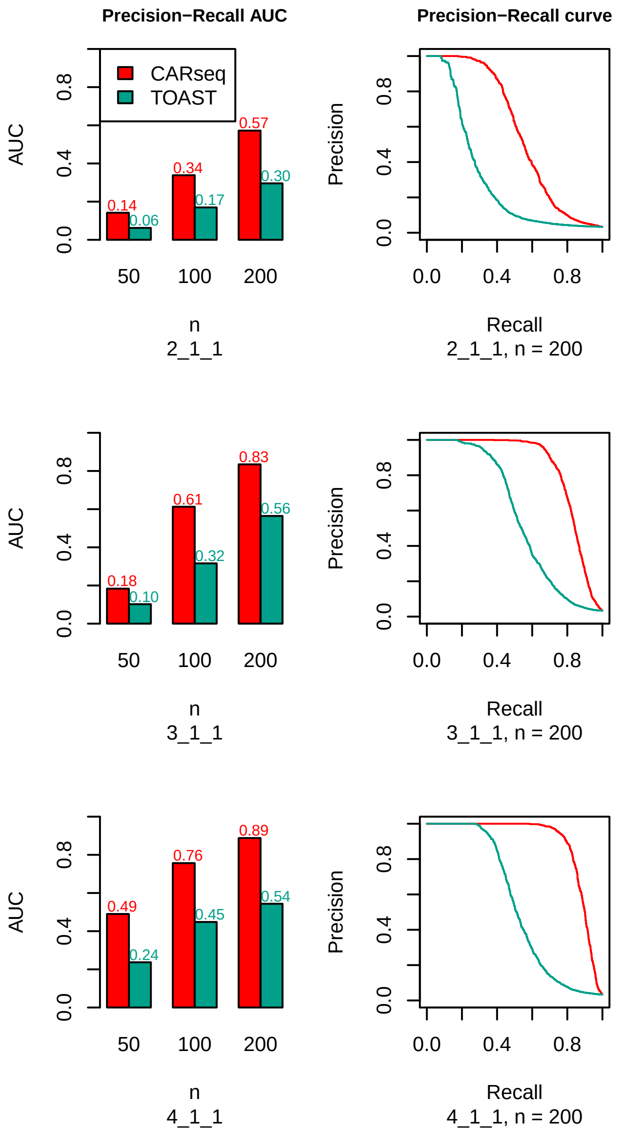

Figure S9: Precision-recall AUC (left column) and curves (right column) for CARseq and TOAST when evaluating DE for each gene and each of cell type. Each row of this figure corresponds to a DE pattern, where the three numbers separated by underscores represent the fold change in the major cell type, the minor cell type, and four other cell types. For example, 2_1_1 indicates that the major cell type is differentially expressed with fold change 2, the minor cell type, and the other four cell type are not differentially expressed. In these simulation settings, 1 of 6 cell types and 2,000 of 10,000 genes are differentially expressed, so a completely random ranking will provide a precision-recall AUC of 1/30.

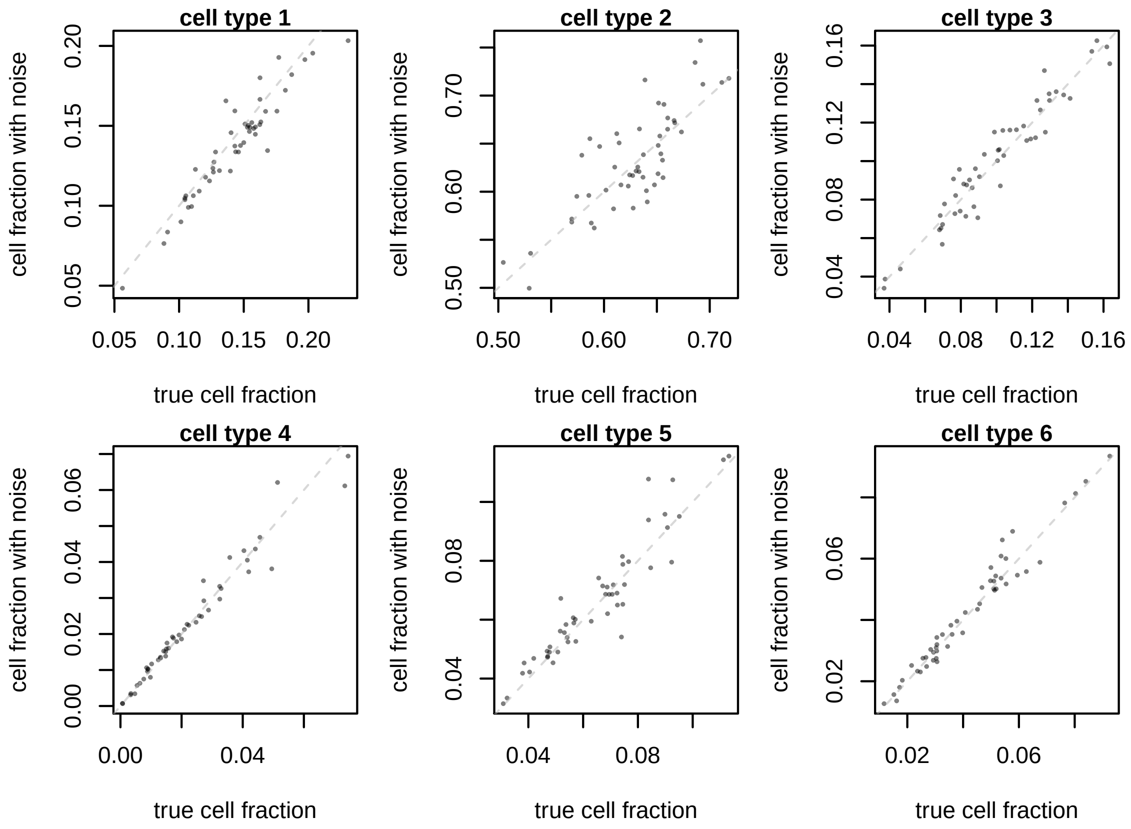

Figure S10: Simulated cell type fraction by adding noise to the true cell type fraction compared to the truth in a simulation with sample size 50 (25 cases vs. 25 controls).

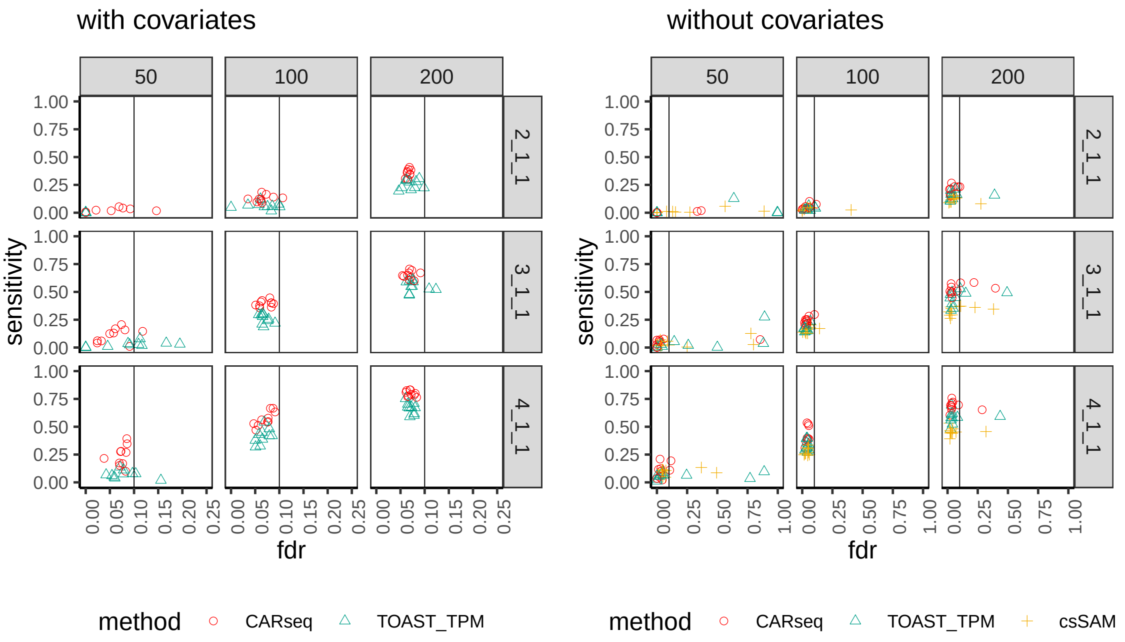

Figure S11: The FDR vs. sensitivity of several methods testing for CT-specific DE, when a confounding covariate is provided (a) or it is missing (b). A zero-centered Gaussian noise with a standard deviation of 0.1 is added to the cell fractions estimates on a logit scale. There are 10 simulation replicates for each combination of total sample size with equal number of cases and controls (columns, e.g., $n = 50$ means 25 cases + 25 controls) and pattern of differential expression (rows). The notation for each pattern represents the fold changes in the major cell type, the minor cell type, and four other cell types, respectively. For example, 2_1_1 indicates that both the major cell type is differentially expressed between the case and control groups by a fold change of 2 and the minor cell type and the other four cell types are equivalently expressed between cases and controls. For each replicate, there are 2,000 genes following the pre-specified pattern of differential expression and 8,000 genes with no differential expression in any of the three cell types. The vertical line indicates the intended FDR level of 0.1. Note that csSAM does not support the inclusion of covariates, and that the scales of the x axis in the two subfigures are different.

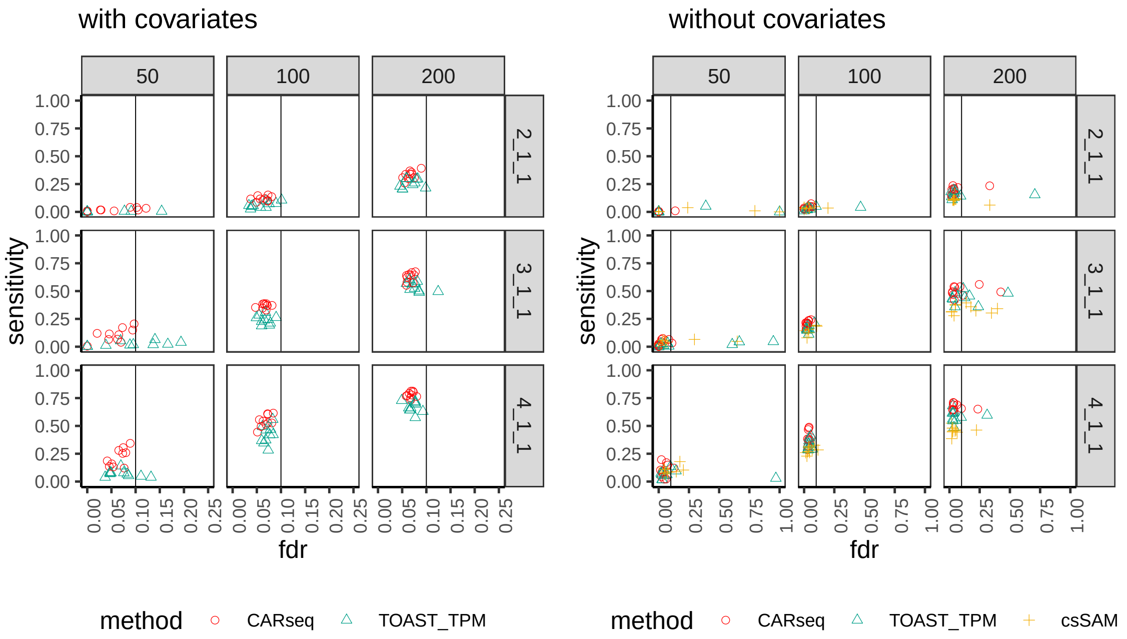

Figure S12: The FDR vs. sensitivity of several methods testing for CT-specific DE, when a confounding covariate is provided (a) or it is missing (b). The cell size factor is misspecified; we intentionally apply a list of wrong size factors (1.2, 1, 1) in the inference and testing instead of the true size factors (1, 1, 1) used in data simulation. There are 10 simulation replicates for each combination of total sample size with equal number of cases and controls (columns, e.g., $n = 50$ means 25 cases + 25 controls) and pattern of differential expression (rows). The notation for each pattern represents the fold changes in the major cell type, the minor cell type, and four other cell types, respectively. For example, 2_1_1 indicates that both the major cell type is differentially expressed between the case and control groups by a fold change of 2 and the minor cell type and the other four cell types are equivalently expressed between cases and controls. For each replicate, there are 2,000 genes following the pre-specified pattern of differential expression and 8,000 genes with no differential expression in any of the three cell types. The vertical line indicates the intended FDR level of 0.1. Note that csSAM does not support the inclusion of covariates, and that the scales of the x axis in the two subfigures are different.

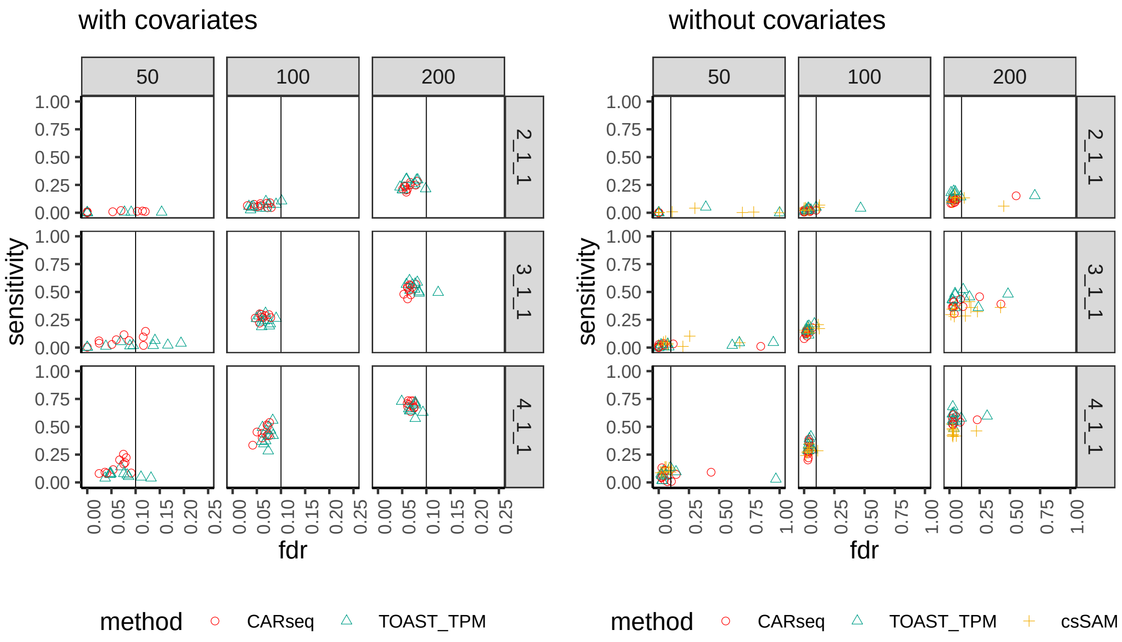

Figure S13: Similar to Figure S12 except the cell size factors are extremely misspecified to be 2, 1, 1.

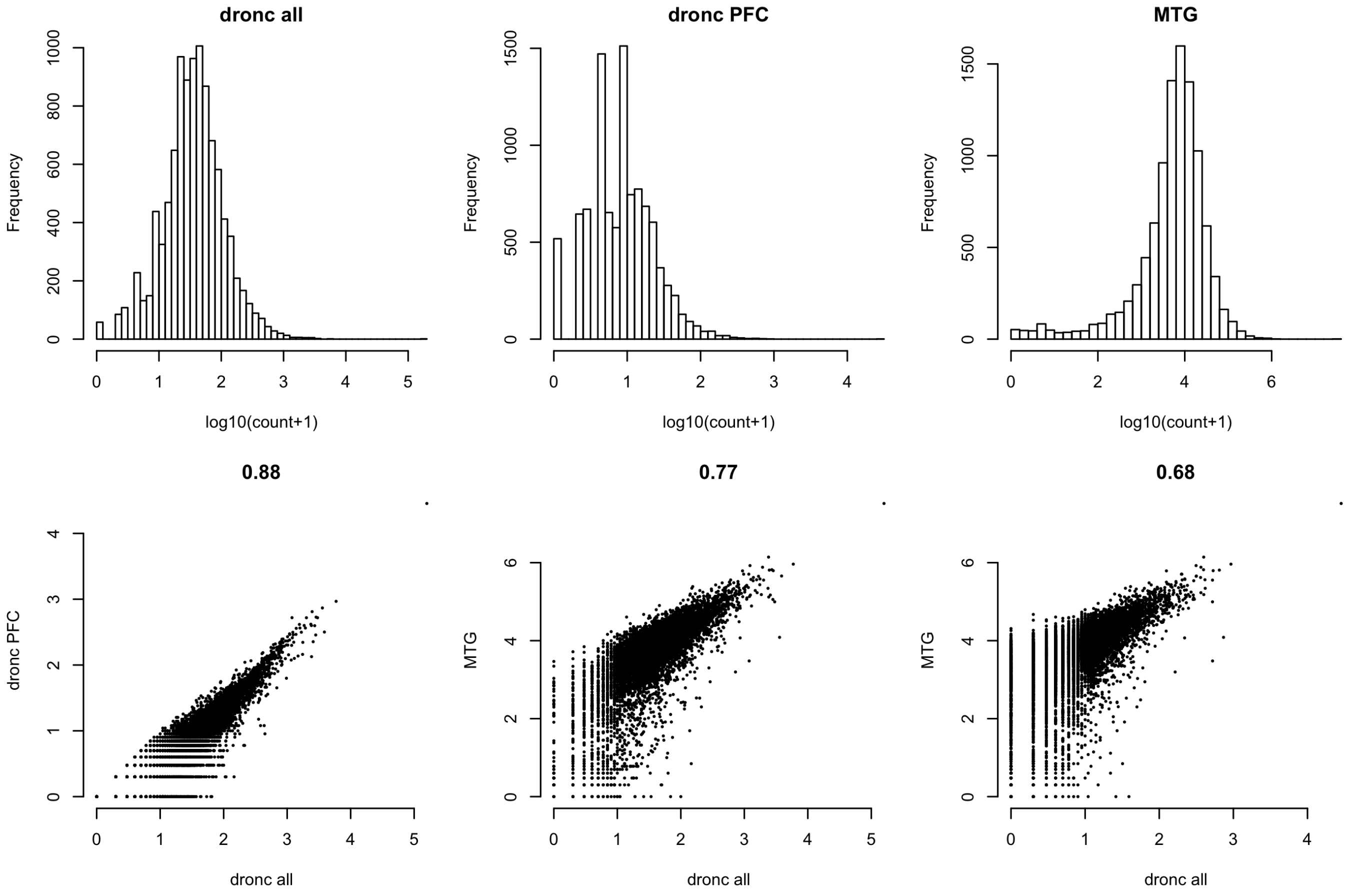

Figure S14: Compare astrocyte-specific gene expression of 10,151 genes derived from DroNC and MTG data. The top panel shows the distribution of gene expression in log transformed raw counts. Note that the counts are summation across 1584, 339, and 287 cells for all DroNC data, DroNC PFC only, and MTG, respectively. The lower panel shows pair-wise scatter plots.

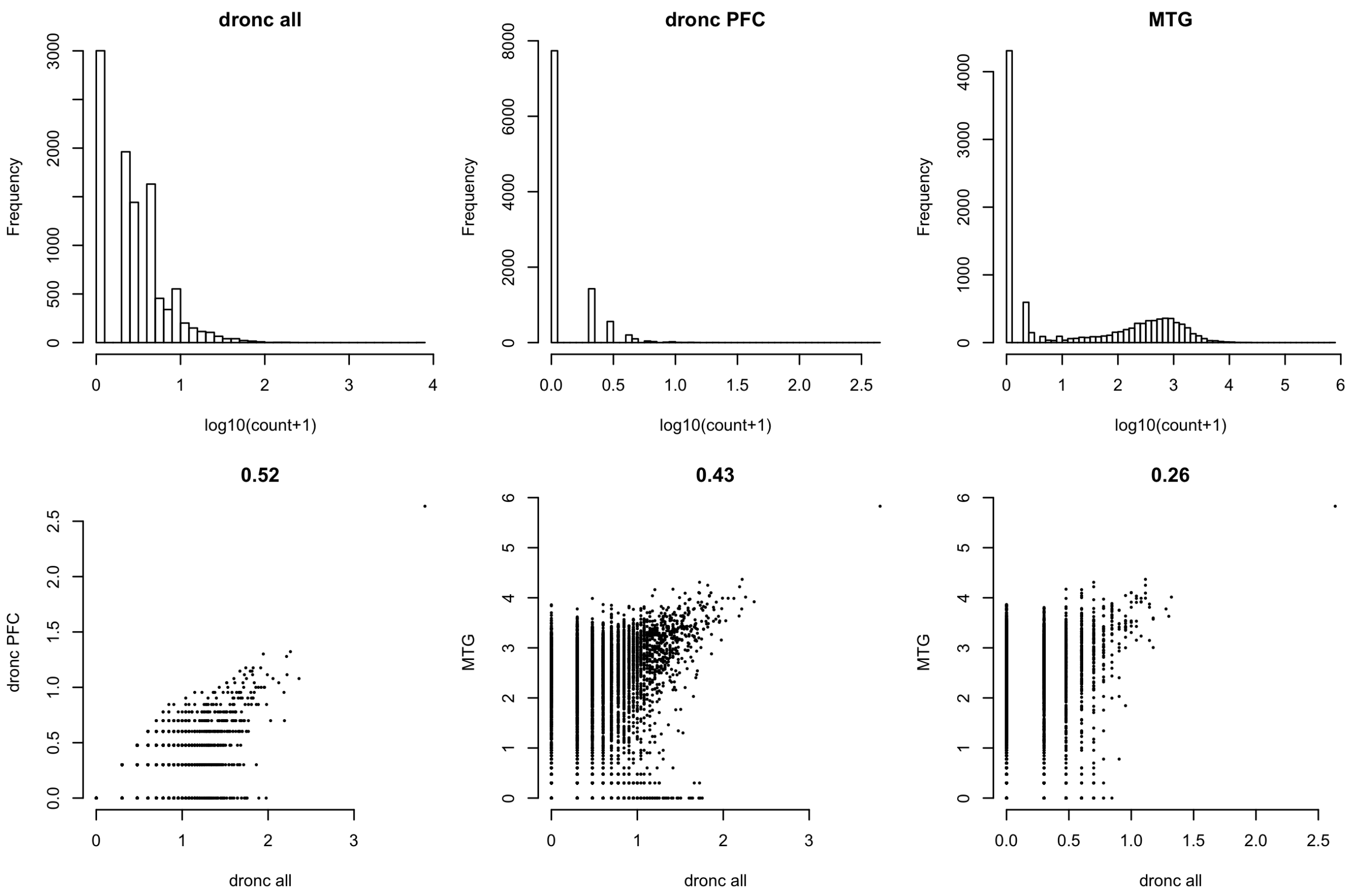

Figure S15: Compare endothelial-specific gene expression of 10,151 genes derived from DroNC and MTG data. The top panel shows the distribution of gene expression in log transformed raw counts. Note that the counts are summation across 68, 8, and 8 cells for all DroNC data, DroNC PFC only, and MTG, respectively. The lower panel shows pair-wise scatter plots.

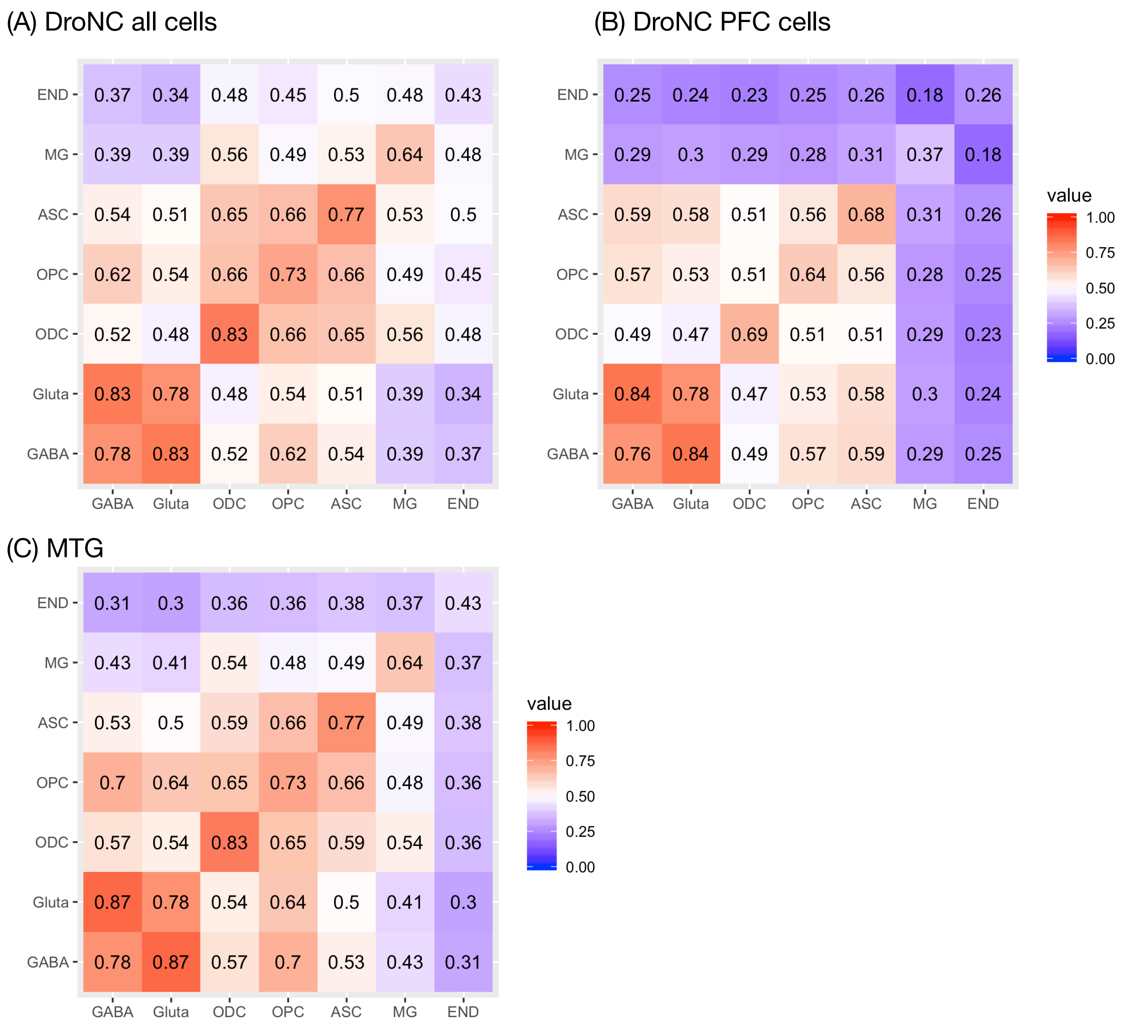

Figure S16: Correlations of genome-wide gene expression (10,151 genes) within each dataset, while the diagonals of the three panels were replaced by the correlation of (A) DroNC (all cells) vs. MTG (B) DroNC (PFC cells) vs. MTG, and (C) DroNC (all cells) vs. MTG. GABA: GABAergic interneurons or inhibitory neurons; Gluta: glutamatergic neurons or excitatory neurons; ODC: oligodendrocytes; OPC: oligodendrocyte precursor cells; ASC: astrocytes; MG: microglia; END: endothelial cells.

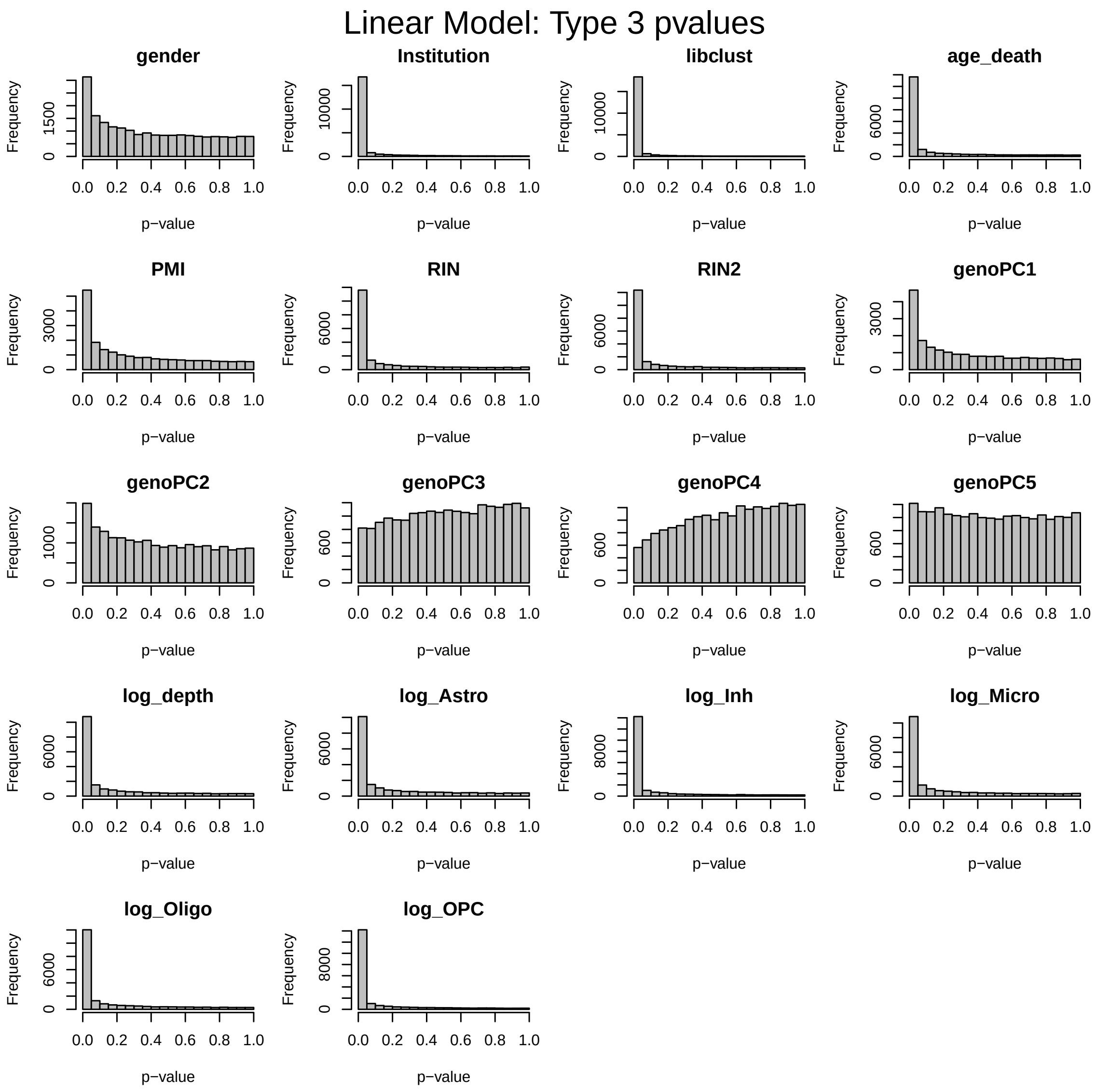

Figure S17: The p-values for each covariate vs. genome-wide gene expression for the schizophrenia study, assessed by a linear model for log-transformed gene expression. Cell type proportions were included as log ratios, e.g., `log_Astro` is log(Astro proportion/Excitatory neuron proportion).

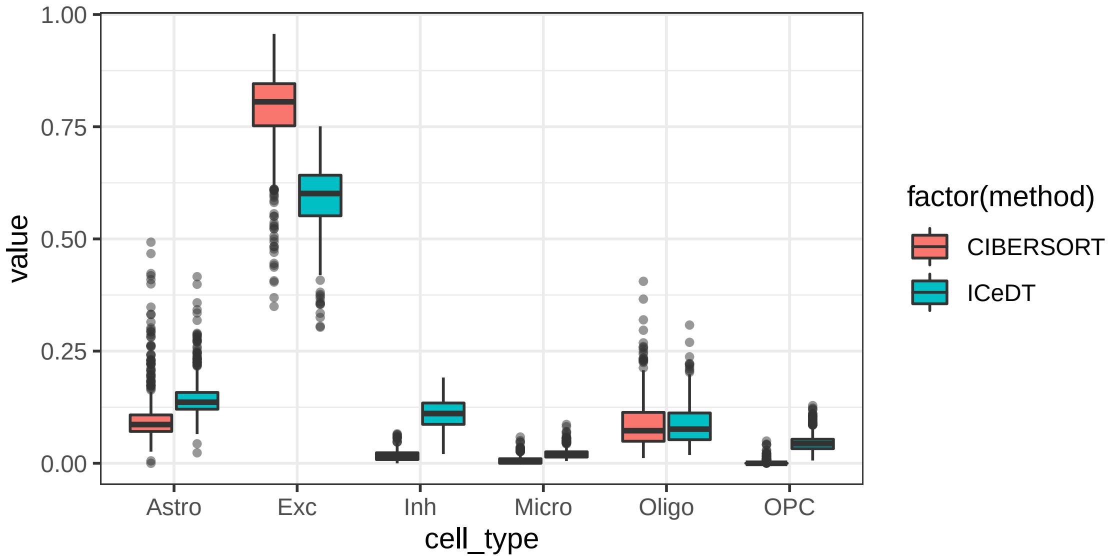

Figure S18: Box plots of cell type fraction estimates for CMC data.

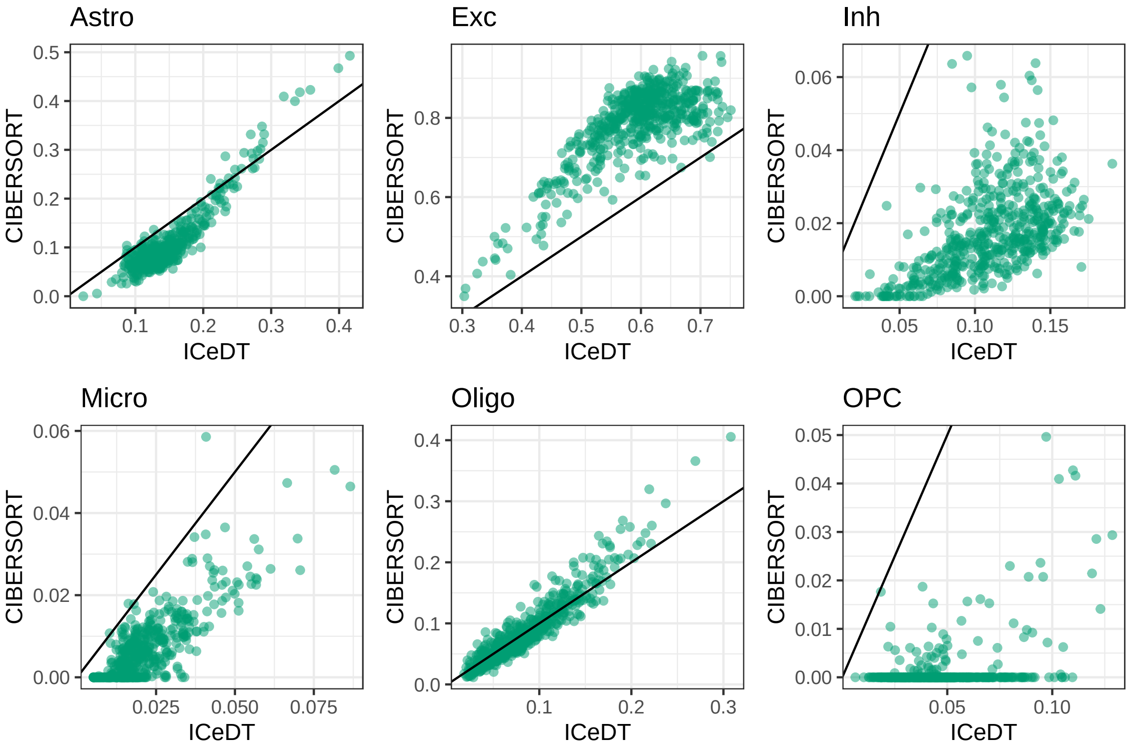

Figure S19: Scatter plots of cell type fraction estimates for CMC data.

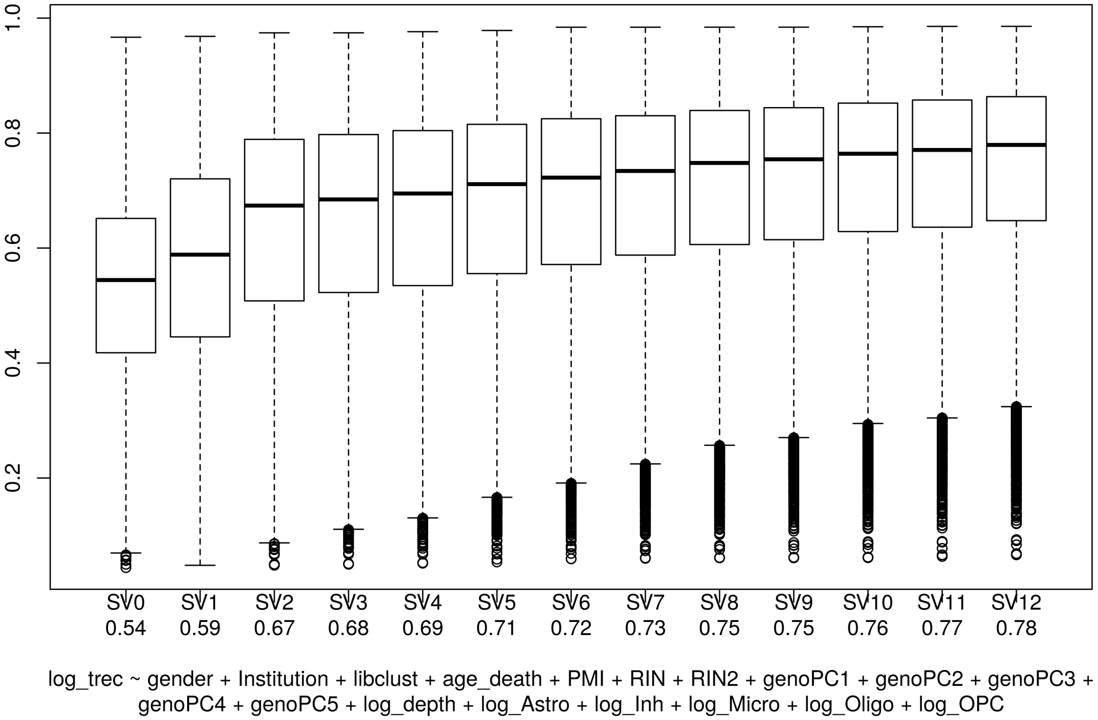

Figure S20: $R^2$ explained by increasing number of surrogate variables for CMC data [Fromer et al., 2016].

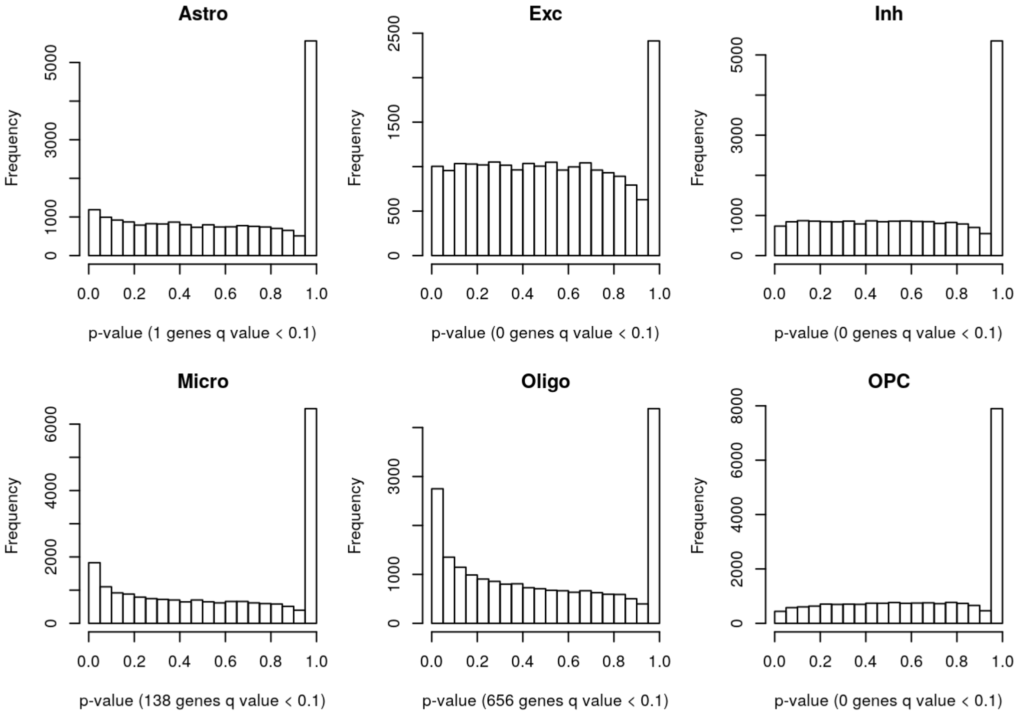

Figure S21: CARseq p-value distribution in the SCZ study.

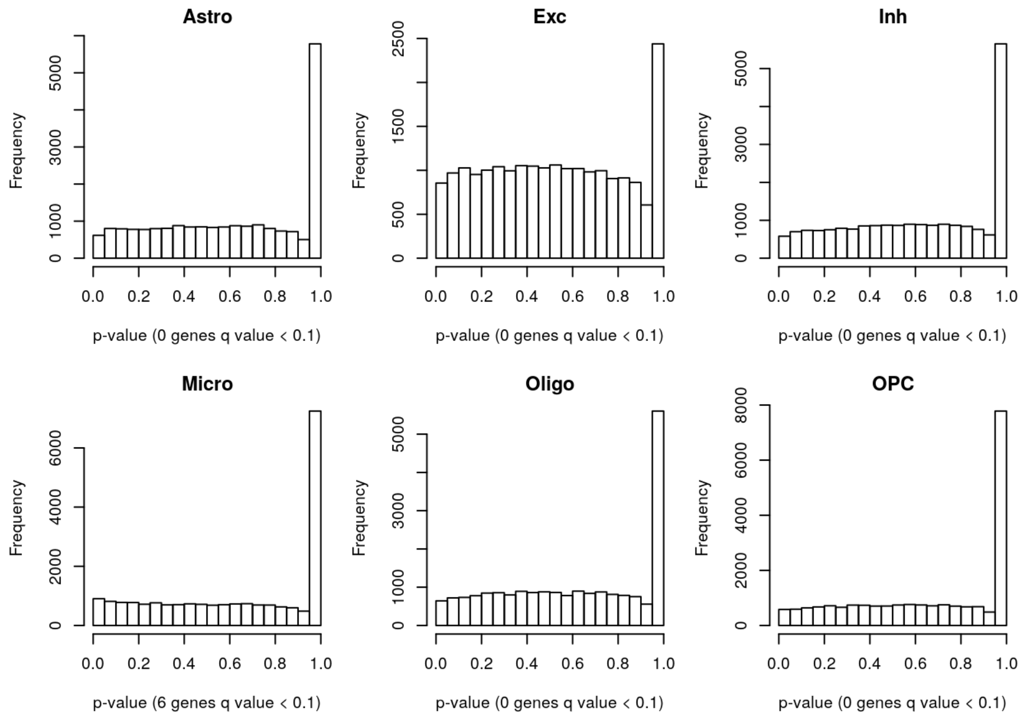

Figure S22: CARseq p-value distribution in the SCZ study where the case-control label has been permuted to reflect the null distribution.

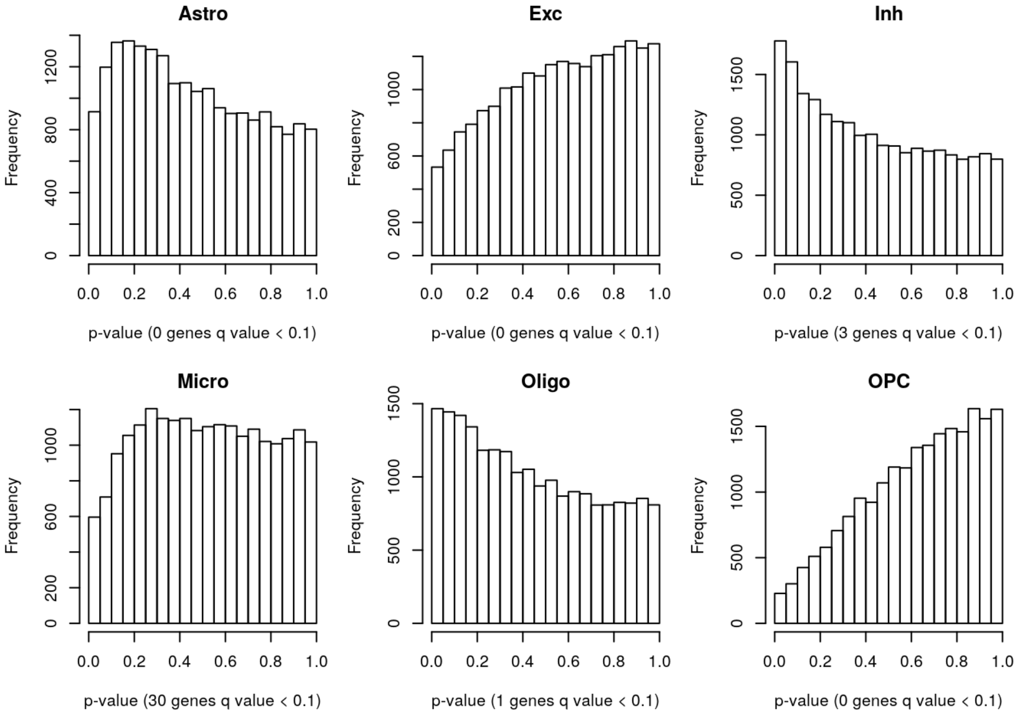

Figure S23: TOAST p-value distribution in the SCZ study.

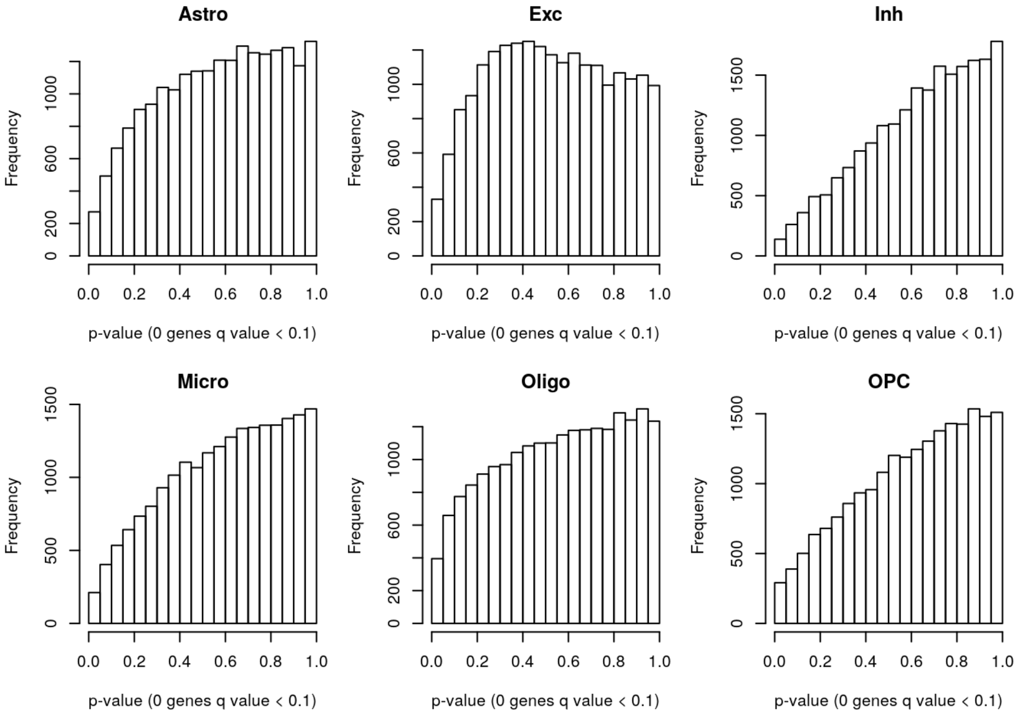

Figure S24: TOAST p-value distribution in the SCZ study where the case-control label has been permuted to reflect the null distribution.

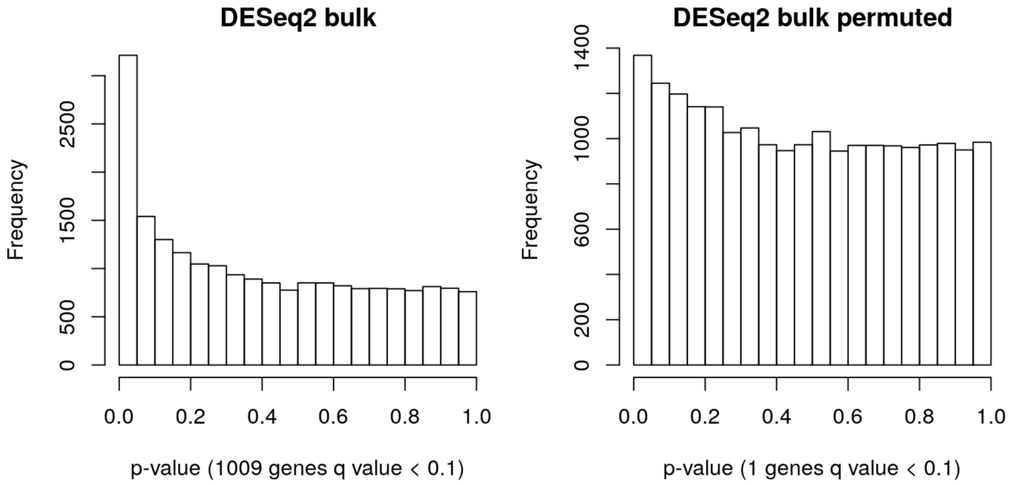

Figure S25: DESeq2 p-value distribution in the SCZ study where the case-control label is either unpermuted or permuted.

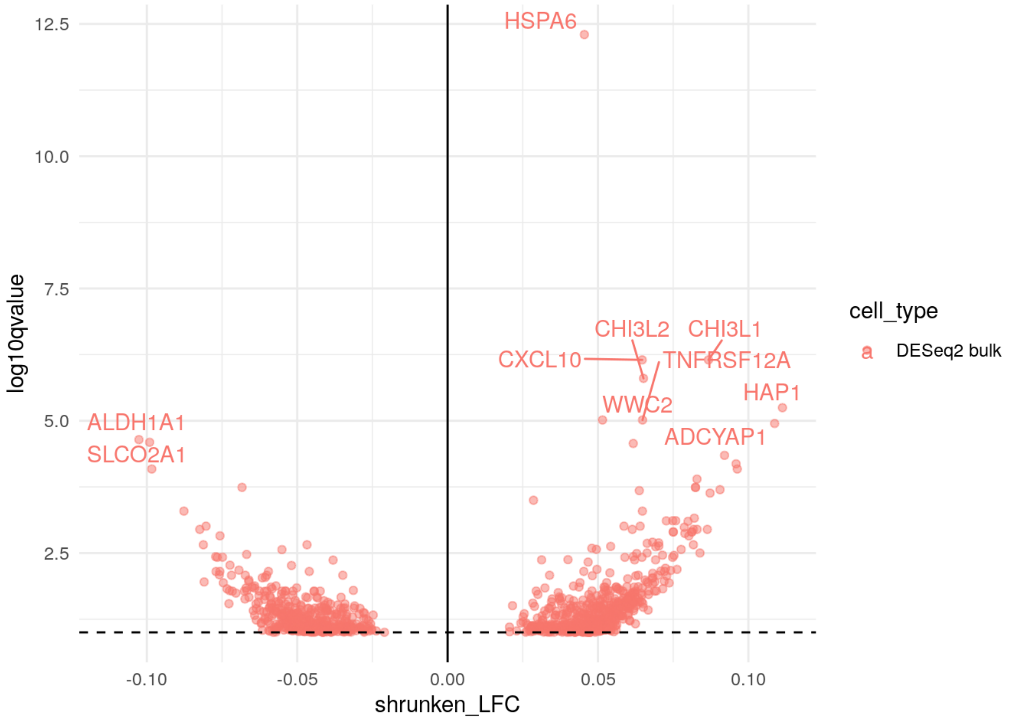

Figure S26: DESeq2 volcano plot in the SCZ study.

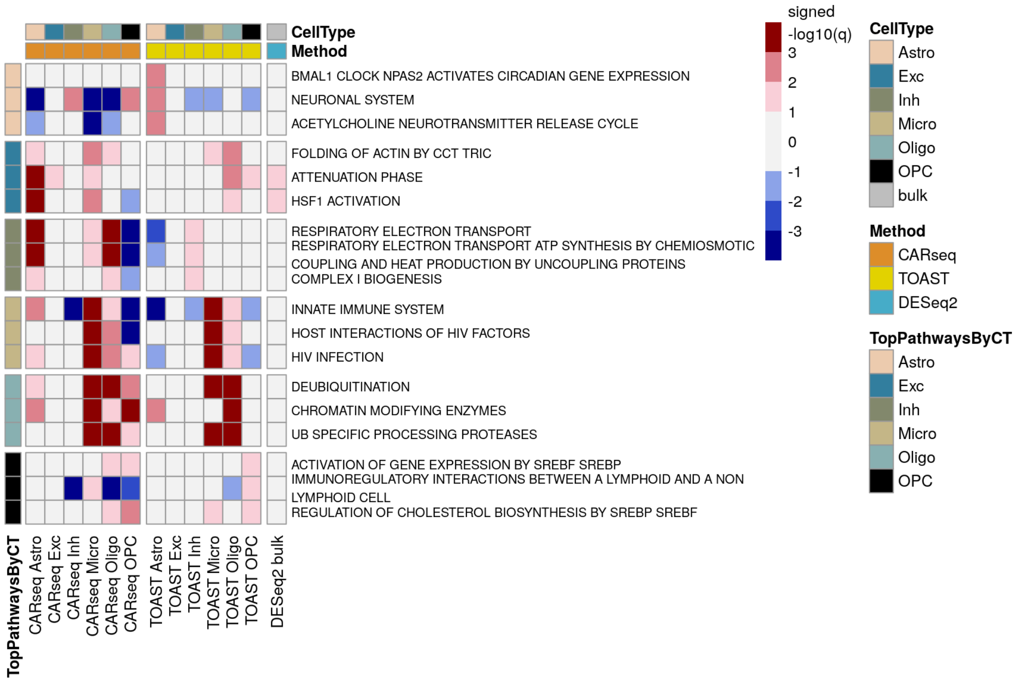

Figure S27: REACTOME GSEA ranked by TOAST in the SCZ study.

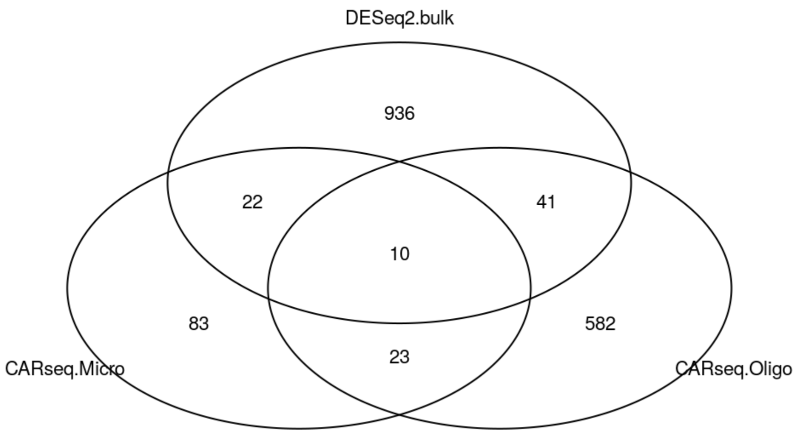

Figure S28: Venn plot of DEGs ($q$-value $< 0.1$) in the SCZ study.

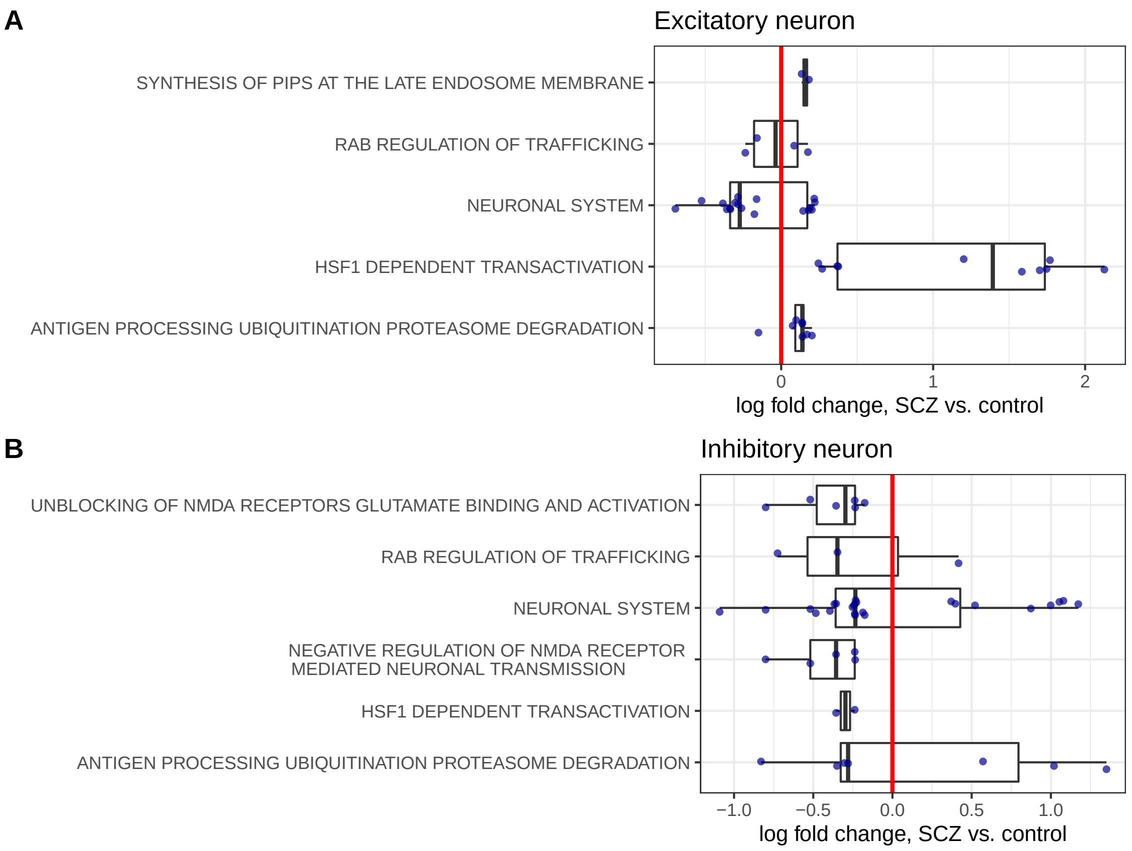

Figure S29: Shrunken log fold change estimates (SCZ vs. controls) for genes belonging to some REACTOME pathways. Panel A includes all the pathways related with NDMA. Panels B and C include the pathway identified by GSEA in excitatory/inhibitory neurons, from either SCZ or ASD studies. Only the genes with CT-specific-DE p-value (comparing SCZ vs. controls) smaller than 0.05 are shown.

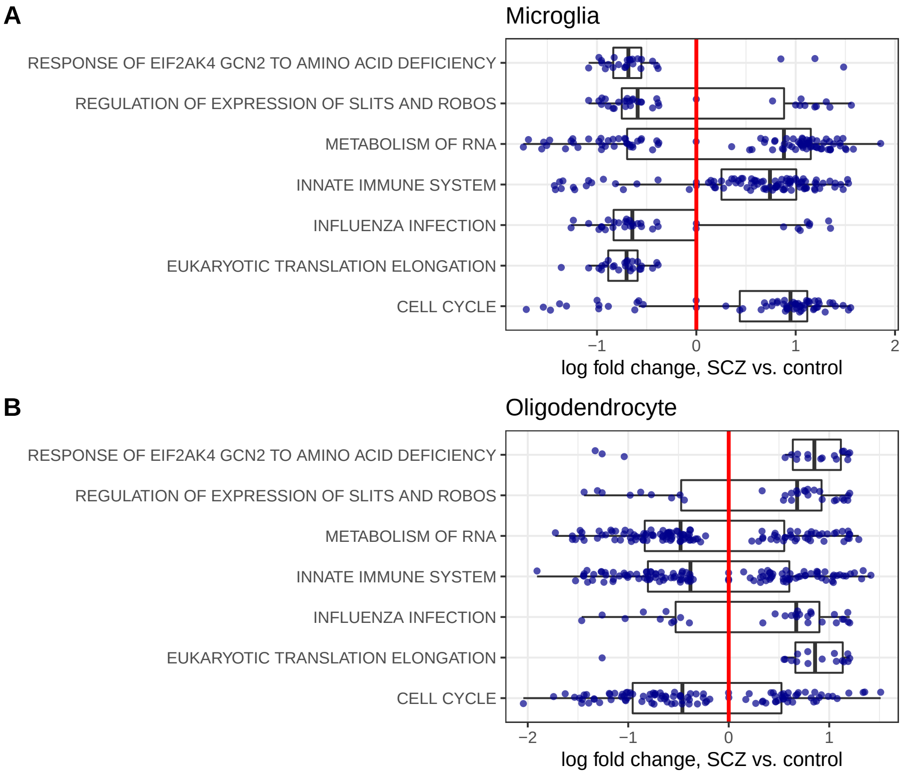

Figure S30: Shrunken log fold change estimates (SCZ vs. controls) for genes belonging to the REACTOME pathways identified by GSEA in microglias or oligodendrocytes, from either SCZ or ASD studies. Only the genes with CT-specific-DE p-value (comparing SCZ vs. controls) smaller than 0.05 are shown.

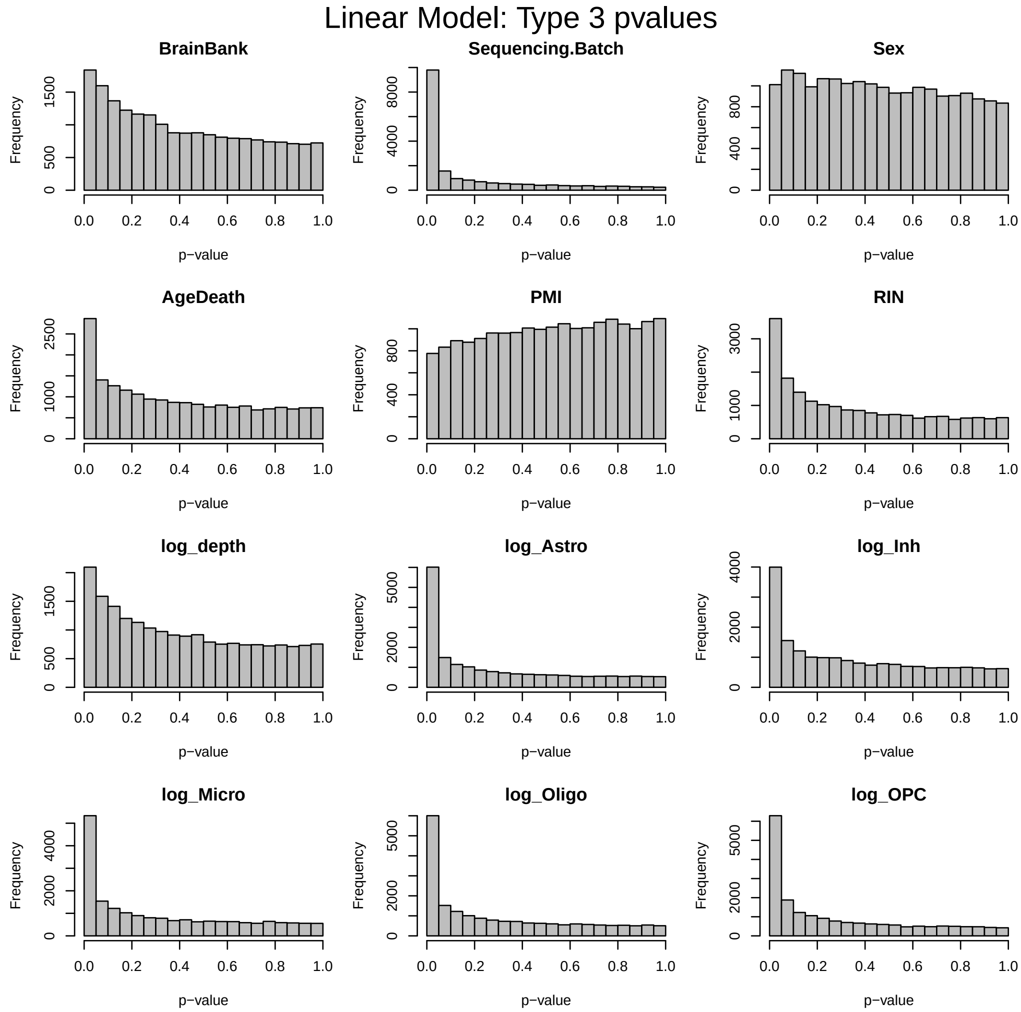

Figure S31: The p-values for each covariate vs. genome-wide gene expression for the ASD study, assessed by a linear model for log-transformed gene expression. Cell type proportions were included as log ratios, e.g., `log_Astro` is log(Astro proportion/Excitatory neuron proportion). The analyses for this figure were done using 82 out of 85 samples with non-missing PMI values.

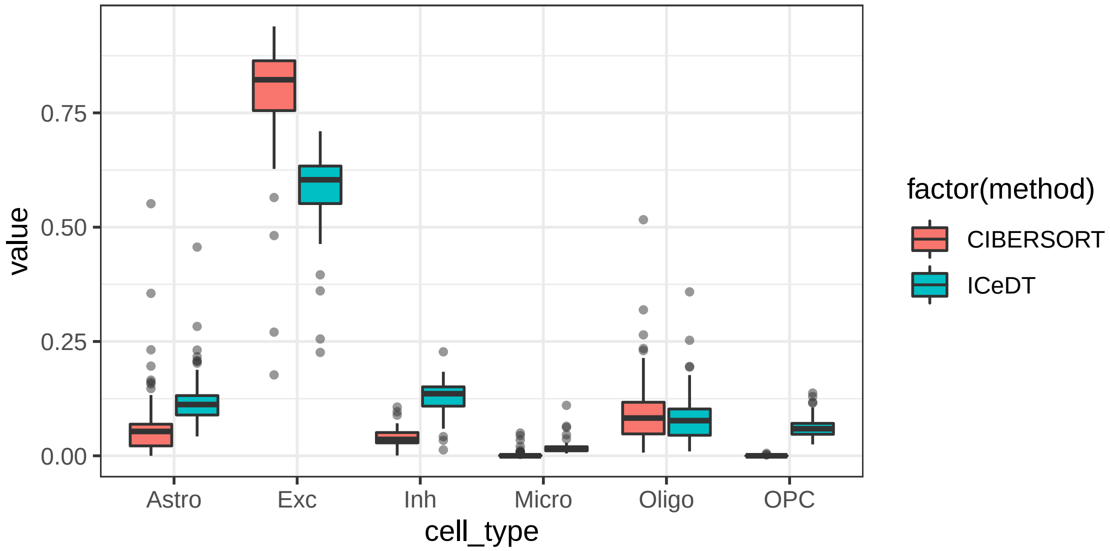

Figure S32: Box plots of cell type fraction estimates for UCLA-ASD data.

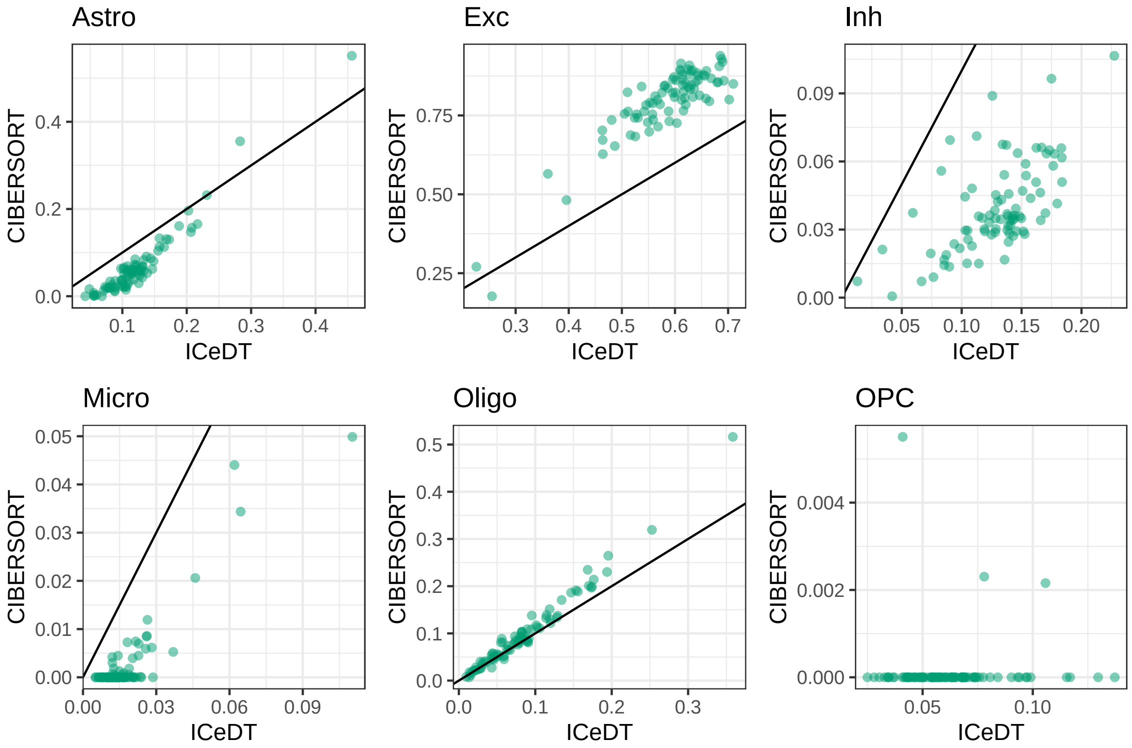

Figure S33: Scatter plots of cell type fraction estimates for UCLA-ASD data.

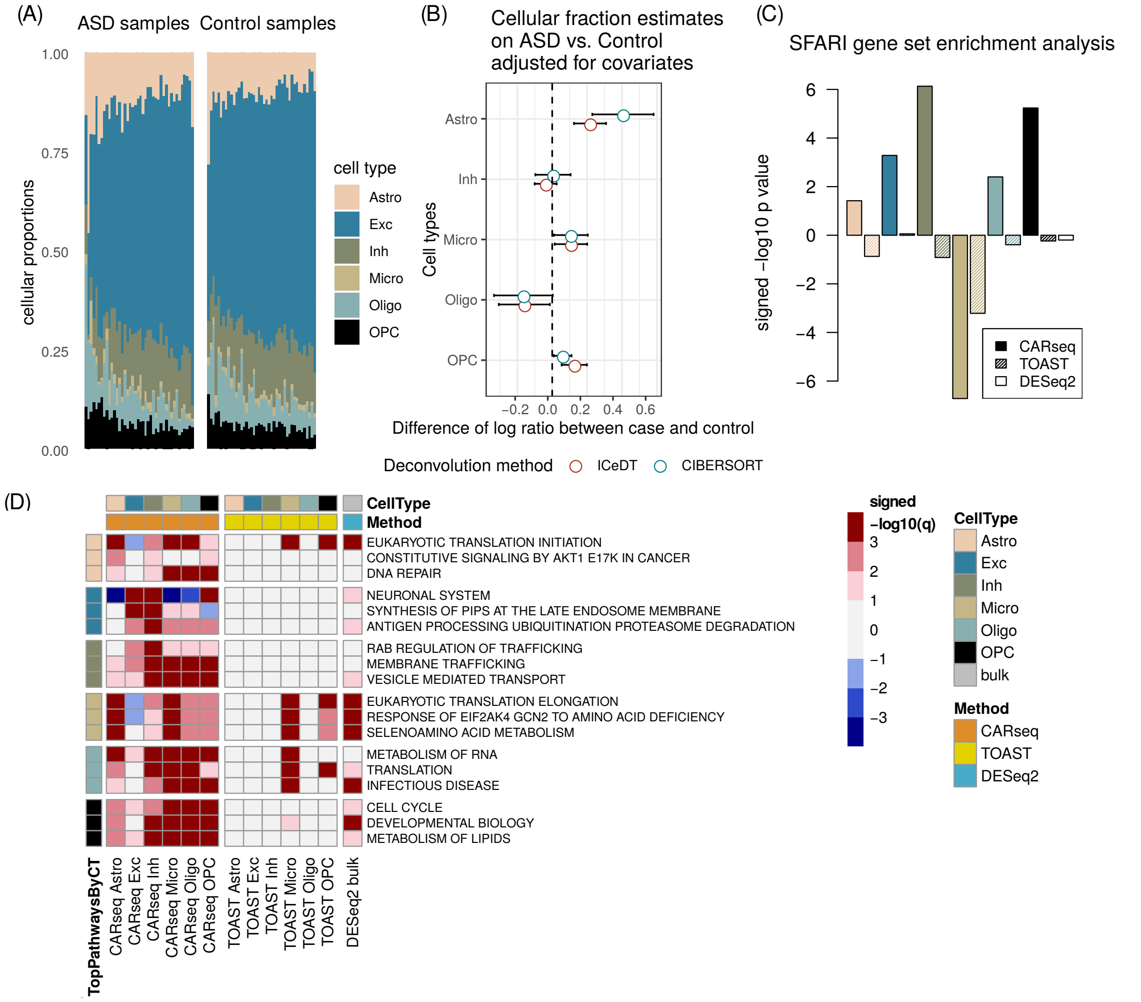

Figure S34: CARseq on the autism spectrum disorder (ASD) bulk expression data. (A) Estimated cell fractions by ICeD-T sorted by increasing fractions of excitatory neurons. (B) The effect size of case-control status on relative cell fractions against excitatory neurons (log ratio of the cell type of interest vs. excitatory neuron). The standard errors are denoted by bars. (C) Gene set enrichment analysis results in -log10 p value with the sign of normalized enrichment score (NES) of the SFARI gene set, a curated list of 328 autism risk genes. (D) Gene set enrichment analysis results on REACTOME pathways. Three top pathways were shown for each cell type, ranked by -log10 q value with the sign of normalized enrichment score (NES). Positive NES indicates enrichment of genes with small p-values.

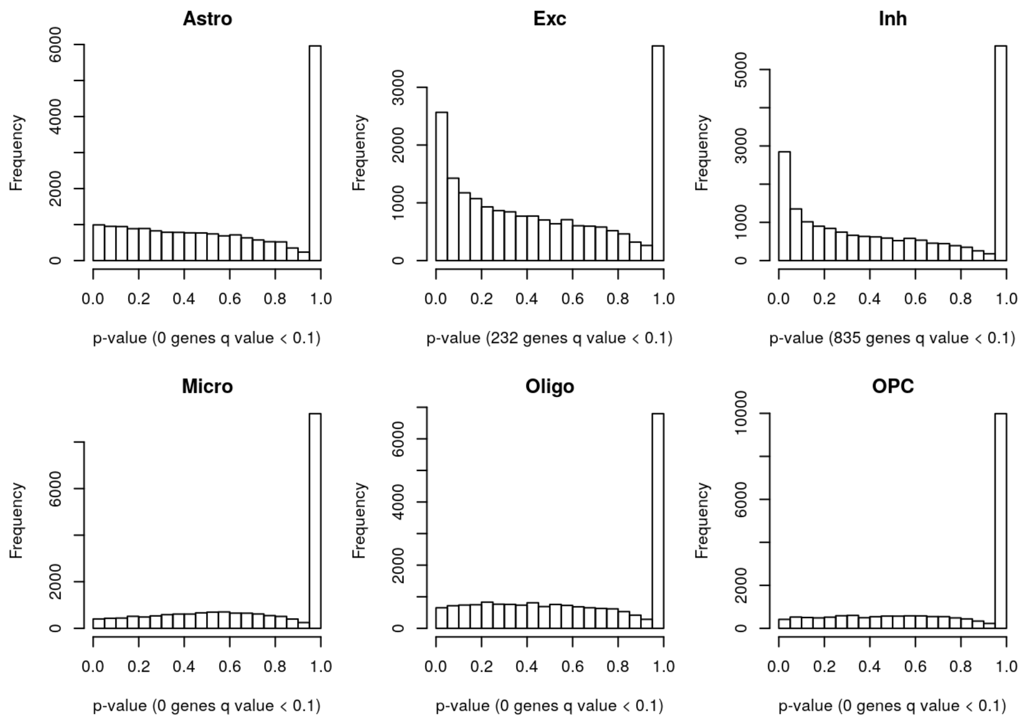

Figure S35: CARseq p-value distribution in the ASD study.

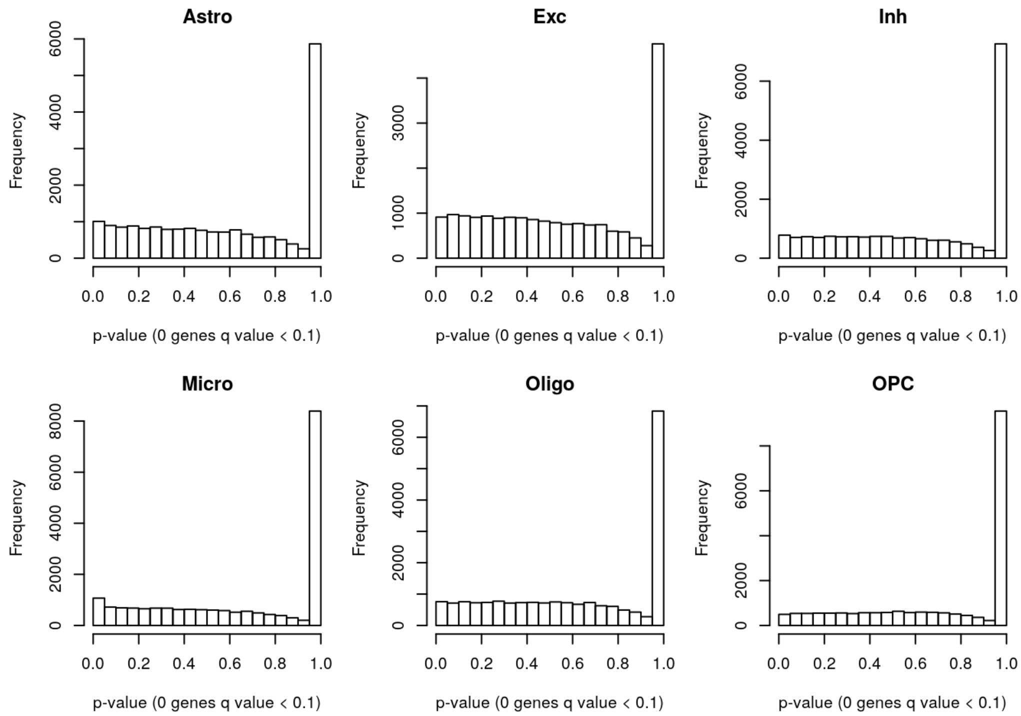

Figure S36: CARseq p-value distribution in the ASD study where the case-control label has been permuted to reflect the null distribution.

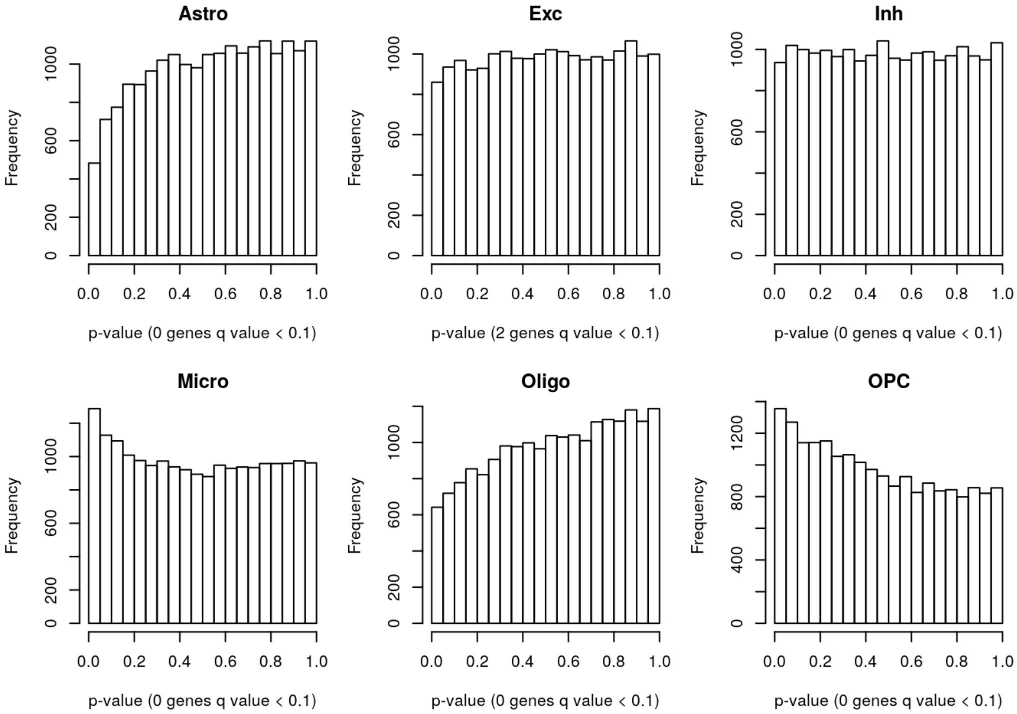

Figure S37: TOAST p-value distribution in the ASD study.

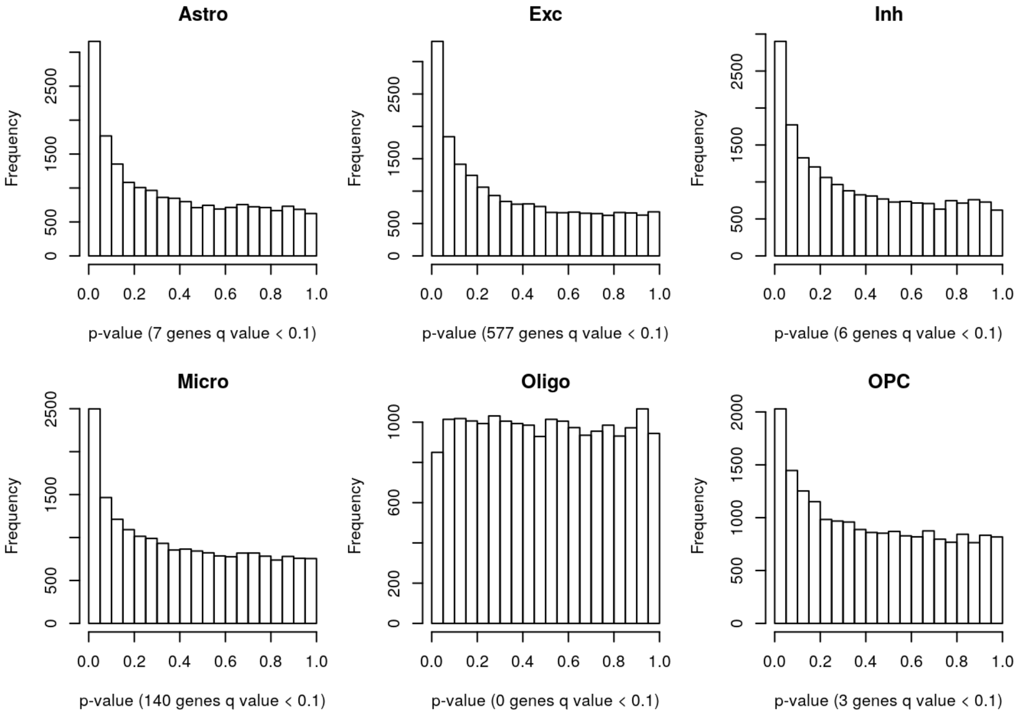

Figure S38: TOAST p-value distribution in the ASD study where the case-control label has been permuted to reflect the null distribution.

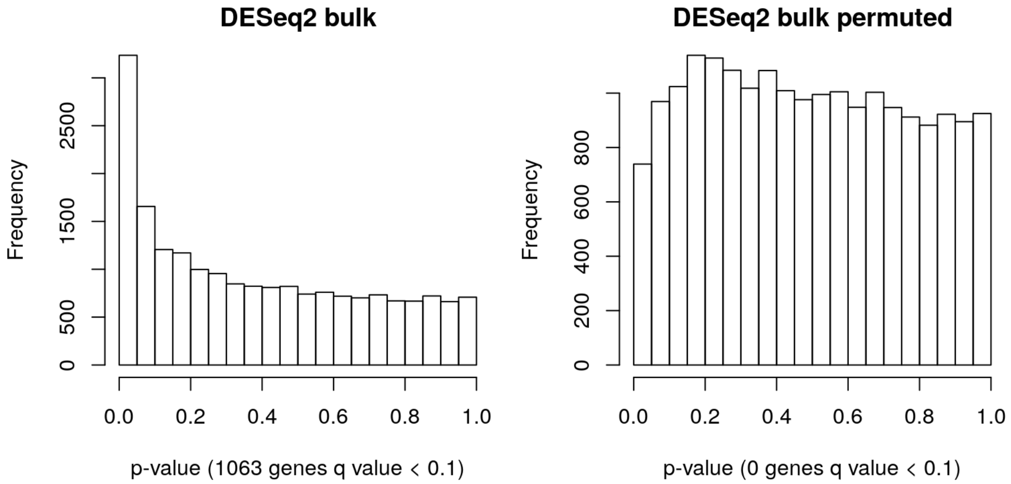

Figure S39: DESeq2 p-value distribution in the ASD study where the case-control label is either unpermuted or permuted.

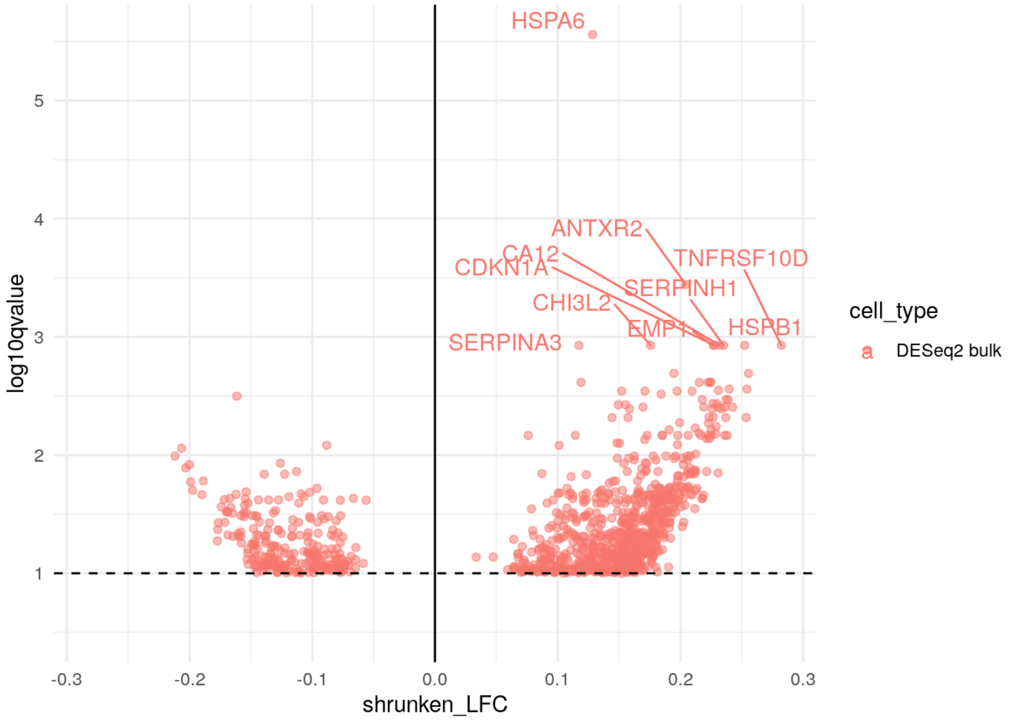

Figure S40: DESeq2 volcano plot in the ASD study.

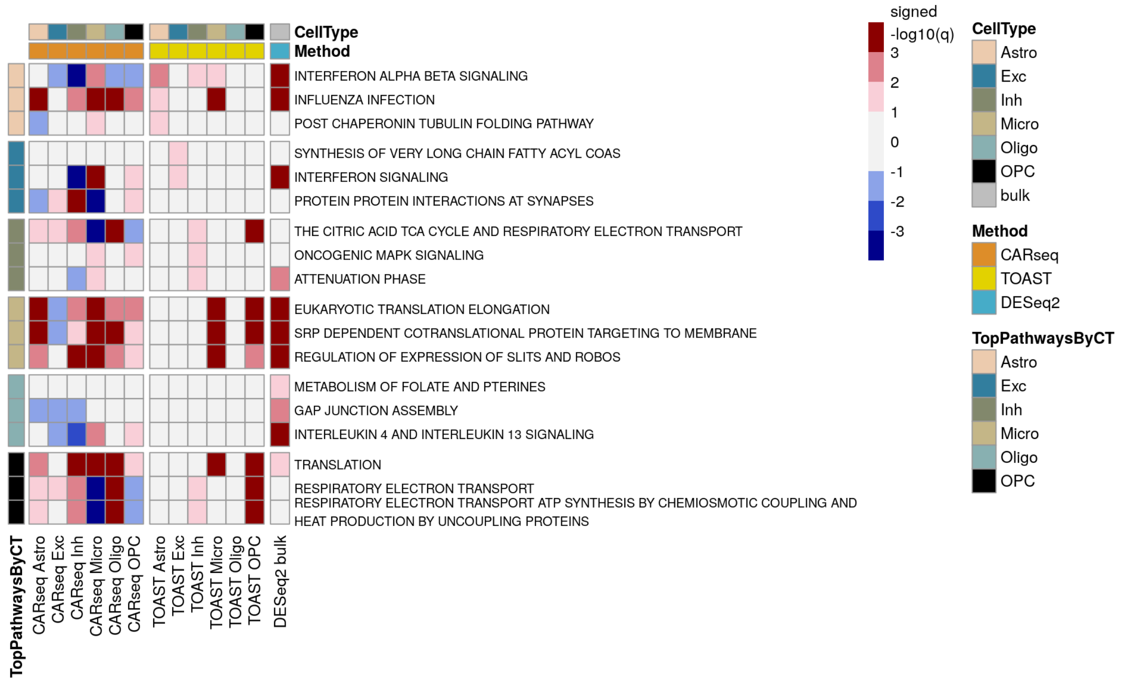

Figure S41: REACTOME GSEA ranked by TOAST in the ASD study.

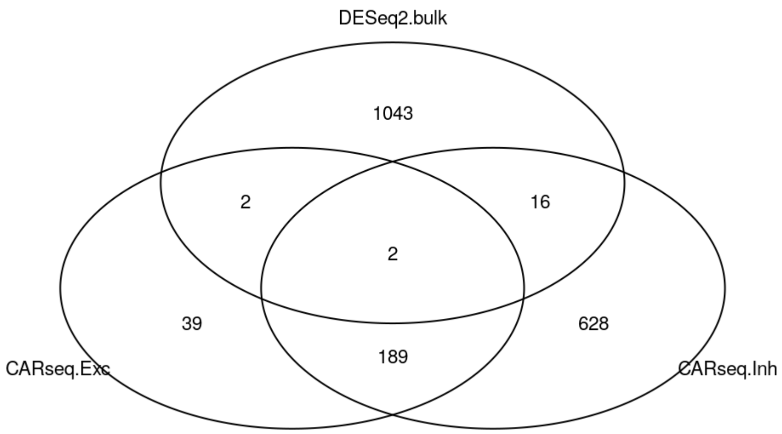

Figure S42: Venn plot of DEGs ($q$-value $< 0.1$) in the ASD study.

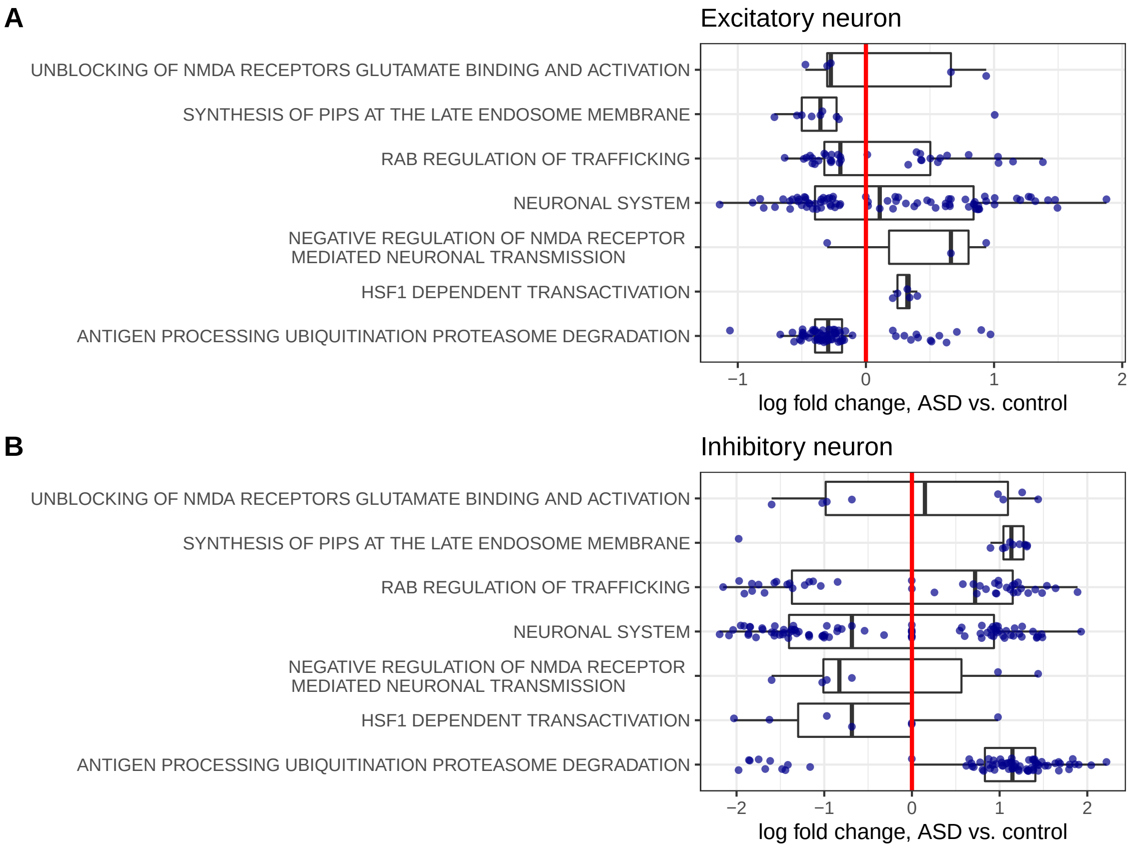

Figure S43: Shrunken log fold change estimates (ASD vs. controls) for genes belonging to some REACTOME pathways. Panel A includes all the pathways related with NDMA. Panels B and C include the pathways identified by GSEA in excitatory/inhibitory neurons, from either SCZ or ASD studies. Only the genes with CT-specific-DE p-value (comparing ASD vs. controls) smaller than 0.05 are shown.

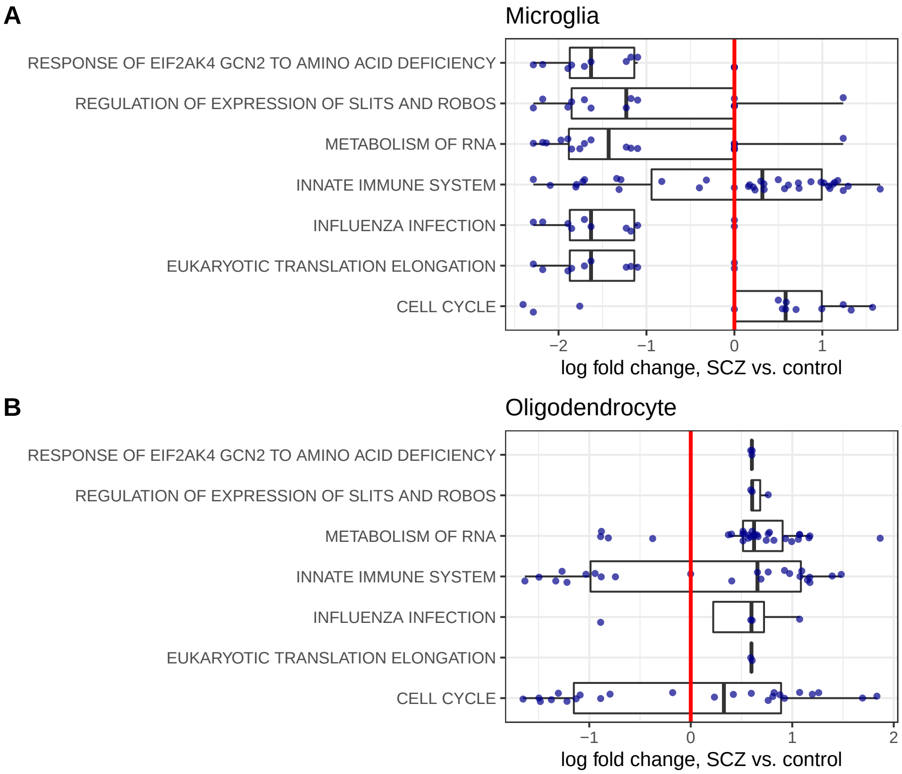

Figure S44: Shrunken log fold change estimates (ASD vs. controls) for genes belonging to the REACTOME pathways identified by GSEA in microglias or oligodendrocytes, from either SCZ or ASD studies. Only the genes with CT-specific-DE p-value (comparing ASD vs. controls) smaller than 0.05 are shown.

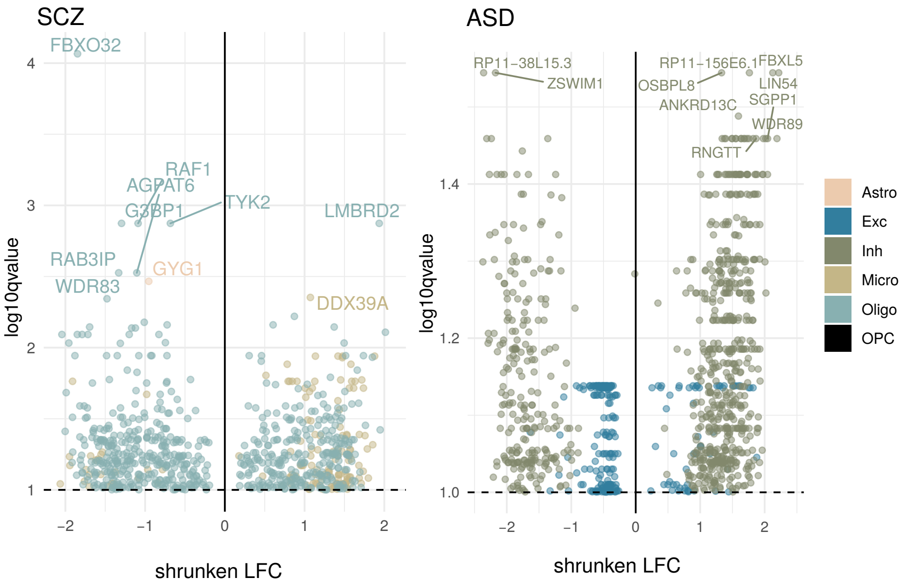

Figure S45: Volcano plot of -log10(q-value) vs. shrunken log fold change (LFC) for differentially expressed genes (DEGs) in SCZ (left panel) and ASD (right panel) inferred by CARseq. Only genes passing a q-value cutoff of 0.1 and an absolute value of LFC > 0.01 are shown. Top 10 CT-specific DEGs are labeled for each study.

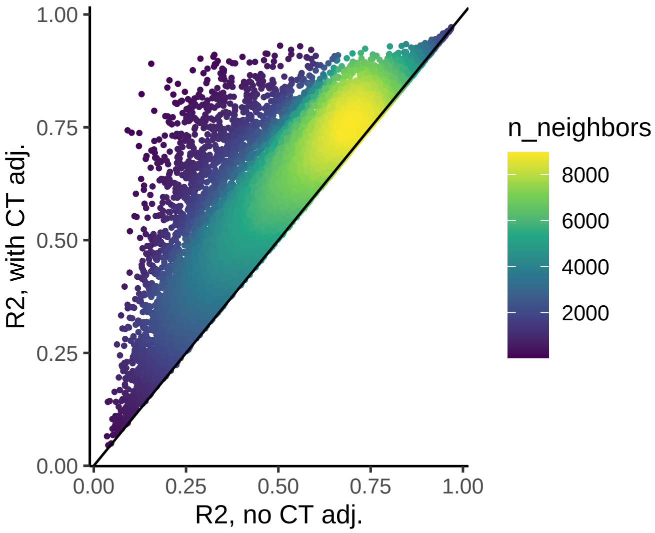

Figure S46: Compare the R2 of two linear models with log-transformed and read-depth corrected gene expression from SCZ and control subjects as response. The first model includes all the covariates used in the CAR-seq analysis. The second model includes all these covariates plus log-transformed cell type composition ratios for 5 cells types with excitatory neurons as reference.

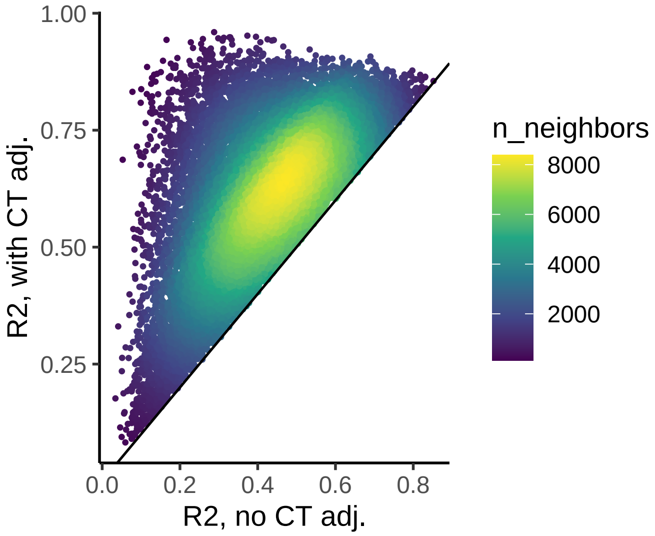

Figure S47: Compare the R2 of two linear models with log-transformed and read-depth corrected gene expression from ASD and control subjects as response. The first model includes all the covariates used in the CAR-seq analysis. The second model includes all these covariates plus log-transformed cell type composition ratios for 5 cells types with excitatory neurons as reference.

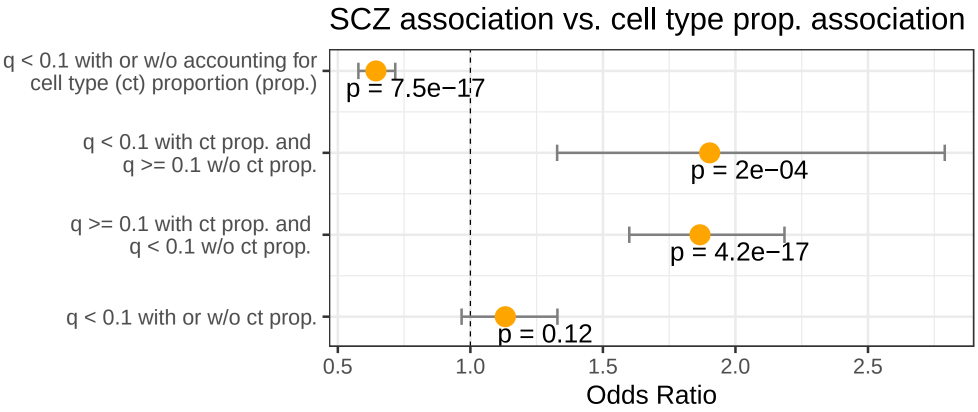

Figure S48: Odds ratio for four patterns of SCZ associations (four rows in the plot) vs. association with cell type composition. Here we consider a gene is associated with cell type composition if its p-value (ANOVA p-value by comparing the two models mentioned in Figure S46) is smaller than the Bonferroni threshold.

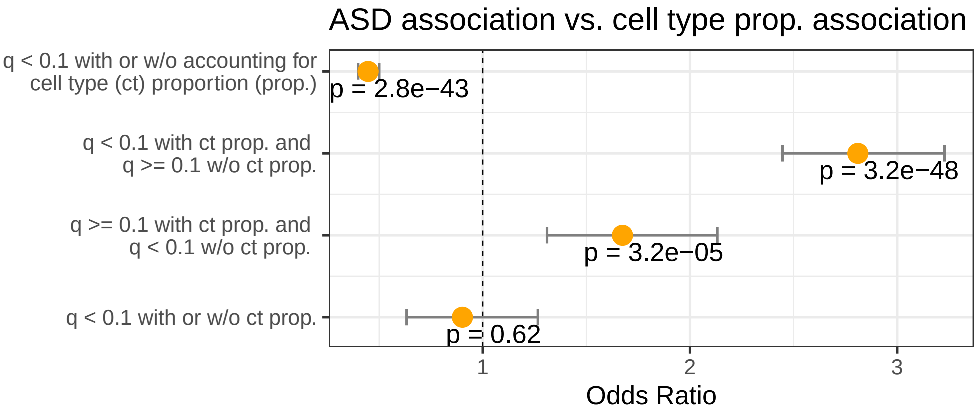

Figure S49: Odds ratio for four patterns of ASD associations (four rows in the plot) vs. association with cell type composition. Here we consider a gene is associated with cell type composition if its p-value (ANOVA p-value by comparing the two models mentioned in Figure S47) is smaller than the Bonferroni threshold.

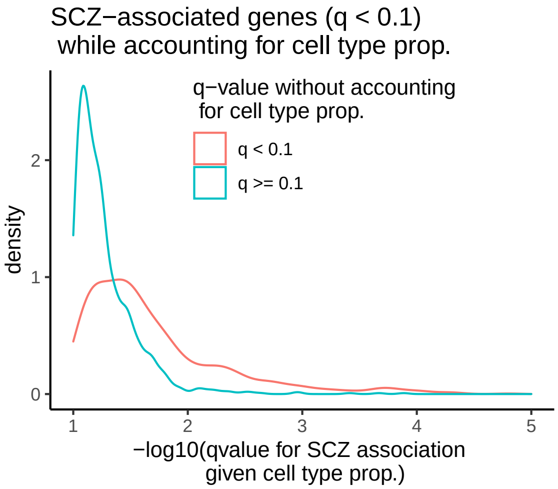

Figure S50: SCZ association strength for two groups of SCZ-associated genes: those with q-value $< 0.1$ or $\geq 0.1$ before accounting for cell type composition.

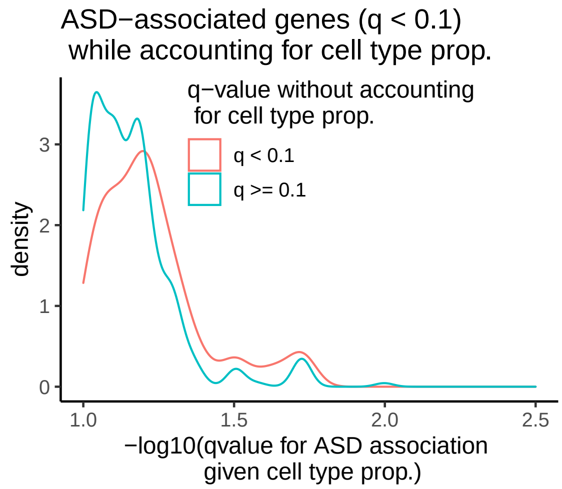

Figure S51: ASD association strength for two groups of ASD-associated genes: those with q-value $< 0.1$ or $\geq 0.1$ before accounting for cell type composition.

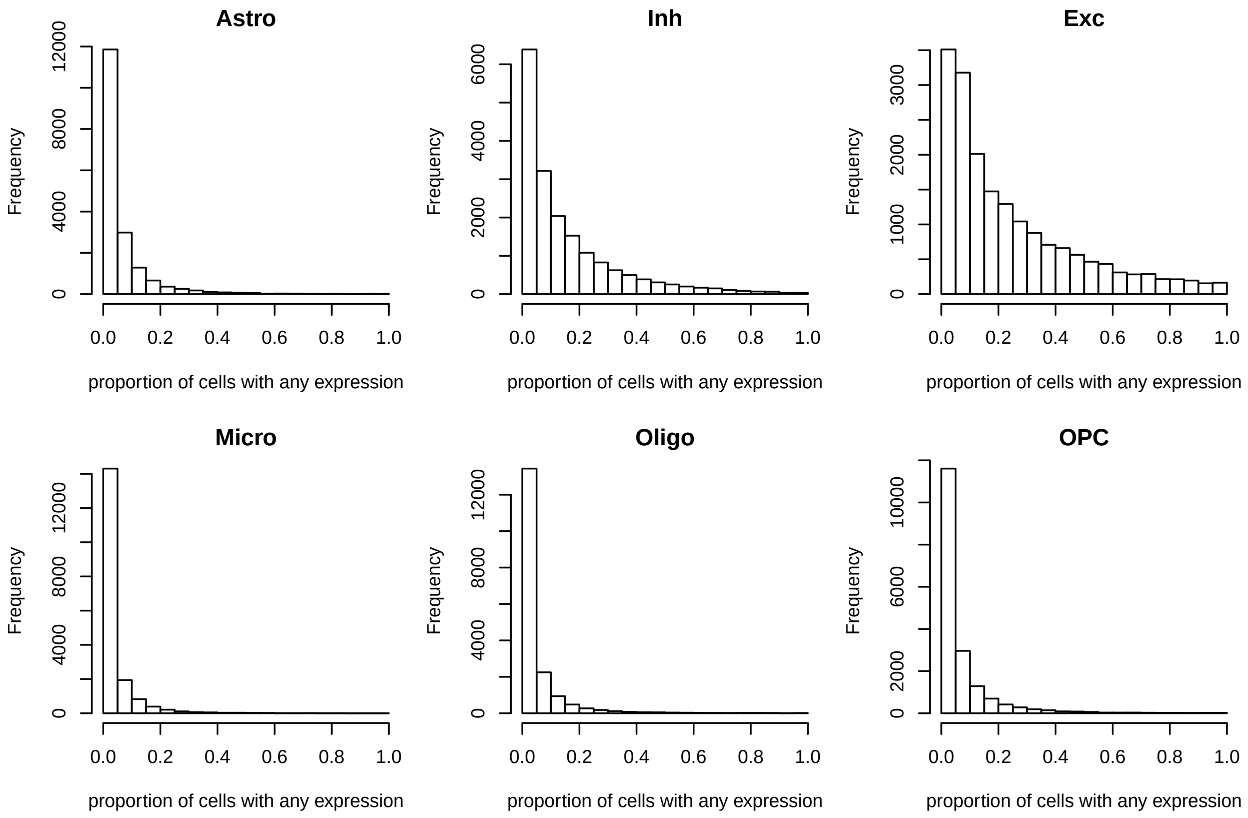

Figure S52: The distribution of the proportion of cells in which each of the 18,041 genes is expressed.

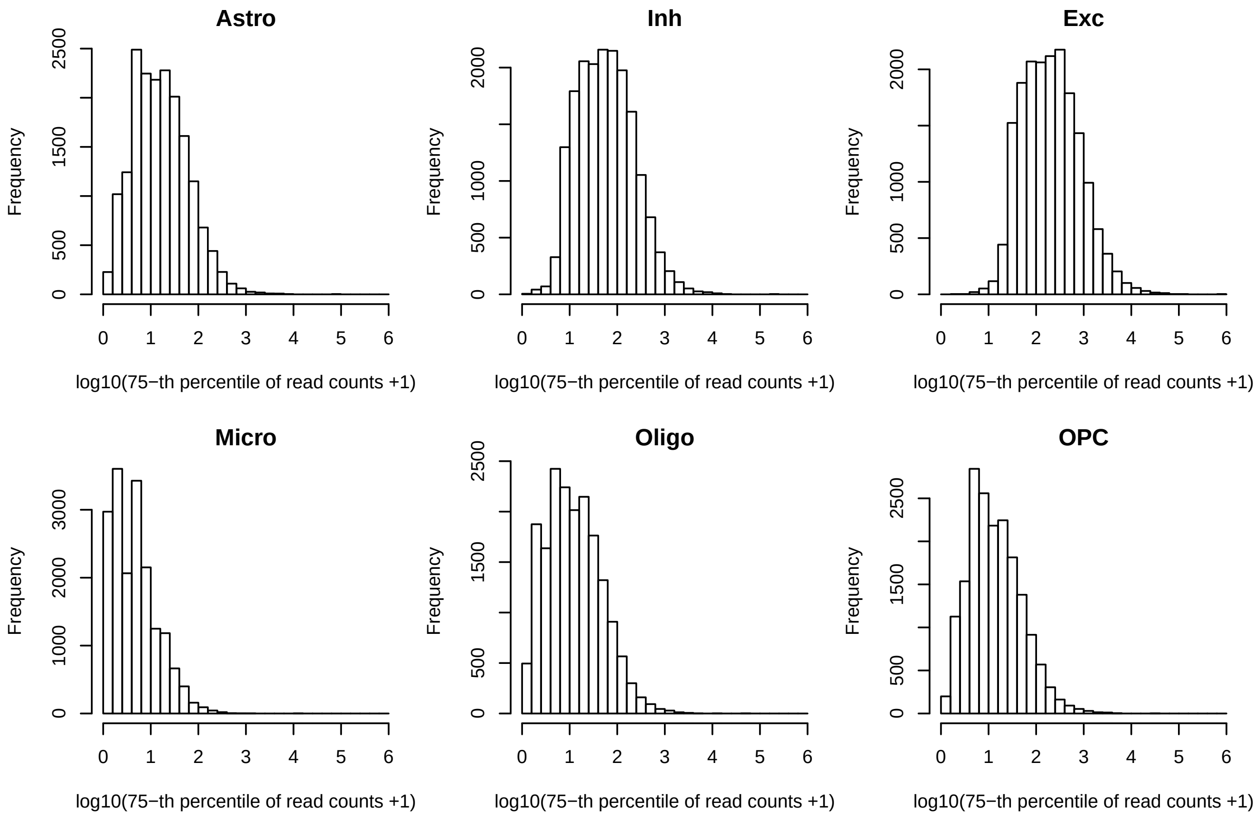

Figure S53: The 75 percentile of gene expression for each gene in pseudo-bulk RNA-seq data that is created by adding up the read counts across all the cells of the same cell type for each individual.

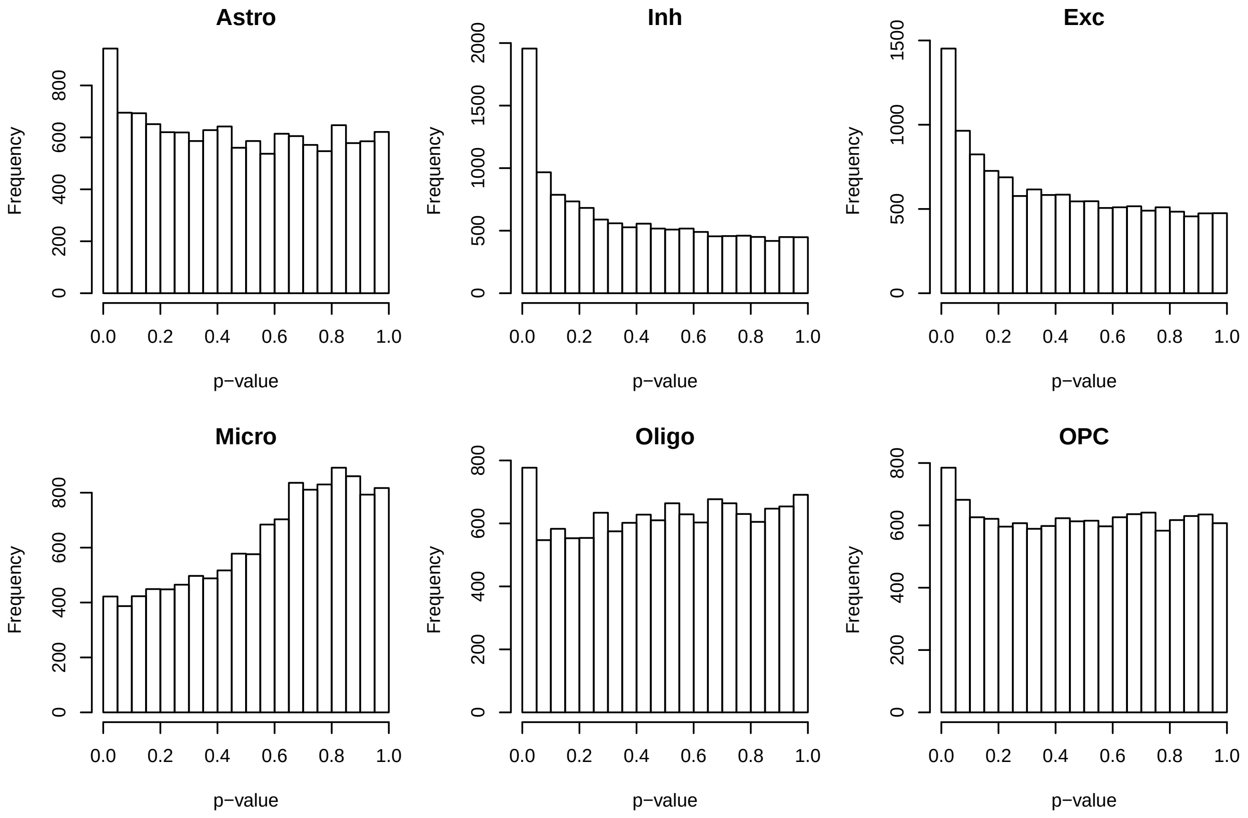

Figure S54: The distribution of DESeq2 p-values to assess DE for each cell type using snRNA-seq data from Velmeshev et al. (2019).

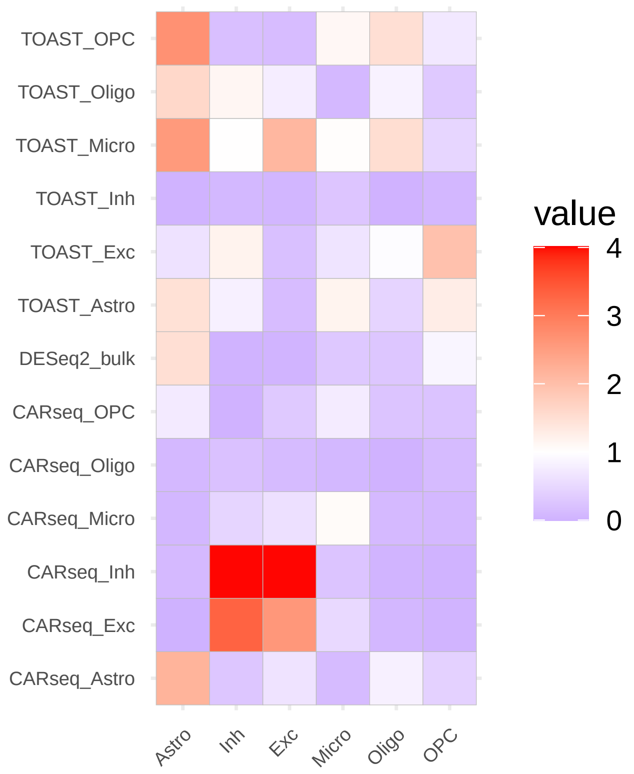

Figure S55: The -log10(p-value) from Fisher's exact test comparing the the overlap of the DE genes between results from bulk RNA-seq data (rows) vs. the results from snRNA-seq analysis. A liberal p-value cutoff 0.05 is used to ensure there are enough DE genes for comparison.

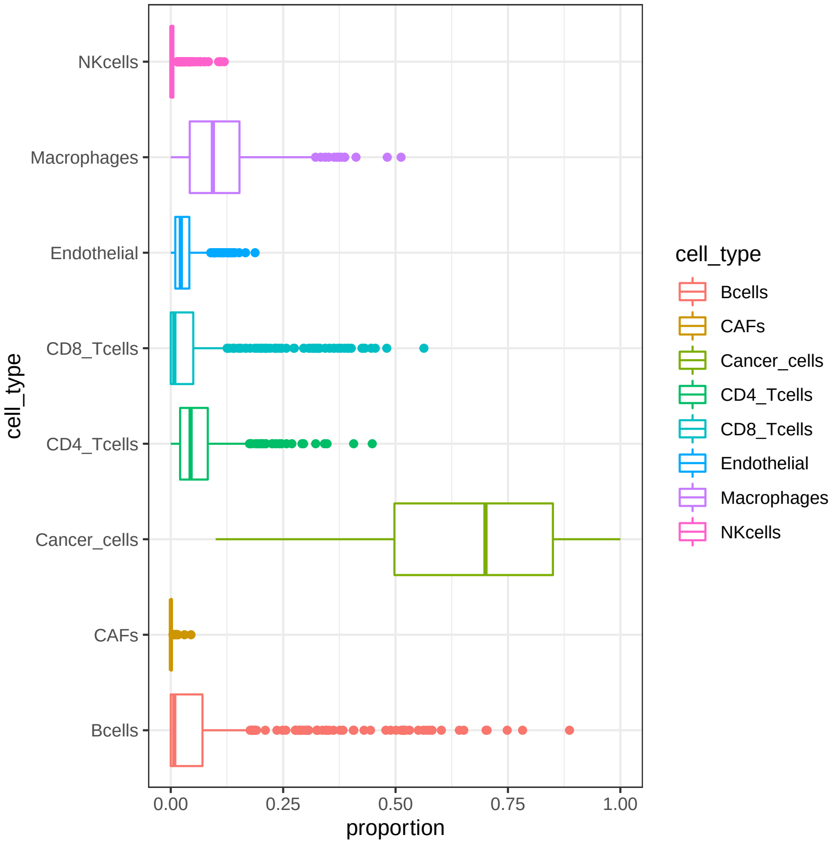

Figure S56: Cell type fraction estimates based on CIBERSOTx, combined with tumor purity estimates.

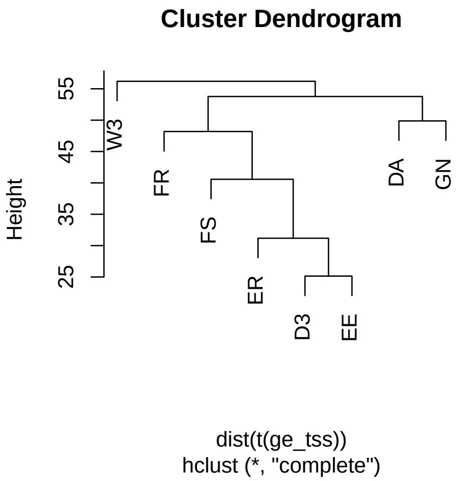

Figure S57: Clustering of average gene expression profiles of different tissue source sites.

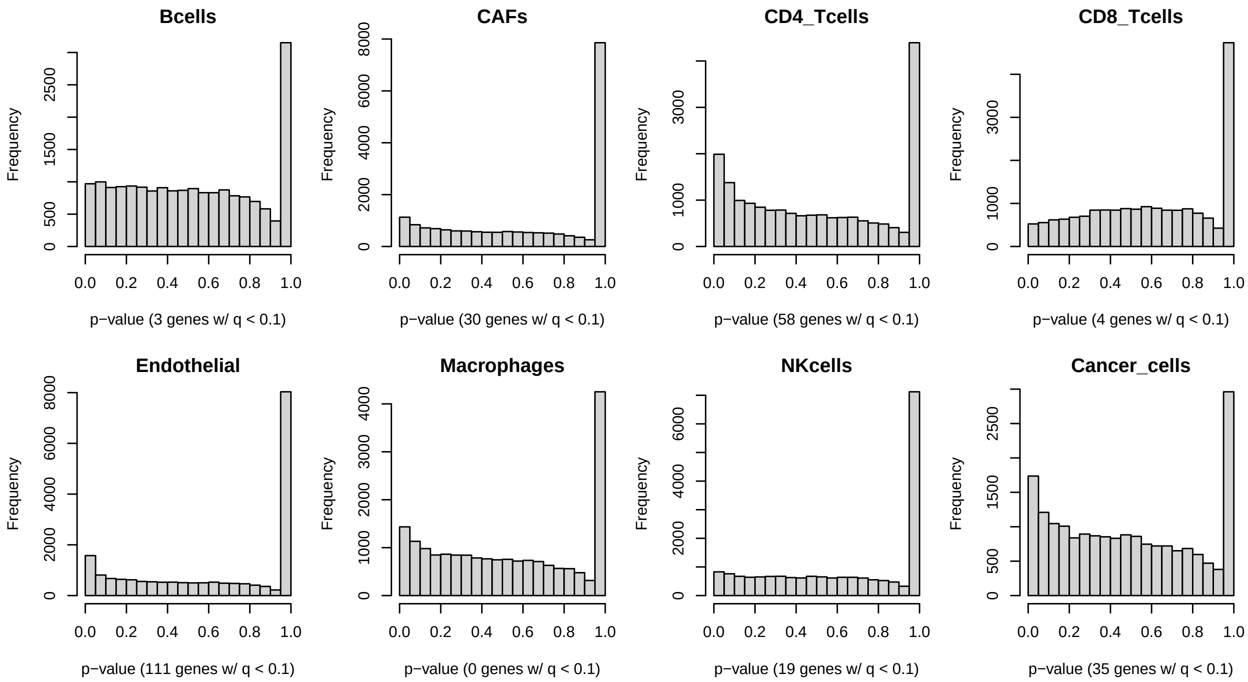

Figure S58: The distribution of CARseq p-values for TCGA SKCM analysis, where we assess the association versus an indicator of 5 year survival.

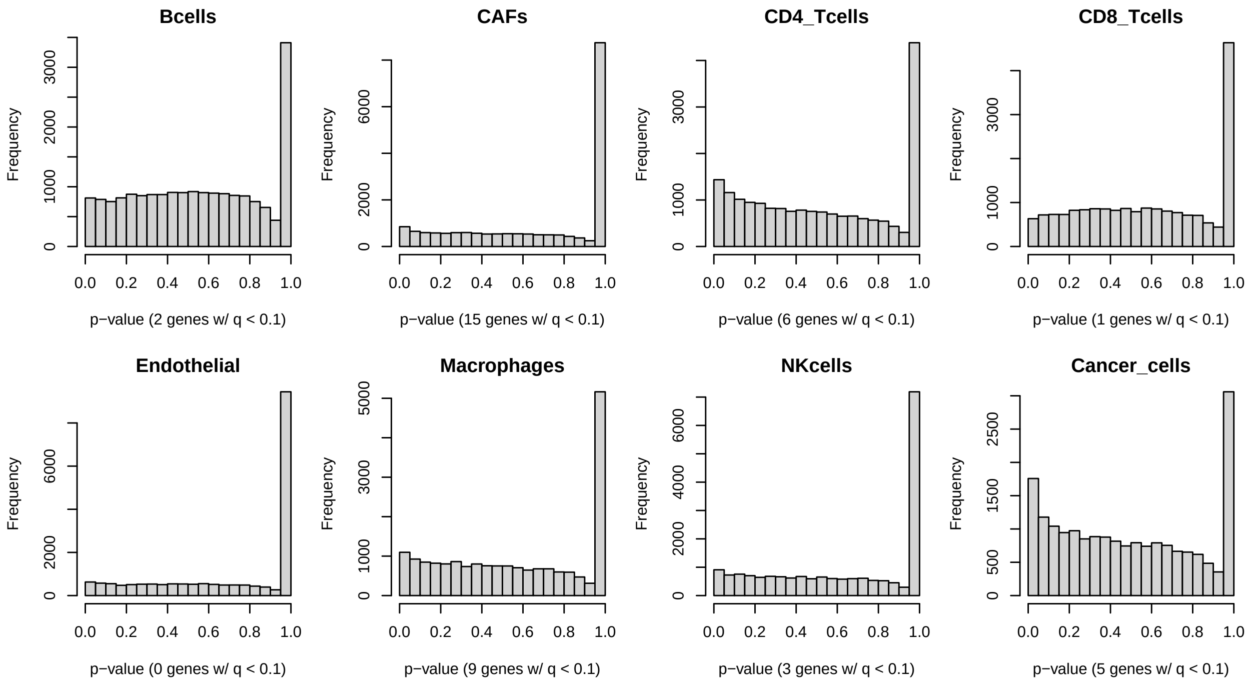

Figure S59: The distribution of CARseq p-values for TCGA SKCM analysis, where we assess the association versus a **permuted** indicator of 5 year survival.

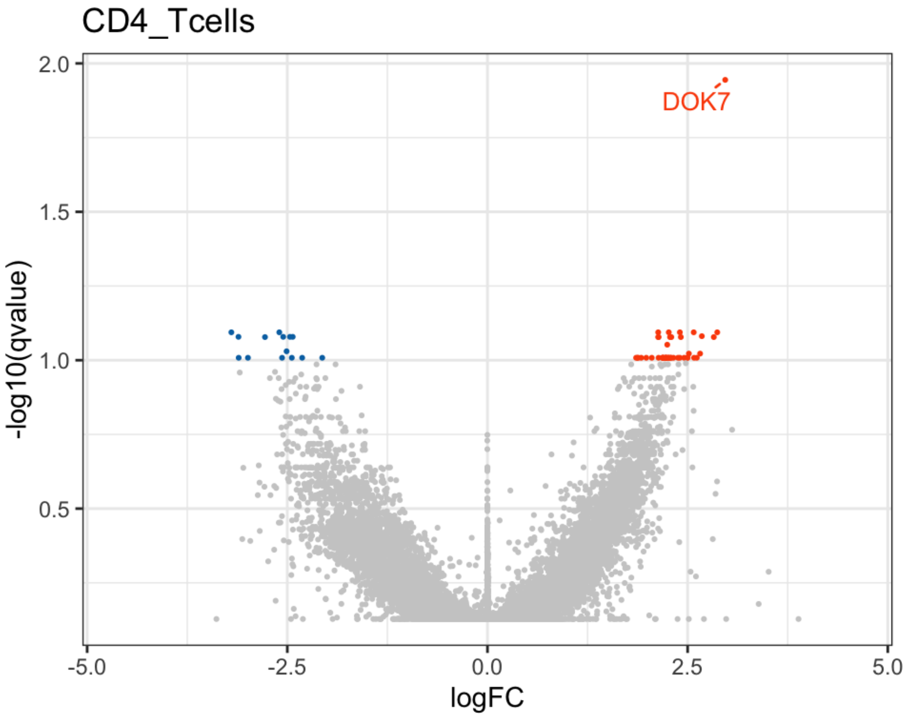

Figure S60: Valcano plot for CARseq results of CD4T cells.

New neurons use slit-robo signaling to migrate through the glial meshwork and approach a lesion for functional regeneration. *Science advances*, **4**(12):eaav0618.

[Li et al., 2019] Li, Z., Wu, Z., Jin, P., and Wu, H., 2019. Dissecting differential signals in high-throughput data from complex tissues. *Bioinformatics*, **35**(20):3898–3905.

[Love et al., 2014] Love, M. I., Huber, W., and Anders, S., 2014. Moderated estimation of fold change and dispersion for RNA-seq data with DESeq2. *Genome Biology*, **15**:550.

[Lun et al., 2016] Lun, A. T., McCarthy, D. J., and Marioni, J. C., 2016. A step-by-step workflow for low-level analysis of single-cell rna-seq data with bioconductor. *F1000Research*, **5**.

[McCartney et al., 2014] McCartney, A. J., Zolov, S. N., Kauffman, E. J., Zhang, Y., Strunk, B. S., Weisman, L. S., and Sutton, M. A., 2014. Activity-dependent pi (3, 5) p2 synthesis controls ampa receptor trafficking during synaptic depression. *Proceedings of the National Academy of Sciences*, **111**(45):E4896–E4905.

[Molenberghs and Verbeke, 2007] Molenberghs, G. and Verbeke, G., 2007. Likelihood Ratio, Score, and Wald Tests in a Constrained Parameter Space. *The American Statistician*, **61**(1):22–27.

[Newman et al., 2015] Newman, A. M., Liu, C. L., Green, M. R., Gentles, A. J., Feng, W., Xu, Y., Hoang, C. D., Diehn, M., and Alizadeh, A. A., 2015. Robust enumeration of cell subsets from tissue expression profiles. *Nature Methods*, **12**(5):453–457.

[Newman et al., 2019] Newman, A. M., Steen, C. B., Liu, C. L., Gentles, A. J., Chaudhuri, A. A., Scherer, F., Khodadoust, M. S., Esfahani, M. S., Luca, B. A., Steiner, D., *et al.*, 2019. Determining cell type abundance and expression from bulk tissues with digital cytometry. *Nature Biotechnology*, :1.

[Nocedal and Wright, 2006] Nocedal, J. and Wright, S. J., 2006. *Numerical optimization*. Springer series in operations research. Springer, New York, 2nd ed edition.

[Parikshak et al., 2016] Parikshak, N. N., Swarup, V., Belgard, T. G., Irimia, M., Ramaswami, G., Gandal, M. J., Hartl, C., Leppa, V., Ubieta, L. d. l. T., Huang, J., *et al.*, 2016. Genome-wide changes in lncRNA, splicing, and regional gene expression patterns in autism. *Nature*, **540**(7633):423–427.

[Park, 2006] Park, M. Y., 2006. Generalized linear models with regularization. *PhD Thesis*, **Stanford University, Department of Statistics**.

[Petrelli et al., 2016] Petrelli, F., Pucci, L., and Bezzi, P., 2016. Astrocytes and microglia and their potential link with autism spectrum disorders. *Frontiers in cellular neuroscience*, **10**:21.

[Prata et al., 2017] Prata, J., Santos, S. G., Almeida, M. I., Coelho, R., and Barbosa, M. A., 2017. Bridging autism spectrum disorders and schizophrenia through inflammation and biomarkers-pre-clinical and clinical investigations. *Journal of neuroinflammation*, **14**(1):179.

[Racle et al., 2017a] Racle, J., de Jonge, K., Baumgaertner, P., Speiser, D. E., and Gfeller, D., 2017a. Simultaneous enumeration of cancer and immune cell types from bulk tumor gene expression data. *elife*, **6**:e26476.

[Racle et al., 2017b] Racle, J., Jonge, K. d., Baumgaertner, P., Speiser, D. E., and Gfeller, D., 2017b. Simultaneous enumeration of cancer and immune cell types from bulk tumor gene expression data. *eLife*, **6**:e26476.

[Robinson et al., 2010] Robinson, M. D., McCarthy, D. J., and Smyth, G. K., 2010. edgeR: a Bioconductor package for differential expression analysis of digital gene expression data. *Bioinformatics*, **26**(1):139–140.

[Robinson and Smyth, 2008] Robinson, M. D. and Smyth, G. K., 2008. Small-sample estimation of negative binomial dispersion, with applications to SAGE data. *Biostatistics*, **9**(2):321–332.

[Self and Liang, 1987] Self, S. G. and Liang, K.-Y., 1987. Asymptotic Properties of Maximum Likelihood Estimators and Likelihood Ratio Tests under Nonstandard Conditions. *Journal of the American Statistical Association*, **82**(398):605–610.

[Stark and L. Parker, 1995] Stark, P. and L. Parker, R., 1995. Bounded-Variable Least-Squares: an Algorithm and Applications. *Computational Statistics*, **10**.

[The Brainstorm Consortium et al., 2018] The Brainstorm Consortium, Anttila, V., Bulik-Sullivan, B., Finucane, H. K., Walters, R. K., Bras, J., Duncan, L., Escott-Price, V., Falcone, G. J., Gormley, P., *et al.*, 2018. Analysis of shared heritability in common disorders of the brain. *Science*, **360**(6395).

[Tirosh et al., 2016] Tirosh, I., Izar, B., Prakadan, S. M., Wadsworth, M. H., Treacy, D., Trombetta, J. J., Rotem, A., Rodman, C., Lian, C., Murphy, G., *et al.*, 2016. Dissecting the multicellular ecosystem of metastatic melanoma by single-cell rna-seq. *Science*, **352**(6282):189–196.

[Velmeshev et al., 2019] Velmeshev, D., Schirmer, L., Jung, D., Haeussler, M., Perez, Y., Mayer, S., Bhaduri, A., Goyal, N., Rowitch, D. H., and Kriegstein, A. R., *et al.*, 2019. Single-cell genomics identifies cell type–specific molecular changes in autism. *Science*, **364**(6441):685–689.

[Wilson et al., 2019] Wilson, D. R., Jin, C., Ibrahim, J. G., and Sun, W., 2019. ICeD-T Provides Accurate Estimates of Immune Cell Abundance in Tumor Samples by Allowing for Aberrant Gene Expression Patterns. *Journal of the American Statistical Association*, **0**(0):1–11.

[Zaitsev et al., 2019] Zaitsev, K., Bambouskova, M., Swain, A., and Artyomov, M. N., 2019. Complete deconvolution of cellular mixtures based on linearity of transcriptional signatures. *Nature Communications*, **10**(1):2209.

[Zhong and Liu, 2012] Zhong, Y. and Liu, Z., 2012. Gene expression deconvolution in linear space. *Nature Methods*, **9**(1):8–9.

[Zhou et al., 2011] Zhou, Y.-H., Xia, K., and Wright, F. A., 2011. A powerful and flexible approach to the analysis of RNA sequence count data. *Bioinformatics*, **27**(19):2672–2678.
